# Single-cell multiome and spatial profiling reveals pancreas cell type-specific gene regulatory programs driving type 1 diabetes progression

**DOI:** 10.1101/2025.02.13.637721

**Authors:** Rebecca Melton, Sara Jimenez, Weston Elison, Luca Tucciarone, Abigail Howell, Gaowei Wang, Denise Berti, Elisha Beebe, Michael Miller, Chun Zeng, Carolyn McGrail, Kennedy VanderStel, Katha Korgaonkar, Ruth Elgamal, Hannah Mummey, Joshua Chiou, Emily Griffin, Irina Kusmartseva, Mark Atkinson, Sebastian Preissl, Fabian J. Theis, Maike Sander, Kyle J. Gaulton

**Affiliations:** Biomedical sciences program, University of California San Diego, La Jolla CA; Department of Computational Health, Institute of Computational Biology, Helmholtz, Munich, Germany; Department of Pediatrics, University of California San Diego, La Jolla CA; Center for Epigenomics, University of California San Diego, La Jolla CA; Bioinformatics and systems biology program, University of California San Diego, La Jolla CA; Pfizer Research and Discovery, Pfizer Inc, Cambridge, MA; Department of Pathology, Immunology and Laboratory Medicine, University of Florida, Gainesville FL; Department of Cellular & Molecular Medicine, University of California San Diego, La Jolla CA; Institute of Experimental and Clinical Pharmacology and Toxicology, Faculty of Medicine, University of Freiburg, Freiburg, Germany; Max Delbrück Center for Molecular Medicine, Berlin, Germany

## Abstract

Cell type-specific regulatory programs that drive type 1 diabetes (T1D) in the pancreas are poorly understood. Here we performed single nucleus multiomics and spatial transcriptomics in up to 32 non-diabetic (ND), autoantibody-positive (AAB+), and T1D pancreas donors. Genomic profiles from 853,005 cells mapped to 12 pancreatic cell types, including multiple exocrine sub- types. Beta, acinar, and other cell types, and related cellular niches, had altered abundance and gene activity in T1D progression, including distinct pathways altered in AAB+ compared to T1D. We identified epigenomic drivers of gene activity in T1D and AAB+ which, combined with genetic association, revealed causal pathways of T1D risk including antigen presentation in beta cells. Finally, single cell and spatial profiles together revealed widespread changes in cell-cell signaling in T1D including signals affecting beta cell regulation. Overall, these results revealed drivers of T1D progression in the pancreas, which form the basis for therapeutic targets for disease prevention.

## Introduction

Type 1 diabetes (T1D) is a complex endocrine disorder characterized by autoimmune destruction of beta cells in the pancreatic islets, leading to lifelong dependence on insulin therapy. The destruction of beta cells in T1D is caused by interactions with multiple cell types in and surrounding the islet microenvironment including infiltrating immune cells, other endocrine cells, and endothelial cells^1–3^. Cell types in the pancreas outside the local islet environment, such as exocrine acinar and ductal cells, are also increasingly implicated in T1D pathogenesis^4,5^. Beta cells themselves likely contribute to the development of T1D as well through response to environmental factors, external signaling to immune, beta, and other cell types, and cellular survival^6^. The sequence of events in the pancreas that drives initiation of beta cell autoimmunity and progression through stages of T1D, however, as well as the role of each pancreatic cell type in these processes, remains poorly understood.

Seroconversion to autoantibody positivity (AAB+) against islet proteins (i.e. self-antigens) precedes T1D onset in nearly all cases and is used as a clinical biomarker of T1D progression^7,8^. Individuals at T1D diagnosis can present with differing number and type of autoantibodies, which are associated with varying rates of disease incidence; for example, the presence of a single islet AAB has a relatively low lifetime risk of T1D while individuals with multiple AAB have disease rates over 90%^9–11^. As clinical presentation of T1D does not occur until a large fraction of beta cells has been destroyed, there is a window of time between seroconversion and T1D onset where disease processes can potentially be halted or reversed^7^. Even after onset of T1D, residual beta cell mass could potentially be modulated therapeutically to restore metabolic function^12^. Defining changes in disease-relevant cell types across the stages of T1D progression would both improve our understanding of the mechanisms of T1D as well as reveal potential targets to prevent or reverse disease. Furthermore, an improved understanding of key changes associated with progression would also help identify novel biomarkers of T1D, which are particularly needed in the early stages of disease to identify progressors and candidates for therapeutic intervention^13^.

Single cell technology, the focus of this work, enables profiling of individual cells within the pancreas^5,14^. Previous single cell studies of the pancreas in T1D have been limited in that they focused primarily on gene expression profiling of dispersed cells^4,15^, which does not provide information on the spatial localization of cellular transcriptomes within the pancreas nor the genomic elements driving changes in gene activity. Recent developments in spatial transcriptomics enables profiling cells in their native location^16^, which enables understanding cell type-specific changes in the context of specific cellular neighborhoods and niches in the pancreas. This is particularly important in the context of T1D which has extensive heterogeneity in disease processes within the pancreas^17^. In addition, single cell epigenome profiling, for example using snATAC-seq or single cell multiome (paired snRNA-seq+snATAC-seq), can reveal transcriptional regulators, *cis*-regulatory elements (cREs), and gene regulatory networks driving altered gene expression in T1D^5,14^. Critically, gene regulatory networks and cREs can be intersected with T1D-associated variation to infer cell type-specific regulatory programs that may play a causal role in driving disease^18,19^.

Previous single cell studies have also been limited in the extent to which they have captured key windows of T1D progression and pathogenesis^4,15^. Specifically, non-diabetic AAB+ donors in these efforts were largely those with single glutamic acid decarboxylase (GAD) autoantibodies^4^, which have a relatively lower risk of developing T1D compared to multiple AAB+ donors and do not reflect the full arc of progression to T1D^20^. Furthermore, many of the T1D donors in these studies had long-standing T1D where disease processes are potentially more dormant, whereas profiling donors who had more recently developed T1D may give greater insight into active disease processes. Third, as these studies profiled purified islets, they offer more restricted insight into genomic changes in cells outside of the islet microenvironment during T1D progression, including in exocrine cells which are both altered in T1D as well as implicated causally in the development of T1D^4,18,21^.

In this study, we addressed these limitations by performing single cell gene expression and epigenome profiling in whole pancreas from 32 non-diabetic, non-diabetic single and multiple AAB+, recent-onset T1D, and long-standing T1D organ donors, as well as spatial transcriptomics in a subset of non-diabetic and recent-onset T1D donors. We determined changes in pancreatic cell type abundance, cellular pathways, gene regulatory networks, and cell-cell signaling across these stages of T1D progression and pathogenesis and, using T1D association data, identified pathways and gene networks that may play a causal role in the development of T1D.

## Results

### A comprehensive, multimodal, spatially resolved map of pancreatic cell types

We selected whole pancreas samples from 32 donors in the nPOD biorepository including 11 non-diabetic (ND), 9 non-diabetic autoantibody positive (ND AAB+), and 12 T1D which we separated into 7 recent onset (<1 year from diagnosis) and 5 longer duration (>5 years from diagnosis) (**Supplementary Table 1**). Within the ND AAB+ group, most organ donors, by our study design, had multiple autoantibodies (multiple ND AAB+). For all samples, we performed single nucleus RNA-seq (snRNA-seq) and single nucleus ATAC-seq (snATAC-seq) assays and, for eight of the samples, we performed single nucleus multiome (joint snRNA-seq and snATAC-seq in the same nucleus) assays. In addition, for six of the samples, we performed spatial transcriptomics assays using the CosMx Spatial Molecular Imager (**Figure 1A**).

**Figure 1.**
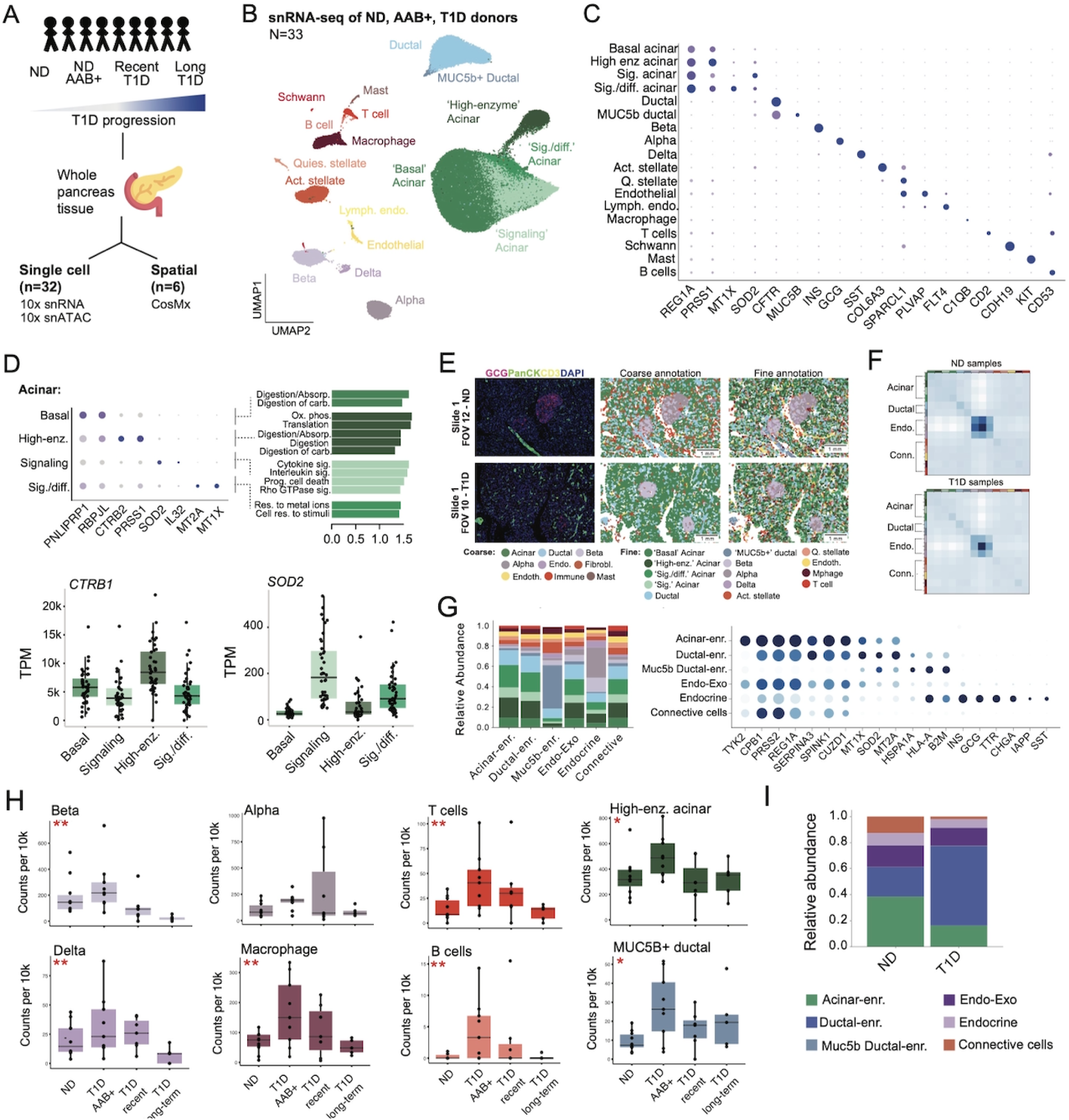
Cell type-specific map of gene expression in the pancreas. (A) Design of study profiling human pancreas from ND, ND AAB+ and T1D donors using single cell assays. (B) Uniform manifold approximation and projection (UMAP) plot showing clustering of 276,906 nuclei from single nuclear RNA-seq of 32 whole pancreas donors from the nPOD biorepository. Clusters are labeled based on cell type and sub-type annotations. (C) Dot plot showing the normalized expression levels of selected known marker genes for pancreatic cell types and sub- types. (D) Dot plot of genes with preferential expression across different sub-types of acinar cells (top left), and normalized enrichment score (NES) of pathways enriched in each subtype using fGSEA (top right). Donor transcripts per million (TPM) of selected genes with preferential expression in different sub-types of acinar cells. (E) Representative FOV per condition (ND: top, T1D: bottom) showing (from left to right) immunofluorescence, coarse cell type annotation with the spatial gene panel directly, and finer-grained cell type annotation transferred from the snRNA-seq data. (F) Matrix plots showing the neighborhood enrichment of cell types based on spatial neighbors. (G) Stacked barplot illustrating the relative abundance of each cell type in each multicellular niche (left). Dot plot showing the normalized gene expression levels of spatially variable genes across multi-cellular pancreatic niches (right). (H) Normalized cell counts for selected pancreatic cell types and sub-types organized by donor T1D and ND AAB+ status. ** FDR<.10, * uncorrected p<.05. (I) Stacked barplot showing the relative abundance of each multi-cellular niche per condition. Niches with * have altered abundance in ND samples (p<0.05).

After extensive barcode quality control and filtering steps, (**see Methods**), we performed integration using Harmony^22^ and clustered 276,906 gene expression profiles (**Figure 1B**, **Supplementary Figures 1-2**). We annotated the resulting 18 clusters based on the expression of known cell type marker genes which revealed 12 pancreatic cell types including exocrine (acinar, ductal), endocrine (alpha, beta, delta), immune (T cell, B cell, macrophage, mast), stellate, endothelial, and Schwann cells (**Figure 1B-C**, **Supplementary Table 2**). Cell type clusters had broadly consistent representation across donors and donor characteristics (**Supplementary Figures 2-3**). We aggregated expression profiles for all cells in the cell type and derived normalized expression levels of each gene using transcripts-per-million (TPM), which revealed, on average, 17,885 genes expressed (TPM>1) per cell type (**Supplementary Data 1**). For each cell type, we further identified genes with expression levels specific (FDR<.10) to the cell type which revealed both known as well as previously unreported sets of genes with cell type-specific expression (**Supplementary Table 3**); for example, well-known genes with expression highly specific to beta cells included *INS, IAPP* and *G6PC2* as well as others with no currently known role in beta cell function (e.g., *PLCH2, NRG2, RBFOX3, MTUS2)*.

Several cell types displayed multiple sub-clusters representing both known cell sub-types, such as active and quiescent stellate cells, blood vessel (BVEC) and lymphatic endothelial (LEC) cells, and MUC5b+ ductal cells, as well as several potential sub-types of acinar cells (**Figure 1B,C**). As the genomic properties of these sub-types have not been completely described in previous efforts, we derived sets of marker genes for each sub-type (**see Methods, Supplementary Table 3**). For BVECs and LECs, in addition to reported marker genes *PLVAP* (BVECs) and *FLT4* (LECs), we observed specific up-regulation of genes in each sub-type such as *INHBB, BMP6, FCN3,* and *PCAT19* in BVECs and *EFNA5, COLEC12,* and *MYCT1* in LECs.

In *MUCB5*+-ductal cells, there was up-regulation of *ERN2, CYP2C18, MYO7B,* and *DMBT1* compared to the primary sub-type of ductal cells. For acinar cells, the primary cluster, which we annotated as ‘basal’ acinar cells, was enriched for genes and pathways involved in digestive enzyme production and secretion. Other clusters included ‘high-enzyme’ acinar cells with higher expression of enzymes such as chymotrypsin (*CTRB1/2),* trypsinogen (*PRSS1, PRSS2*), lipase (*PNLIP*), carboxyl ester lipase (*CEL*), chymotrypsin-like elastase (*CELA3A/B*) and increased oxidative phosphorylation and translation, ‘signaling’ acinar cells with increased signaling and stress-response activity, and ‘signaling/differentiation’ acinar cells with increased signaling, metallothionein (*MT1/MT2)*, and identity and differentiation genes (*REG1A/B, PTF1A*) (**Figure 1D**).

To next characterize the spatial organization of pancreatic cell types, we performed RNA *in situ* hybridization (ISH) of 1,010 genes with CosMx from a subset of donors including three ND and three recent-onset T1D (**Supplementary Table 1, Supplementary Table 4**). We imaged a total of 82.6M transcripts from 71 fields of view (FOV) in whole pancreas sections from three ND (32 FOV) and three recent-onset T1D donors (39 FOV) (**Supplementary Figure 4A**) and assigned transcripts to 392,248 cells overall (80 median genes, 200 median transcripts per cell), using the CosMx default segmentation. We first performed unsupervised clustering of cellular gene expression profiles, which revealed nine distinct clusters including exocrine, endocrine, endothelial, immune and mast cells (**Supplementary Figure 4B**). We next mapped finer-grained cell type annotations from the snRNA-seq atlas using moscot^23^ (**Supplementary Figure 4B,C**), which revealed 14 total cell types and sub-types that were confirmed based on marker gene expression (**Figure 1E**). Spatial neighborhood enrichment using squidpy^24^ revealed expected cell types clustering together including acinar sub-types, ductal sub-types, endocrine cells (beta, alpha, delta), and connective cells (e.g., endothelial, immune, stellate) (**Figure 1F**).

Next, we sought to determine whether spatial neighborhoods form recurrent niches across the pancreas, by defining niches involving a cell type using a gene-gene covariance matrix^25^ in a spatial neighborhood of 30 cells. We recovered six niches in total, characterized by cell type abundance (**see Methods**), including three exocrine (acinar-enriched, ductal-enriched, and *MUC5b* ductal-enriched) niches, one endocrine niche, one niche including both endocrine and exocrine cells (endo-exo), and one niche consisting of connective cells (**Figure 1G**). To characterize each niche, we identified spatially variable genes (Moran’s I >0.2, p<.05) that captured gene signatures specific to the niche (**Figure 1G**). In the acinar-enriched niche, marker genes from the ‘basal’ and ‘high-enzyme’ cell types showed strong spatial clustering (*PRSS2, REG1A*). In comparison, the ductal-enriched niche had more spatial association with ‘signaling’ and ‘signaling/differentiation’ acinar cells (*MT1X, SOD2, MT2A*). Interestingly, in the *MUC5b* ductal-enriched niche, spatially variable genes were strongly associated with immune interactions (*HSPA1A, HLA-A,* and *B2M*). In addition, the endocrine niche had highly distinct patterns which highlighted multiple endocrine-specific genes (e.g. *INS, GCG, SST,* and *IAPP*) (**Figure 1G**).

Finally, we determined whether there were changes in abundance of cell types and sub-types in T1D progression based on snRNA-seq data (**see Methods**). There was a significant decrease (likelihood ratio test [LRT], FDR<.10) in beta cells (**Figure 1H**, **Supplementary Table 5**) although we still observed residual beta cell proportion in T1D particularly in recent-onset (ND=1.5%, recent-onset T1D=0.93%). We also observed significant decrease (FDR<.10) in delta cells in T1D, and increased abundance of multiple immune populations in ND AAB+ and recent-onset T1D. There was also more nominal evidence (p<.05) for altered abundance of specific cell sub-types including ‘enzyme-producing’ acinar (p=.037) and MUC5b+ ductal cells (p=.049). We next asked whether there were corresponding changes in the abundance of specific niches in T1D in spatial profiles. First, we quantified the pairwise similarity between ND and T1D spatial graphs using Wasserstein distance^26^ (**Supplementary Figure 4D**), which revealed significant changes in the underlying structure of endocrine cells (alpha and beta) in T1D. We moreover observed significant changes in the abundance of the endocrine niche, as well as the *MUC5b+* ductal cell niche, in T1D (p<.05) (**Figure 1I**).

### Comprehensive map of pancreatic cell type accessible chromatin

To understand how the epigenome may drive changes in cell type-specific gene expression in T1D, we next created a matched map of accessible chromatin in pancreatic cell types. For 29 nPOD donors we performed snATAC-seq assays, and we also used the snATAC-seq profiles from single cell multiome assays of eight donors described above. After quality control, filtering and initial clustering steps (**see Methods**), we annotated cell type identity by label transfer of the gene expression map using Seurat^27^. After filtering nuclei with low transfer predictions (<0.5), there were 203,348 chromatin profiles mapping to the same cell types and sub-types (**Figure 2A**, **Supplementary Figures 5,6**). We estimated that label transfer was >97% accurate at the cell type level by comparing the predicted and actual identity of accessible chromatin profiles in single cell multiome data. We also confirmed that predicted cell types had accessible chromatin at the promoter regions of key marker genes (**Figure 2B**). The proportions of each cell type were highly correlated between expression and accessible chromatin maps (r=.98, P=1.7×10^-13^; **Supplementary Figure 7**).

**Figure 2.**
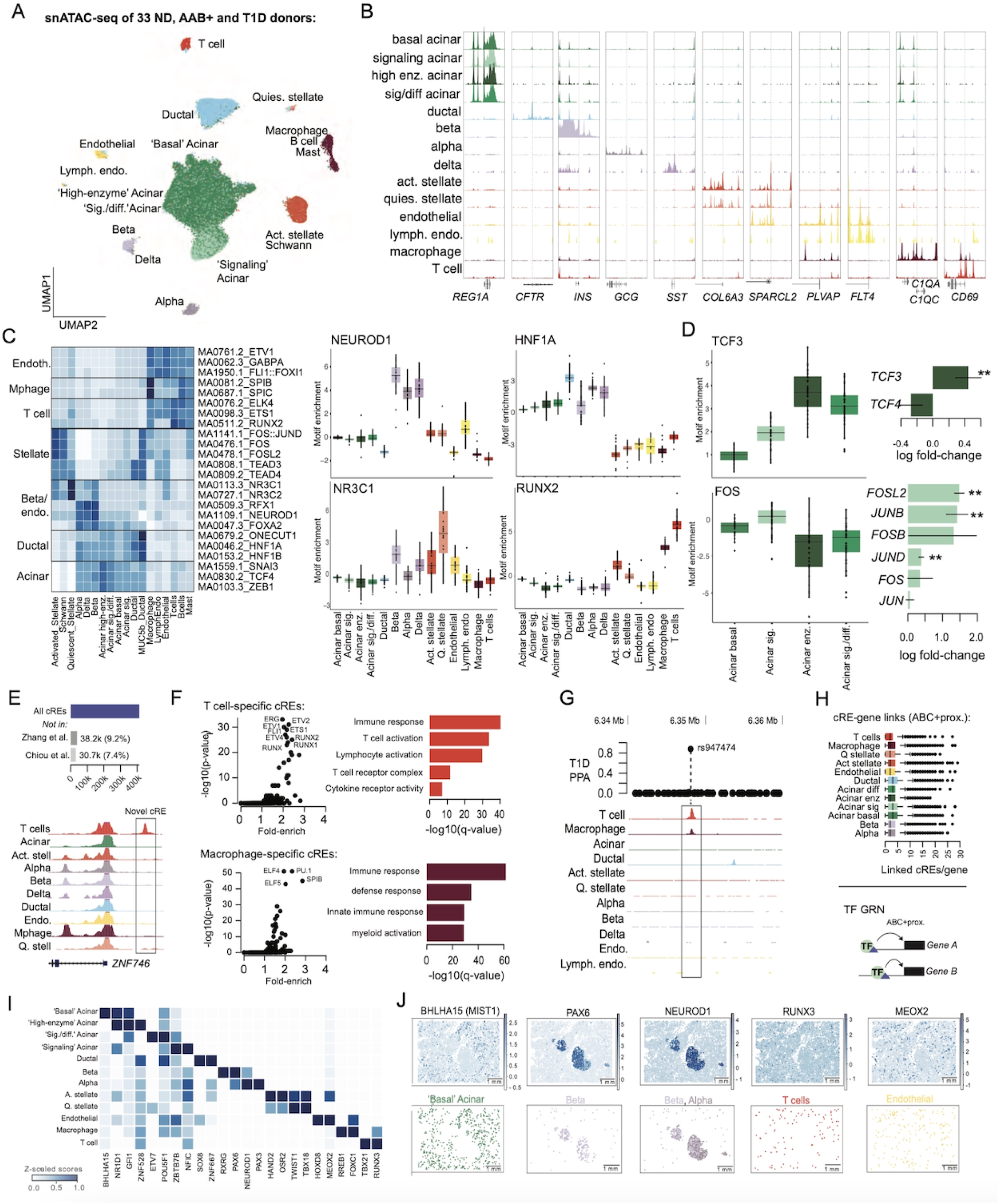
Cell type-specific map of accessible chromatin in the pancreas. (A) Uniform manifold approximation and projection (UMAP) plot showing clustering of 203,348 nuclei from single nuclear ATAC-seq of 29 whole pancreas donors from the nPOD biorepository. Clusters are labeled with cell type and sub-type identity based on label transfer from the gene expression map. (B) Genome browser showing cell type-specific accessible chromatin signal at the promoter regions of known marker genes for pancreatic cell types. (C) Heatmap showing genome-wide accessibility from chromVAR of sequence motifs for selected transcription factors (TF) across cell types (left), and boxplots showing donor-level accessibility of selected TF sequence motifs across cell types (right). (D) Genome-wide accessibility of sequence motifs for TFs with preferential enrichment in specific sub-types of acinar cells (left), and log fold-change in expression for genes in structural sub-families for the enriched TF motifs (right). (E) Number of cREs identified across all cell types and the percentage of cREs that do not overlap previous catalogs of cREs from Zhang et al and Chiou et al (top). Example of a pancreatic T cell-specific cRE novel to this study compared to previous catalogs at the *ZNF746* locus. (F) Sequence motifs for TFs enriched in cREs with activity specific to each cell type (left) and barplots showing -log10 p-values of gene sets enriched for proximity to cell type-specific cREs using GREAT. (G) Example of a cRE active in pancreatic T cells and macrophages that overlaps a candidate causal T1D risk variant at the *PRCKQ* locus. (H) Number of gene-CRE links per gene per cell type (top) and schematic of TF gene regulatory network (GRN) creation (bottom). (I) Matrix plot showing the scaled Z-score of TF activities for top TFs identified for each cell type using a t-test with overestimated variance. (J) Spatial plot of selected TFs showing the TF activity profile (top), and cell type distribution for the respective cell type (bottom).

We identified transcription factor (TF) binding motifs preferentially enriched in each pancreatic cell type and sub-type using chromVAR^28^. At the cell type level, enriched sequence motifs revealed key regulators of each cell type; for example, beta cells and other endocrine cells were enriched for RFX and FOXA motifs, ductal cells for HNF1, ONECUT and TEAD motifs, endothelial cells for ETV, FLI and GABPA motifs, and T cells for RUNX, ETV, and ETS motifs, among others (**Figure 2C**, **Supplementary Table 6**). Motif enrichments also highlighted regulators that distinguished accessible chromatin profiles of cell types within specific lineages; for example, NEUROD1 and NR3C1 had stronger enrichment in beta compared to other endocrine cells (**Figure 2C)**. Acinar cells showed distinct sets of enriched TF motifs across different sub-clusters, including ‘signaling’ acinar cells which were more enriched for FOS/JUN, ATF and NFE motifs (**Figure 2C,D**, **Supplementary Table 6)**. In ‘high-enzyme’ acinar cells, the strongest enrichments were for TFs such as ZEB, SNAI1-3, and TCF3-4, which were also the most enriched motifs in acinar cells overall compared to other cell types (**Figure 2C,D**, **Supplementary Table 6**). As structurally related TFs often have similar motifs, we linked TF motifs enriched in sub-clusters to specific TFs in the same structural sub-family with concordant expression patterns. For example, *FOSL2* and *JUNB/D*, as well as *ATF3*, *NFE2L2* and *BACH1/2,* were increased in ‘signaling’ acinar cells, and *TCF3* had increased expression in ‘high-enzyme’ acinar cells (**Figure 2D**).

For each cell type and sub-type, we next defined candidate *cis*-regulatory elements (cREs). We derived ‘pseudo’-bulk accessible chromatin profiles by aggregating reads from all cells for that cell type or sub-type and identified cREs by performing peak calling with MACS2. In total, there were 368,688 cREs across all cell types and an average of 94.3k cCREs per cell type (**Supplementary Data 2)**. Given expanded profiling of pancreatic cell types in our study we determined the proportion of cREs not present in previous catalogs. Among cREs in our study, 9.4% and 7.4% were unique compared to a pan-tissue^29^ and pancreas-specific^5^ cRE catalog, respectively, such as a T cell-specific cRE directly upstream of *ZNF746* (**Figure 2E**). We identified cREs with cell type-specific activity by comparing accessible chromatin profiles across cell types (**Supplementary Data 3, see Methods)**. Cell type-specific cREs were enriched for sequence motifs of key cell type TFs as well as proximity to genes involved in cell type-specific function (**Supplementary Table 7,8**). For example, beta cell-specific cREs were significantly enriched (FDR<.10) for proximity to insulin secretion-related pathways and RFX, FOXA, NEUROD, and NKX6.1 TF motifs, whereas endothelial-specific cREs were significantly enriched for proximity to angiogenesis, blood vessel morphogenesis, and vasculature pathways and FLI, ETS, and ETV TF motifs (**Supplementary Table 7,8**). We also identified cREs specific to several of the sub-types within acinar cells; for example, ‘signaling’ acinar-specific cREs were enriched for JUN, FOS, and ATF motifs.

Due to the scarcity of immune populations in the pancreas, the epigenome of resident and infiltrating pancreatic immune cells has not been extensively described. In our study, we identified multiple immune cell types including T cells, macrophages, B cells and mast cells, although available cell numbers only enabled defining cREs in T cells and macrophages. T cell-specific cREs were significantly enriched for proximity to genes involved in T cell activation, T cell receptor complex, and cytokine receptor activity, and motifs for ETS, ETV and RUNX TFs, and macrophage-specific cREs were enriched for immune-related processes and PU.1 and SPIB motifs (**Figure 2F**). Compared to a previous study which profiled several whole pancreas donors, more than double the number of cREs were identified in each cell type (T cells: 58.8k vs 24.5k; Macrophages: 114.3k vs 55.7k). The increased number of cREs improved annotation of T1D-associated variants at immune-related loci; for example, at the *PRCKQ* locus likely causal T1D variant rs947474 (PPA=.88) from published fine-mapping data^5^ overlapped a pancreatic T cell and macrophage cRE not identified in these cell types in the pancreas previously, and not active in other pancreatic cell types (**Figure 2G**).

We next predicted networks of genes regulated by TF activity in each pancreatic cell type (**see Methods**). We linked cREs to target genes in each cell type using the activity-by-contact (ABC) method, which revealed an average of 46,474 cRE-target gene links per cell type, as well as based on promoter proximity (**Supplementary Data 4**). Using ABC and promoter proximity, genes were linked to, on average, 2.8 cREs per cell type (**Figure 2H**). We identified genes which had highly cell type-specific cRE links (**see Methods**), and genes with highly cell type-specific cRE links included key marker genes such as *INS* in beta cells, *GCG* in alpha cells, *IL2, IFNGR1* and *GZMA* in T cells, and *MARCOS* in macrophages. In each cell type, we next constructed gene regulatory networks (GRNs) for 366 TFs by combining (i) cRE-target gene links, (ii) TF sequence motif predictions in cREs, and (ii) TF and target gene expression levels (**Figure 2H**, **see Methods, Supplementary Data 5**). We then annotated likely cellular functions of TF GRNs by identifying biological pathways with gene sets that significantly overlapped TF GRNs. There were thousands of significant relationships linking TF GRNs to biological pathways across all cell types (Fisher’s test, FDR<.10) (**Supplementary Data 6**), which annotated many known regulators of biological pathway activity as well as many putative functions of TFs.

Finally, we utilized spatial transcriptomics data in combination with cell type-specific TF GRNs to infer TF activity within cell types and sub-types in the pancreas. Briefly, we used a univariate linear model to predict the observed gene expression based on TF–gene interaction weights, from which we scored TFs as active or inactive in each cell type^30^. We identified TFs with endocrine-specific activity in line with the previously described regulators of endocrine cell activity, such as *NEUROD1*, as well as high activity of *PAX6* in beta cells, where it is a key regulator of beta cell identify, function and survival^31^ (**Figure 2I,J**). Among other cell types, we inferred high activity for *BHLHA15*/*MIST1* in acinar cells, where it may play a role in the maintenance of pancreatic acinar identity^32^, and highly specific activity for *MEOX2* in endothelial cells and *RUNX3* in T cells (**Figure 2I,J**). Integrating GRNs with spatial transcriptomic profiles thus confirmed the specificity of key TFs regulating pancreatic cell types, including for TFs not measured on the spatial panel directly.

### Pancreatic cell type gene expression in T1D progression

Changes in genome-wide gene activity within each pancreatic cell type during progression to T1D are poorly understood. We therefore identified genes and biological pathways in each cell type with altered activity in ND AAB+ and T1D. To increase our power to detect changes in endocrine cell types, we also utilized single cell RNA-seq from purified islets of 48 non-diabetic, ND AAB+ (primarily single AAB+), and T1D donors from the HPAP consortium^4,33,34^. For each cell type and sub-type, we derived gene counts per sample, tested for differential expression in single and multiple ND AAB+ and recent and long-standing T1D compared to non-diabetes, and considered genes significant at FDR<.10 (**see Methods**). We further performed gene set enrichment of differential expression results for each cell type and sub-type and identified pathways with significant (FDR<.10) changes in activity in each condition (**see Methods**).

Marked gene expression changes were observed in beta cells in T1D (**Figure 3A**). In recent-onset T1D, 665 genes in beta cells had significant change (FDR<.10) in expression, where the most up-regulated genes included MHC class I and related (*CD74, B2M*) genes, cytokines and cytokine-induced genes (*IL15*, *GBP2, IFIT3*), cytokine-responsive TFs (*STAT1/4, IRF1*), and components of the 20S proteosome (**Figure 3B**, **Supplementary Figure 8**, **Supplementary Data 7**). We also observed up-regulation of MHC class II genes in T1D, particularly *HLA-DPB1.* At the pathway level, there was up-regulation of antigen processing and presentation, interferon signaling, interleukin signaling and JAK-STAT signaling, and down-regulation of oxidative phosphorylation, translation, mitochondrial function, mitosis, mRNA processing, protein folding and localization, ER-Golgi transport, and autophagy (**Figure 3C**, **Supplementary Table 9**). We examined whether specific pathways up-regulated in T1D showed heterogeneity in expression across beta cells, and several had evidence for bimodal expression patterns most prominently ECM-related pathways but also antigen presentation, while others such as interferon and JAK-STAT signaling did not (**Supplementary Figure 9**). Compared to recent-onset T1D, the largest changes generally differed in long-standing T1D (**Supplementary Figure 8**), where antigen presentation and class I MHC genes were less pronounced, interferon signaling was less pronounced although specific IRF TFs had higher expression, and class II MHC genes had stronger up-regulation. There was also stronger down-regulation in long-standing T1D of insulin secretion and beta cell function and genes such as *GLIS3* and *G6PC2* (**Supplementary Data 7, Supplementary Table 9**).

**Figure 3.**
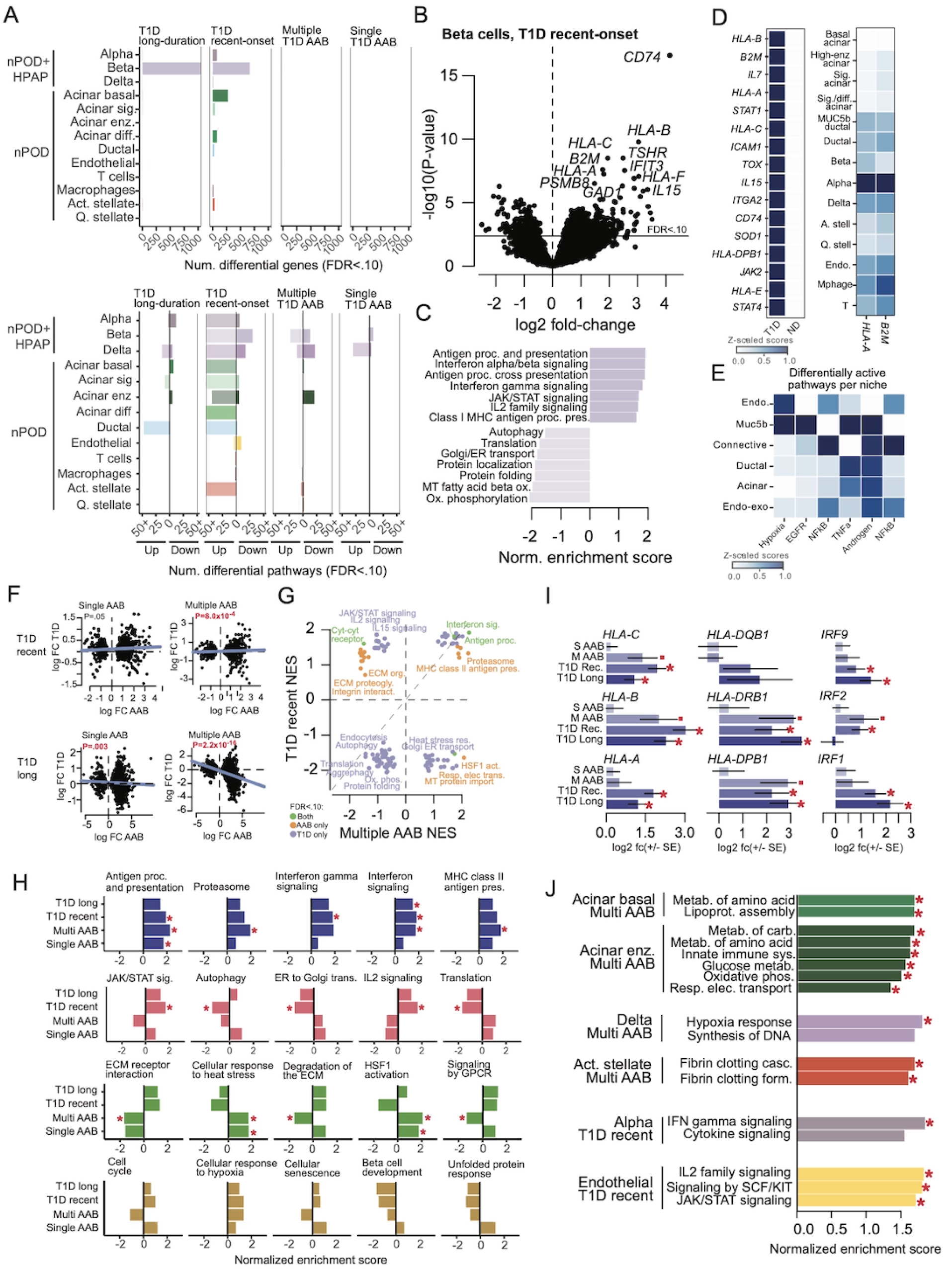
Cell type-specific changes in gene expression in T1D progression. (A) Number of genes in each pancreatic cell type with significant (FDR<.10) changes in expression in ND AAB+ or T1D status compared to non-diabetes. Endocrine cell types include scRNA-seq data from HPAP donors (top). Number of biological pathways enriched in genes with up- and down- regulated expression in each cell type in ND AAB+ or T1D (bottom). (B) Volcano plot showing differential expression results in beta cells in recent-onset T1D compared to ND. (C) Bar plot showing normalized enrichment score (NES) of biological pathways enriched in up- and down- regulated genes in beta cells in recent-onset T1D (bottom). (D) Scaled expression in spatial profiles of genes with up-regulated expression in T1D in beta cells (left). Spatially-dependent expression of selected genes (HLA-A, B2M) up-regulated in T1D in each cell type (right). (E) Biological pathways with differential expression within spatial niches in T1D compared to ND in spatial profiles. (F) Scatterplot of log fold-change in expression of genes in beta cells in single or multiple ND AAB+ compared to recent-onset T1D (top) and longer-duration T1D (bottom). Line shown in each plot is from a linear model of log fold-change values, and p-values are from Spearman correlation. (G) Normalized enrichment score (NES) of biological pathways enriched in differential expression results of recent-onset T1D and multiple ND AAB+. Pathways are colored based on significant enrichment (FDR<.10) in either, or both, disease states. (H) Normalized enrichment score of biological pathways enriched in differential expression results in beta cells across each T1D state (single ND AAB+, multiple ND AAB+, recent-onset T1D, and long-duration T1D) compared to non-diabetes. Red stars are for pathways with significant enrichment (FDR<.10) in each state. (I) Log fold-change of expression of selected MHC and interferon related genes in beta cells in each state compared to ND. Red stars indicate genes with significant change in expression (FDR<.10) and red dots indicate more nominal change in expression (un-corrected p<.05). (J) Normalized enrichment score (NES) of biological pathways enriched in genes with up- and down-regulated expression in ND AAB+ and T1D in other pancreatic cell types. Red stars indicate pathways with significant enrichment (FDR<.10).

Given marked changes in gene expression in beta cells in recent-onset T1D, we further characterized whether these pathways had altered activity within specific localizations in the pancreas. Of the genes with altered expression in beta cells in recent onset T1D and present in the spatial gene panel, almost all (95%) were up-regulated in T1D in spatial profiles (**Figure 3D**, **Supplementary Figure 10**). Furthermore, multiple up-regulated genes in T1D such as MHC class I genes (e.g. *HLA-A*, *B2M*) showed spatially-dependent expression patterns (Moran’s I >0.2) within endocrine, immune and ductal cells (**Figure 3D**). We further characterized pathways in recent-onset T1D with expression profiles dependent on specific niches and altered in T1D progression. We identified pathways in the PROGENy database in LIANA+^35^ to predict pathways preferentially active in a niche using a multivariate linear model. We identified multiple pathways with niche-dependent expression, including hypoxia in the endocrine niche (**Figure 3E**). When further assessing T1D-specific changes in pathway expression, pathways related to hypoxia and inflammation such as TNFa and JAK-STAT were differentially active in T1D (**Supplementary Figure 10**).

In contrast to T1D, few individual genes had significant changes in expression in beta cells in either single or multiple ND AAB+ (**Figure 3A**). We determined whether more subtle changes might be occurring at these stages. Genes altered in recent-onset T1D had significantly correlated effects in multiple ND AAB+, although not in single ND AAB+ (**Figure 3F**). At the pathway level, antigen processing and presentation was up-regulated in both single and multiple ND AAB+, and interferon signaling was up-regulated in multiple ND AAB+ (**Figure 3G,H**, **Supplementary Table 9**). Among key genes in these pathways, MHC class I genes (*HLA-A, HLA-B, HLA-C*) and interferon signaling *IRF* TFs were up-regulated in multiple but not in single ND AAB+ (**Figure 3I**, **Supplementary Data 7**). We also identified pathways altered specifically in single and multiple ND AAB+ and not in T1D; for example, heat stress response was up- regulated in single and multiple ND AAB+, extracellular matrix organization, cytokine-cytokine interactions, and GPCR ligand binding were all down-regulated in multiple ND AAB+, and TGF beta signaling was down-regulated in single ND AAB+ (**Figure 3H**, **Supplementary Table 9)**. Additionally, class II MHC antigen presentation was strongly up-regulated in multiple ND AAB+, but not single ND AAB+, including the class II MHC genes *HLA-DBP1* and *HLA-DRB1* (**Supplementary Data 7, Supplementary Table 9**). These results highlight that single and multiple ND AAB+ have both shared and distinct genomic changes in beta cells compared to T1D.

Changes have been reported in the exocrine pancreas in both T1D and at-risk individuals^21^ and in our study, we observed marked changes in exocrine cell gene expression in T1D progression. In ‘basal’ acinar cells, there were 276 genes with altered expression in recent-onset T1D, almost all of which (95%) had decreased expression (**Figure 3A**, **Supplementary Data 7**). Basal acinar and other acinar sub-types showed down-regulation of numerous pathways in recent-onset T1D including those related to signaling, stimulus response, metabolism, and protein transport (**Figure 3A**, **Supplementary Table 9**). In multiple ND AAB+, the ‘basal’ and ‘high-enzyme’ acinar sub-types showed higher expression of genes related to amino acid metabolism, which is necessary for enzyme production, as well as carbohydrate and glucose metabolism, insulin signaling, immune signaling, transcriptional activity, and respiration (**Figure 3J**, **Supplementary Table 9**). We also observed down-regulation of genes in ductal cells in recent-onset and long-duration T1D associated with small molecule transport, stimulus response, cytokine signaling and RNA processing, but no evidence for changes in ND AAB+ (**Supplementary Table 9**).

Other cell types in islets and the surrounding micro-environment also had significant changes in activity across entire pathways during progression to T1D. In alpha cells, while antigen presentation, interferon signaling, and other pathways were increased in T1D, in contrast to beta cells there were few changes in single or multiple ND AAB+ (**Figure 3J**, **Supplementary Table 9**). Delta cells showed more prominent changes in multiple ND AAB+, including increased hypoxia and heat stress response and cell cycle-related pathways and decreased signaling pathways, as well as in single ND AAB+ (**Figure 3J**, **Supplementary Table 9**). In endothelial cells there was increased IL2 and JAK-STAT signaling as well as SCF/KIT signaling, which promotes angiogenesis^36,37^, in recent-onset T1D (**Figure 3J**, **Supplementary Table 9**). In activated stellate cells, there was increased expression of genes associated with fibrin clotting and decreased expression of translation in ND AAB+, and down-regulation of many pathways in recent-onset T1D (**Supplementary Table 9**). While we did not observe evidence for significant changes in gene or pathway activity in immune (T cell, macrophage) cells, this could be due to the small number of cells profiled for these cell types.

Together, these results reveal key genes and pathways altered in pancreatic cell types in ND AAB+ and T1D donors with both shared and distinct changes in ND AAB+ compared to T1D, which in ND AAB+ included antigen presentation, interferon signaling, ECM-related and stress response pathways in beta cells and metabolism and immune signaling in acinar cells.

### Changes in the pancreatic cell type-specific epigenome in T1D progression

We next examined to what extent altered gene and pathway activity in pancreatic cell types in T1D progression is driven by changes in the epigenome of ND AAB+ and T1D donors using snATAC-seq profiles from 29 nPOD donors.

First, we identified cREs in each cell type with altered activity in T1D progression using a linear mixed model to account for pseudo-replication (**see Methods**). We observed significant changes (FDR<.10) in cRE activity in ND AAB+ and T1D for most pancreatic cell types (**Supplementary Data 8**). Beta cell cREs with increased activity in recent-onset T1D were significantly enriched (FDR<.10) for sequence motifs of steroid hormone receptors (NC3C1, NR3C2), NF-Y (NFYA, NFYB, NFYC), interferon response factors (IRF2, IRF7), and stress-response TFs (ATF4, STAT1, CEBPG) among others (**Figure 4A**, **Supplementary Table 10**). Conversely, cREs with decreased activity in T1D were significantly enriched for sequence motifs of TFs involved in core beta cell functions, such as HNF1 and RFX, with many beta cell identity TFs (NKX6.1, PDX1) and other TF families including FOXA and MEF showing more nominal enrichment (**Figure 4A**, **Supplementary Table 10**). We also identified sequence motifs enriched in beta cell cREs altered in ND AAB+, including IRF, TCF and STAT TF motifs in cREs with increased activity and MEF, RFX, and NFAT TFs in cREs with decreased activity, although other T1D-associated motifs such as HNF1 showed no change in ND AAB+ (**Figure 4A**, **Supplementary Table 10**). Sequence motifs were also enriched cREs altered in T1D progression for other pancreatic cell types, such as MEF and RFX TF motifs in alpha cells, RUNX TF motifs in activated stellate cells, STAT TF motifs in endothelial cells, and FOS/JUN motifs in ductal cells.

**Figure 4.**
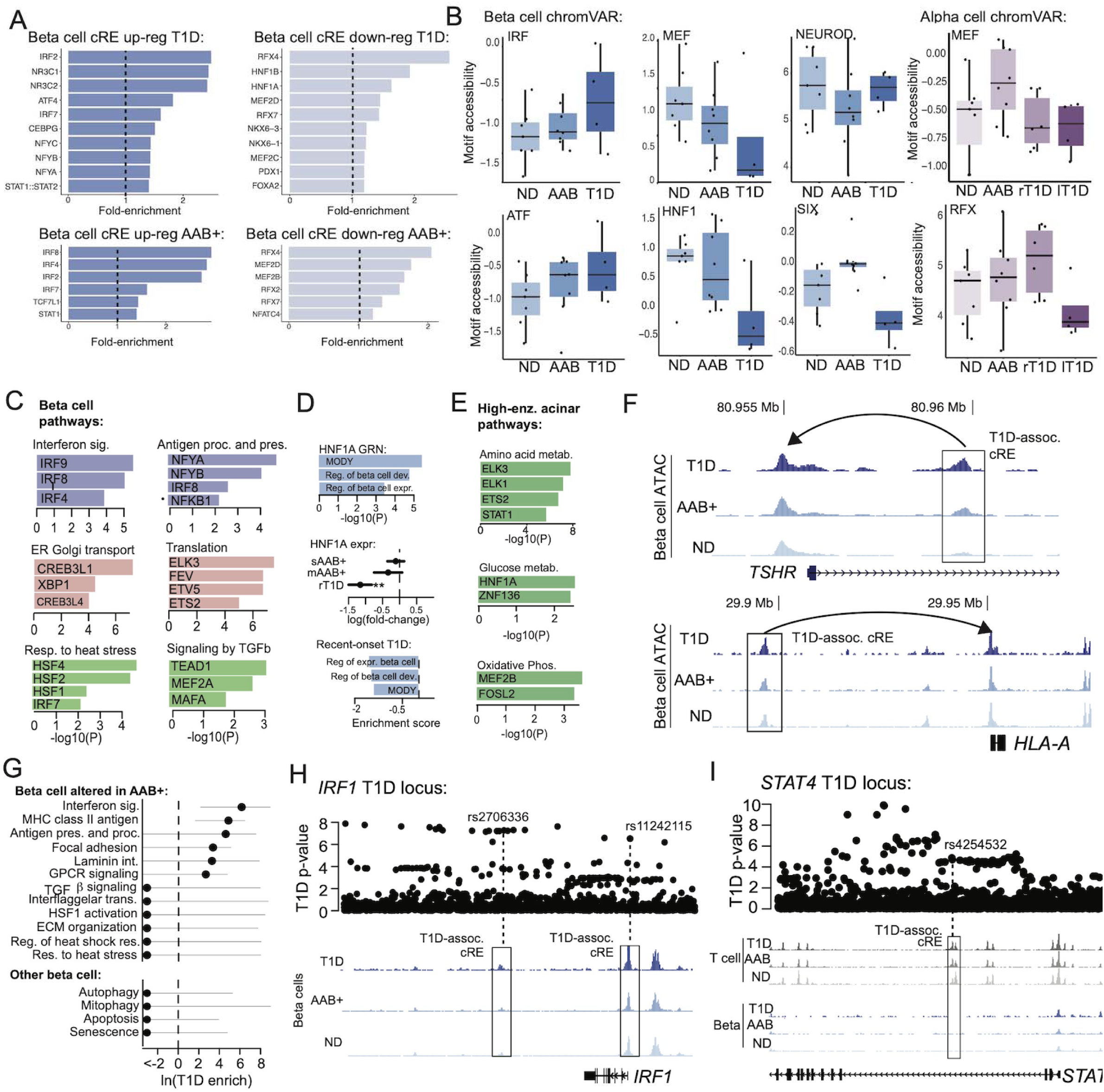
Epigenomic changes in pancreatic cell types in T1D progression. (A) Fold enrichment of sequence motifs for transcription factors (TFs) enriched in beta cell cREs with up- regulated or down-regulated activity in recent-onset T1D (top) or ND AAB+ (bottom). (B) Box plots showing donor-level genome-wide accessibility of selected TF motifs in beta cells (left) and alpha cells (right) from chromVAR across non-diabetes (ND), ND AAB+ (AAB) and recent-onset T1D (T1D). (C) TF GRNs enriched for overlap with genes in biological pathways in beta cells altered in T1D progression. (D) Biological pathways in beta cells enriched for overlap with the HNF1A GRN (top). Beta cell expression of HNF1A in T1D progression (middle). Beta cell activity of biological pathways linked to the HNF1A GRN in T1D progression (bottom). (E) Genome browser views of the *TSHR* (top) and *HLA-A* (bottom) loci where beta cell cREs with altered activity in T1D were linked to genes with concordant changes in expression in T1D. (F) Genome-wide enrichment of T1D-associated variants in beta cell cREs linked to pathways with altered expression in ND AAB+. (G) Genome browser view of T1D associated variants and beta cell accessible chromatin in non-diabetes, ND AAB+ and T1D at the *IRF1* locus, where candidate T1D variant overlaps a beta cell cRE with altered activity in T1D progression. (H) Genome browser view of T1D associated variants and both beta cell and T cell accessible chromatin in non-diabetes, ND AAB+, and T1D at the *STAT4* locus. Candidate T1D variants at this locus overlap a T cell cRE but not a beta cell cRE.

We determined next whether TF motifs enriched in T1D-associated cREs in pancreatic cell types had broader, genome-wide changes in activity in T1D progression by modeling sequence motif accessibility across individual cells using chromVAR^28^ (**see Methods**). In beta cells, we observed consistent changes in the genome-wide accessibility of specific sequence motifs in T1D progression, including increasing accessibility of IRF motifs and decreasing accessibility of RFX, FOXA, and MEF motifs from ND AAB+ to T1D states (**Figure 4B**, **Supplementary Table 11**). In other cases, sequence motifs had different patterns in ND AAB+ and T1D, such as decreased accessibility of HNF1 and increased accessibility of PAR-related and hormone receptor TFs in T1D only and opposed accessibility of SIX TFs in ND AAB+ and T1D. While alpha cells showed similar increases in accessibility of hormone receptor, stress-response, and PAR-related TFs in T1D progression as in beta cells, there were also several marked differences such as increased accessibility of MEF and RFX motifs in ND AAB+ and recent-onset T1D, respectively (**Figure 4B**, **Supplementary Table 11).**

We used TF GRNs to determine which TFs drive changes in pathway activity in T1D progression. In beta cells, pathways altered in ND AAB+ and T1D had highly specific links to TF GRNs, suggesting key regulators of pathway activity in T1D progression (**Figure 4C**, **Supplementary Data 6**). For example, pathways up-regulated in beta cells in T1D and ND AAB+ such as interferon signaling were linked to GRNs for IRF TF motifs and antigen processing and presentation were linked to NFY, IRF and NFkB TF GRNs, while down- regulated pathways in T1D such as ER and Golgi-related processes were linked to CREB3L1, XBP1 and other TF motifs (**Figure 4C**). We also identified TF GRNs linked to pathways altered specifically in ND AAB+, such as heat stress related pathways and HSF TF GRNs, extracellular matrix-related pathways and ETS, ELK and ELF TFs, and GPCR signaling pathways and RFX and FOXA GRNs (**Figure 4C**, **Supplementary Data 6**). While we observed a strong change in HNF1 motif accessibility, as well as HNF1A expression, in beta cells in T1D (**Figure 4B,D**), no pathways linked to the HNF1 GRN had significant change in expression in T1D. However, there was a more nominal change in beta cell development and function pathways linked to the HNF1 GRN in T1D (**Figure 4D**), supporting that reduced HNF1 activity likely underlies altered beta cell function in T1D, as has been shown in the context of type 2 diabetes^38^.

Similarly, in other pancreatic cell types, TF GRNs were linked to pathways with altered activity in ND AAB+ or T1D. For example, in enzyme-high acinar cells, metabolic pathways altered in ND AAB+ were linked to GRNs for specific TFs such as glucose metabolism and HNF1, amino acid metabolism and STAT1, and oxidative phosphorylation and MEF and FOS TF GRNs (**Figure 4E**, **Supplementary Data 6**). In activated stellate cells, fibrin-related pathways up-regulated in ND AAB+ were significantly linked to ELK, HOX, CEBP and other TF GRNs. In endothelial cells, IL2 and JAK-STAT signaling pathways up-regulated in T1D were strongly linked to NFkB (REL, RELA) and IRF TF GRNs, and SCF/KIT signaling was also linked to HOX family TF GRNs, among others. We further explored changes in TF activity inferred from spatial gene expression profiles of TF GRNs across cell types, which revealed increased activity of immune regulation, inflammation and signaling TFs (e.g. STAT3, RBPJ, FOSL2, JUND), and reduced activity of endocrine-related TFs (e.g. PAX6, GLI3, MAFA, INSM1, NEUROD1), in T1D compared to non-disease (**Supplementary Figure 11**).

We next annotated specific beta cell cREs altered in T1D progression with putative target genes and assessed changes in regulatory programs at specific loci. There were 114 beta cell cREs with altered activity in T1D progression linked to genes with significant changes in expression. For example, a beta cell cRE on chromosome 14 in the first intron of *TSHR* had increased accessibility in recent-onset T1D and was linked to *TSHR,* which had among the largest increases in expression in recent-onset T1D (**Figure 4F**). We identified similar cREs up- regulated in recent-onset T1D linked to genes with highly up-regulated expression including *HLA-A* (**Figure 4F)**, as well as *CD74, GAD1, IL15,* and *STAT1/4*. In other cases, we observed epigenomic changes in beta cells that may precede changes in expression of cognate target genes. For example, a cRE upstream of *IAPP* had reduced accessibility in early T1D although *IAPP* itself only had a significant decrease in expression in longer-duration T1D.

Given pathways and transcriptional regulators with altered cell type activity in T1D progression, we determined whether any changes prior to T1D onset showed evidence for a role in genetic risk of T1D. We tested for enrichment of cREs linked to genes in each pathway for T1D associated variants genome-wide (excluding the MHC locus) using fgwas^18,39^ (**see Methods**). In beta cells, several pathways altered in ND AAB+ were enriched for T1D-associated variants including antigen processing and presentation (log enrich=4.48), class II MHC antigen presentation (log enrich=4.74), and interferon signaling (log enrich=6.00) as well as several extracellular interaction-related processes (focal adhesion, laminin interactions) and GPCR signaling (**Figure 4G**). By comparison, multiple other pathways previously implicated in driving T1D risk in beta cells such as apoptosis, autophagy, mitophagy, and senescence, showed limited to no enrichment (**Figure 4G**). Among other cell types, we found evidence for enrichment of immune, metabolism, and transcription related pathways in ‘high-enzyme’ as well as ‘basal’ acinar cells (**Supplementary Figure 12**).

We further identified specific T1D risk loci that may alter regulatory activity of disease-enriched pathways in key cell types such as beta cells, T cells and other immune populations, and exocrine cells during T1D progression. We identified candidate causal variants at known T1D loci by overlapping cREs altered in T1D progression with published fine-mapping data^5^. In beta cells, multiple candidate causal variants at the *IRF1* locus overlapped cREs with increased activity in T1D including at the promoter and downstream of *IRF1* (**Figure 4H**, **Supplementary Table 12**). There was increased beta cell expression of *IRF1* through stages of T1D progression and *IRF1* is a driver of beta cell interferon responses, which is a pathway broadly enriched for T1D associated variants (**Figure 4G**). Conversely, at the *STAT4* locus we identified cREs with increased activity in beta cells as well as T cells, although candidate causal variants for T1D at the *STAT4* locus only overlapped cREs active in T cells (**Figure 4I**). This finding supports that while increased *STAT4* activity in beta cells is observed in T1D, the *STAT4* locus more likely affects T1D risk through altered T cell function.

Together these results reveal transcriptional regulators and networks altered in T1D progression, including those regulating pathways that likely play a causal role in the development of T1D such as antigen presentation, interferon signaling, and extracellular interactions in beta cells.

### Changes in pancreatic cell-cell signaling in T1D progression

External signaling between cell types is a key driver of changes in cell type-specific regulation and function, and therefore we finally identified cell-cell signaling interactions in the pancreas altered in T1D progression.

We first inferred cell-cell interactions in snRNA-seq data for non-diabetes, ND AAB+ and T1D using 1,939 ligand-receptor pairs in CellChat^40^ (**see Methods**), which revealed 87,650 interactions significant (FDR<.10) in at least one condition (**Supplementary Table 13)**. Grouping ligands into functional categories revealed classes of outgoing signals preferentially produced by each cell type; for example, hormones, neuropeptides, and cell adhesion molecules from endocrine cells, and enzymes from exocrine cells (**Supplementary Figure 13**, **Supplementary Table 14**).

We identified cell-cell interactions with changes in activity in T1D progression using a permutation test and considered changes significant at FDR<.10 (**see Methods**). Overall, there was a reduction in the number and strength of interactions in recent-onset and long-standing T1D compared to non-diabetes, which was largely driven by exocrine cells (**Figure 5A**). In both ND AAB+ and recent-onset T1D there was increased strength of interactions involving endocrine cells and other cell types, although the total number of interactions was reduced (**Figure 5A**). We further identified cell-cell interactions among cells in spatial niches and determined changes in T1D using spatial transcriptome profiles. We identified spatially co-expressed ligand-receptor (LR) interactions by Moran’s bivariant extension in SpatialDM^41^ using LR pairs from CellChat^40^. We compared the number of detected interactions considering each FOV as technical replicates of a donor and observed significant heterogeneity across donors and, like dispersed cell data, fewer interactions in T1D compared to ND donors (H-statistic=19.6, p=.0015) (**Figure 5A**).

**Figure 5.**
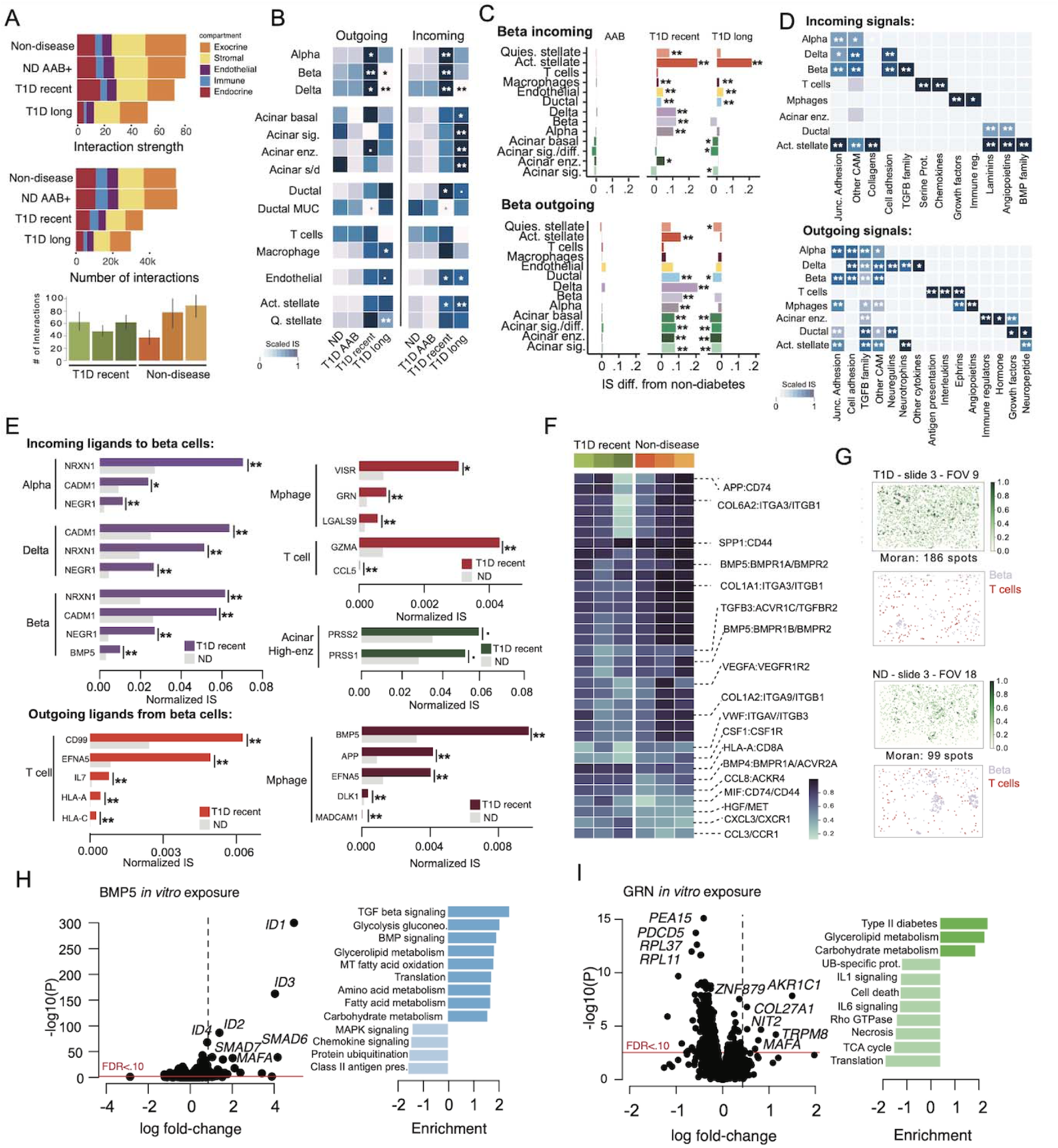
Cell-cell signaling networks altered in T1D progression. (A) Summary of total interaction strength (top) and number of interactions (middle) for each pancreatic cell type lineage in non-diabetes, ND AAB+, recent-onset T1D and long-duration T1D. Bar plot showing the number of ligand receptor interactions identified per donor in spatial slides (bottom). (B) Heatmap showing normalized interaction strength of outgoing and incoming signals for each cell type among donors which were non-diabetes, ND AAB+, recent-onset T1D and long-duration T1D. Stars represent the significance of the difference in interaction strength in each disease state compared to non-diabetes using permutations. **FDR<.01, *FDR<.05. (C) Difference in strength of interactions between beta cells and other pancreatic cell types and sub-types in ND AAB+, recent-onset T1D and long-duration T1D. **FDR<.01, *FDR<.05. (D) Interaction strength of outgoing and incoming signals for each cell type summarized by broad functional categories. **FDR<.01, *FDR<.05. (E) Normalized interaction strength in recent-onset T1D and non- diabetes for ligands with significant change in incoming or outgoing signal involving beta cells. (F) Heatmap per donor showing the interaction score of the top ligand-receptor interactions from a likelihood ratio test comparing ND and T1D donors. (G) Spatial plots of a representative FOV per condition (T1D: top, ND: bottom) highlighting, from left to right, spots where the interaction between HLA-C and CD8A presented a significant spatial pattern and the cell types where this interaction occurs. (H) Volcano plot showing genes with up- and down-regulated expression in EndoC-BH1 cells after treatment with BMP5 compared to no treatment (left). Biological pathways enriched in genes with up- and down-regulated expression in BMP5 exposure (right). (I) Volcano plot showing genes with up- and down-regulated expression in EndoC-BH1 cells after treatment with granulin (GRN) compared to no treatment (left). Biological pathways enriched in genes with up- and down-regulated expression after GRN exposure (right).

Among specific cell types, endocrine cells displayed significant increases in both outgoing and incoming signaling in recent-onset T1D (**Figure 5B,C**). We also observed significant increases in incoming signaling to endothelial, ductal, and activated stellate cells, as well as nominal changes in ‘basal’ and ‘high-enzyme’ acinar, immune, and stellate cells, in recent-onset T1D. Summarizing signaling by functional category revealed broad classes of cell type-specific signals altered in T1D; for example, beta and other endocrine cells had increased signaling from cell adhesion molecules, whereas T cells had increased antigen presentation and interleukin signaling (**Figure 5D**). We further examined changes in signaling between specific pairs of cell types in T1D progression (**Supplementary Table 15**). Significant changes (FDR<.10) in recent-onset T1D included increased incoming and outgoing signaling involving beta cells, including between beta cells themselves (**Figure 5C**), as well as increased signaling for alpha cells, outgoing signaling from ‘high-enzyme’ acinar cells and incoming signaling to endothelial cells.

Given the importance of external signaling to beta cells in T1D, we focused specifically on signals involving beta cells. In recent-onset T1D, autocrine/paracrine signals incoming to beta cells with significant changes in activity included cell adhesion molecules *NRXN1*, *CADM1,* and *NEGR1* from all endocrine cell types and the secreted factor *BMP5* from beta cells (**Figure 5E)**. In addition, high-enzyme acinar cells had increased signaling of trypsinogen (**Figure 5E)**, and stellate cells had increased signaling of ECM and cell adhesion molecules to beta cells. Among immune cells, signals with significant changes in signaling to beta cells included *GZMA* and *CCL5* from T cells and *VSIR*, *GRN*, and *LGALS9* from macrophages (**Figure 5E**). In return, beta cells had increased signaling of *IL7* and MHC class I genes *HLA-A* and *HLA-C* to T cells, as well as increased signaling of *BMP5*, *EFNA5*, *DLK1,* and *ANGPTL2* to macrophages. Notably, multiple beta-immune cell signals altered in T1D map to T1D risk loci (e.g. *DLK1, HLA-A, HLA-C, IL7R*)^18^.

We next identified differential interactions (p<0.05) in spatial profiles by performing a likelihood ratio test, which provided support for many T1D-associated interactions identified in dispersed cell data. For example, interactions involving HLA class I (e.g., *HLA-C*)*, APP, SPP1,* and *BMP5,* as well as ECM-related interactions, were altered in T1D (**Figure 5F,G**). We also identified additional interactions enriched in T1D donors, for example between migration inhibitory factor *MIF* and its transmembrane receptor *CD74*, consistent with previous studies^42^, and involving several chemokines. Next, we identified spatially-co-expressed ligand-receptor pairs using Morans’ I score in Liana+^35^. We obtained the top interactions associated with each niche using non-negative matrix factorization (**see Methods**). In T1D, an interaction between *APP* and *CD74* was enriched in the endocrine niche, where APP is involved in inflammation and could promote immune responses in T1D (**Supplementary Figure 14**). Conversely, interactions involving *INS*, *IGF1R*, and *INSR* and *CALM1*, among others, were enriched in the endocrine niche from non-disease donors (**Supplementary Figure 14**).

Several ligands altered in T1D progression BMP5 and GRN (granulin) are growth factors that have not been previously implicated in T1D. We determined the effects of *in vitro* exposure to these ligands on gene expression using the beta cell model EndoC-BH1. Exposing beta cells to BMP5 in culture revealed 1,926 genes with significant change (FDR<.10) in expression, where the most up-regulated genes were *ID1-4* and *SMAD6-7,* known targets of BMP that regulate proliferation and differentiation, and the beta cell identity gene *MAFA* (**Figure 5H**, **Supplementary Table 16**). More broadly, BMP5 exposure up-regulated pathways (FDR<.10) related to TGF beta signaling, glycolysis, secretion, and lipid metabolism, and down-regulated pathways such as antigen presentation and chemokine signaling (**Figure 5H)**. Second, granulin encodes secreted proteins produced by macrophages and ductal cells. Upon exposure to granulin, 491 genes had significant change (FDR<.10) in expression including up-regulation of beta cell function and insulin secretion genes *MAFA, ISL1*, *SOX4, CRY2* and down-regulation of apoptosis related genes *PEA15, PDCD5,* and *CCAR1* (**Figure 5I**, **Supplementary Table 17**). More broadly, granulin up-regulated pathways related to cholesterol and glycerolipid metabolism and down-regulated interleukin signaling and inflammation, transcription and translation, and cell death.

These results together reveal broad changes in predicted cell-cell signaling in T1D progression most prominently among endocrine cells and niches but also involving other cell types, including altered signals in T1D that modulate T1D-relevant regulatory programs in beta cells.

## Discussion

Single cell and spatial profiling of human pancreas donors revealed extensive changes in the abundance, regulation, and signaling of specific cell types in T1D progression, including processes that play a likely causal role in driving disease. In beta cells, class I and class II MHC antigen presentation and interferon signaling pathways, TF regulators of these pathways, and cREs linked to genes in these pathways all had up-regulated activity in recent-onset T1D and ND AAB+. Antigen presentation was altered as early as single ND AAB+ donors, suggesting that aberrant antigen presentation in beta cells may be an initial triggering event in T1D. Further, the heterogenous activity of antigen presentation pathways in beta cells suggests that subsets of beta cells may initiate the immune responses. Antigen presentation and interferon signaling pathways in beta cells were broadly enriched for T1D-associated variants and specific risk loci for T1D were linked to key genes in these pathways such as *IRF1*. In contrast, we found limited evidence that pathways directly related to apoptosis, as well as other processes implicated in T1D in beta cells such as autophagy, senescence, and mitophagy, harbor T1D risk, and are thus more likely consequences of disease. It has been long hypothesized that beta cells affect genetic risk of T1D through increased cell death^43–49^. Our results support that beta cells may primarily contribute to T1D risk via the initiation or exacerbation of immune responses, which necessitates different cellular models and phenotypic readouts to understand their role in disease.

In addition to shared pathways, gene activity in beta cells and other pancreatic cell types had distinct changes in ND AAB+ compared to recent-onset T1D, revealing that genomic profiles prior to T1D onset are only partially intermediate to those in T1D. In addition, the lack of individual genes with highly significant changes expression in ND AAB+ suggests that changes at these stages are likely more subtle, in contrast to previous reports^4^. Several pathways in beta cells were altered specifically in multiple and single ND AAB+ such as heat shock response and ECM organization. Heat shock responses are activated by a variety of stressors, promote antigen presentation in beta cells, and can act as chaperones for antigens and thus may plausibly contribute to the initiation of autoimmunity^50,51^. The breakdown of ECM is also an important process in T1D, as both a precursor to immune invasion as well as by affecting intrinsic beta cell function^52^. We observed a similar pattern of both shared and distinct changes in the epigenome of beta cells in ND AAB+ compared to T1D, including increased NEUROD1 activity and decreased SIX TF activity. There were also shared and distinct features in T1D based on the duration of disease; for example, a more pronounced reduction in beta cell function in long-standing T1D.

In contrast to beta cells, changes in gene activity in alpha cells were largely restricted to after the onset of T1D, including for antigen presentation and interferon response pathways and transcriptional regulators of these pathways. This supports that immune responses are more pronounced within beta cells compared to alpha cells prior to T1D onset, which may reflect differences in immune targeting as well as the intrinsic properties of each cell type. The latter is supported by multiple *in vitro* studies showing pronounced responses of beta cells to external T1D-relevant stressors^19^. A previous study revealed changes in alpha cell function and gene expression in single ND AAB+ donors using data from the HPAP consortium^53^, although there were overall few genes with altered expression in this study which supports our findings that genomic changes in alpha cells prior to T1D onset are likely subtle. In addition, several TF families such as RFX and MEF2 had different patterns of accessibility between alpha cells and beta cells in T1D progression, further highlighting the unique responses of each cell type to disease progression. Conversely to alpha cells, delta cells had altered activity of multiple pathways related to stress and inflammatory responses in single and multiple ND AAB+, as well as decreased abundance in T1D, suggesting they may play an as-of-yet unappreciated role in T1D progression.

Given that we profiled whole pancreas donors, our study was uniquely placed to reveal changes in the exocrine pancreas compared to previous single cell studies which used purified islets^4,15^. We identified multiple clusters of acinar and ductal cell types which had distinct genomic profiles and may represent heterogeneous sub-types of these cell types. In acinar cells, sub-clusters were broadly related to enzyme production and signaling responses, and previous reports highlighted similar heterogeneity in secretory and idling acinar cells^54^. Similar hormone producing and signaling states have been reported in endocrine cells^55^, and thus may represent a common property of secretory cells. Resolving exocrine sub-clusters revealed genomic changes within specific exocrine sub-types in T1D. Enzyme-producing acinar and *MUC5B*+ ductal cells were more abundant in ND AAB+ donors, and multiple acinar sub-types had altered metabolism, immune, and transcriptional pathways, as well as increased signaling to beta cells, in T1D progression. Specific pathways within acinar cells altered in T1D progression also harbored T1D-associated variants, further supporting a role for exocrine pancreas in T1D risk^4,18^ and providing new in-roads to determine how cellular processes in acinar cells contribute causally to T1D.

Signaling relationships between pancreatic cell types revealed incoming and outgoing external signals during progression to T1D. Cell-cell signaling between immune and beta cells highlighted known signals in T1D, such as granzyme B incoming to beta cells and class I MHC presented by beta cells^56,57^, as well as potential mechanisms of genes implicated in T1D genetic risk such as *DLK1* and *IL7* signaling from beta cells to immune cells^18^. Additional signals incoming to beta cells in T1D such as BMP5 and granulin have no prior known role in disease. BMP5 has increased autocrine/paracrine signaling in T1D and *in vitro* suppressed antigen presentation- and chemokine-related genes and enhanced expression of several genes linked to beta cell proliferation and function. Other BMP proteins have been shown to both enhance and inhibit beta cell function, maturity, and proliferation^58,58–60^, where the direction of effect may depend on the level of BMP signaling. Granulin suppresses class I MHC expression and T cell infiltration of ductal adenocarcinoma cells in the context of pancreatic cancer and has been shown to promote proliferation in mouse models of beta cells^61,62^. Signaling pathways altered in T1D, particularly those involved in T1D genetic risk, may represent therapeutic areas for preserving beta cell function to prevent or reverse T1D.

There were multiple limitations of our study that could be used to inform directions for future studies. For example, while we grouped non-diabetic donors by number of autoantibodies, there is additional granularity in stages of T1D stages; for example, stage 2 of T1D is marked by both autoantibody positivity and reduced beta cell function^63^. Future studies may therefore utilize islet functional measures to help refine characterization of T1D stages for genomic analyses. In addition, heterogeneity in T1D pathogenesis has been defined based on criteria such as first-developed autoantibody, HLA background, age and other factors^64–67^, and continued collection of larger sample numbers will enable understanding genomic changes within donors mapping to disease sub-groups. As immune subsets infiltrating the pancreas arise from pancreatic lymph nodes (pLNs)^68^, studies combining pLN data with pancreas data from matched donors will be valuable in understanding the role of immune cells in driving T1D progression in the pancreas. Finally, expanded spatially-resolved profiling of cells will continue to help reveal cell type-specific changes within disease-related cellular niches and neighborhoods.

In summary, our study revealed gene regulatory changes in pancreatic cell types in T1D progression and highlighted pathways, regulatory networks, and signals that may play a causal role in T1D; efforts that inform both new directions for mechanistic studies and novel targets for therapies to prevent or reverse T1D. We provide these data and maps in visualization tools (available at http://t1d-pancreas.isletgenomics.org) to further enhance their utility to the research community. More broadly, our study highlights the utility of single cell multiomics and spatial analysis to reveal insight into cellular processes underlying progression to complex disease.

## Methods

### Sample selection

Whole pancreas tissue was obtained from the Network for Pancreatic Organ Donors with Diabetes (nPOD) biorepository according to federal guidelines with informed consent obtained from each donor’s legal representative. Studies were considered exempt and approved by the Institutional Review Board (IRB) of the University of California San Diego. We selected 7 T1D donors with more recent onset (<1 year from diagnosis) and 5 T1D donors with longer duration (>5 years from diagnosis), along with 11 age- and sex-matched non-diabetic (ND) individuals. We also selected 9 non-diabetic donors with T1D autoantibodies (ND TD AAB+), the majority of which had multiple antibodies although one donor was single GAD+. In total, 32 donors were obtained for genomic profiling (**Supplementary Table 1**).

### Single cell assays Tissue homogenization

Flash-frozen pancreas tissue was homogenized using mortar and pestle on liquid nitrogen. ∼40 mg of ground tissue was used as input for the different single nucleus assays.

### Generation of single nucleus ATAC-seq data

Roughly 40 mg of ground pancreas tissue was resuspended in 1 ml of nuclei permeabilization buffer (10 mM Tris-HCl (pH 7.5), 10 mM NaCl, 3LmM MgCl_2_, 0.1% Tween-20 (Sigma), 0.1% IGEPAL-CA630 (Sigma), 0.01% digitonin (Promega) and 1% fatty acid-free BSA (Proliant, 68700) in molecular biology-grade water). Nuclei suspension was filtered with a 30-µm filter (CellTrics, Sysmex) and then incubated for 5 min at 4 °C on a rotator. Nuclei were pelleted with a swinging-bucket centrifuge (500L×L*g*, 5 min, 4 °C; Eppendorf, 5920 R) and washed with 1 ml wash buffer (10 mM Tris-HCl (pH 7.5), 10 mM NaCl, 3LmM MgCl_2_, 0.1% Tween-20, 1% BSA (Proliant, 68700) in molecular biology-grade water). Nuclei were pelleted and resuspended in 10 µl of 1x Nuclei Buffer (10x Genomics). Nuclei were counted using a hemocytometer, and 15,360 nuclei were used for tagmentation. snATAC-seq libraries were generated using the Chromium Single Cell ATAC Library & Gel Bead Kit v1.1 (10x Genomics, 1000175), Chromium Chip H Single Cell ATAC Kit (10x Genomics, 1000161) and indexes (Single Index Kit N Set A, 1000212) following manufacturer instructions. Final libraries were quantified using a Qubit fluorometer (Life Technologies), and the nucleosomal pattern was verified using a TapeStation (High Sensitivity D1000, Agilent). Libraries were sequenced on NextSeq 500, HiSeq 4000 and NovaSeq 6000 sequencers (Illumina) with the following read lengths (Read1L+LIndex1L+LIndex2L+LRead2): 50L+L8L+L16L+L50.

### Generation of single nucleus RNA-seq data

Roughly 40 mg of ground pancreas tissue was suspended in 500 µL of nuclei buffer: 0.1% Triton-X-100 (Sigma-Aldrich, T8787), 1× EDTA free protease inhibitor (Roche or Pierce), 1 mM DTT, and 0.2 U/µL RNase inhibitor (Promega, N211B), 2% BSA (Sigma-Aldrich, SRE0036) in PBS. Sample was incubated on a rotator for 5 min at 4 °C and then pelleted with a swinging bucket centrifuge (500× *g*, 5 min, 4 °C; 5920LR, Eppendorf). Supernatant was removed and pellet was resuspended in 400 μl of sort buffer [1 mM EDTA and RNase inhibitor (0.2 U/μl) in 2% BSA (Sigma-Aldrich, SRE0036) in PBS] and stained with DRAQ7 (1:100; Cell Signaling Technology, 7406). 75,000 nuclei were sorted using an SH800 sorter (Sony) into 50 μl of collection buffer [RNase inhibitor (1 U/μl) and 5% BSA (Sigma-Aldrich, SRE0036) in PBS]. Sorted nuclei were then centrifuged at 1000*g* for 15 min (Eppendorf, 5920R; 4°C, ramp speed of 3/3), and supernatant was removed. Nuclei were resuspended in 18 to 25 μl of reaction buffer [RNase inhibitor (0.2 U/μl) and 1% BSA (Sigma-Aldrich, SRE0036) in PBS] and counted using a hemocytometer. 16,500 nuclei were loaded onto a Chromium controller (10x Genomics). Libraries were generated using the 10x Genomics, Chromium Next GEM Single Cell 3’ GEM, Library & Gel Bead Kit v3.1 (10x Genomics, 1000121), Chromium Next GEM Chip G Single Cell Kit (10x Genomics, 1000120) and indexes (Single Index Kit T Set A, 10x Genomics, 1000213 or Dual Index Kit TT Set A, 10x Genomics, 1000215) according to the manufacturer specifications. Complementary DNA was amplified for 12 PCR cycles. SPRISelect reagent (Beckman Coulter) was used for size selection and cleanup steps. Final library concentration was assessed by the Qubit dsDNA HS Assay Kit (Thermo Fisher Scientific), and fragment size was checked using a TapeStation (High Sensitivity D1000, Agilent). Libraries were sequenced using a NextSeq 500 and a Novaseq6000 (Illumina) with the following read lengths (Read1L+LIndex1L+LIndex2L+LRead2): 28 + 8 + 0 + 90 (single index) or 28L+L10L+L10L+L90 (dual index).

### Generation of joint single-nucleus RNA and ATAC-seq data (Multiome)

∼40 mg ground tissue was resuspended in 1Lml of wash buffer (10 mM Tris-HCl (pH 7.4), 10 mM NaCl, 3LmM MgCl_2_, 0.1% Tween-20 (Sigma), 1% fatty acid-free BSA (Proliant, 68700), 1 mM DTT (Sigma), 1x protease inhibitors (Thermo Fisher Scientific, PIA32965), 1LULµl^−1^ RNasin (Promega, N2515) in molecular biology-grade water). Nuclei suspension was filtered with a 30-µm filter (CellTrics, Sysmex) and pelleted with a swinging-bucket centrifuge (500L×L*g*, 5 min, 4 °C; Eppendorf, 5920 R). Nuclei were resuspended in 400Lµl of sort buffer (1% fatty acid-free BSA, 1x protease inhibitors (Thermo Fisher Scientific, PIA32965), 1LULµl^−1^ RNasin (Promega, N2515) in PBS) and stained with 7-aminoactinomycin D (7-AAD; 1 µM; Thermo Fisher Scientific, A1310). A total of 120,000 nuclei were sorted using an SH800 sorter (Sony) into 87.5 μl of collection buffer (1LULµl^−1^ RNasin (Promega, N2515), 5% fatty acid-free BSA (Proliant, 68700) in PBS). Nuclei suspension was mixed in a ratio of 4:1 with 5x permeabilization buffer (50 mM Tris-HCl (pH 7.4), 50 mM NaCl, 15LmM MgCl_2_, 0.5% Tween- 20 (Sigma), 0.5% IGEPAL-CA630 (Sigma), 0.05% digitonin (Promega), 5% fatty acid-free BSA (Proliant, 68700), 5 mM DTT (Sigma), 5x protease inhibitors (Thermo Fisher Scientific, PIA32965), 1LULµl^−1^ RNasin (Promega, N2515) in molecular biology-grade water) and incubated on ice for 1 min before pelleting with a swinging-bucket centrifuge (500L×L*g*, 5 min, 4L°C; Eppendorf, 5920 R). Supernatant was gently removed, and ∼10 µl were left behind to increase nuclei recovery. A total of 650 µl of wash buffer (10 mM Tris-HCl (pH 7.4), 10 mM NaCl, 3LmM MgCl_2_, 0.1% Tween-20 (Sigma), 1% fatty acid-free BSA (Proliant, 68700), 1 mM DTT (Sigma), 1x protease inhibitors (Thermo Fisher Scientific, PIA32965), 1LULµl^−1^ RNasin (Promega, N2515) in molecular biology-grade water) was added with minimal disturbance of the pellet, and samples were centrifuged again with a swinging-bucket centrifuge (500L×L*g*, 5 min, 4L°C; Eppendorf, 5920 R). Supernatant was gently removed without disturbing the pellet, leaving ∼2–3Lµl behind. Approximately 7–10Lµl of 1x Nuclei Buffer (10x Genomics) were added, and nuclei were gently resuspended. Nuclei were counted using a hemocytometer, and 18,300 nuclei were used as input for tagmentation. Single-cell multiome ATAC and gene expression libraries were generated following manufacturer instructions (Chromium Next GEM Single Cell Multiome ATACL+LGene Expression Reagent Bundle, 10x Genomics, 1000283; Chromium Next GEM Chip J Single Cell Kit, 10x Genomics, 1000234; Dual Index Kit TT Set A, 10x Genomics, 1000215; Single Index Kit N Set A, 10x Genomics, 1000212) with the following PCR cycles: 7 cycles for pre-amplification, 8 cycles for ATAC index PCR, 7 cycles for complementary DNA (cDNA) amplification, and 12 cycles for RNA index PCR. Final libraries were quantified using a Qubit fluorometer (Life Technologies), and the size distribution was checked using a TapeStation (High Sensitivity D1000, Agilent). Libraries were sequenced on NextSeq 500 and NovaSeq 6000 sequencers (Illumina) with the following read lengths (Read1L+LIndex1L+LIndex2L+LRead2): ATAC (NovaSeq 6000), 50L+L8L+L24L+L50; ATAC (NextSeq 500 with custom recipe), 50L+L8L+L16L+L50; RNA (NextSeq 500, NovaSeq 6000), 28L+L10L+L10L+L90

### Quality control and filtering

Single nuclei ATAC data was processed and aligned to reference genome hg38, and duplicate reads were removed using Cellranger ATAC (version 1.1.0). Chromatin accessibility for each sample was quantified in 5kb genome windows as previously described^69^. Nuclei with less than 1000 unique ATAC-seq fragments were removed. Initial quality control was performed to retain cells in each sample using the following metrics unique usable reads > 5000, fraction promoters used > 0.01, TSS enrichment (TSSe) > 0.3 using scanPy v1.8.0. Doublets were removed using Amulet v1.0 per sample^70^. Single nucleus ATAC-seq datasets were processed and aligned to reference genome hg38, and duplicate reads were removed using Cellranger ATAC v1.1.0. Chromatin accessibility for each sample was quantified in 5kb genome windows as previously described^69^. Nuclei with less than 1000 unique ATAC-seq fragments were removed. Initial quality control was performed to retain cells in each sample using the following metrics unique usable reads > 5000, fraction promoters used > 0.01, TSS enrichment (TSSe) > 0.3. Doublets were removed using Amulet v1.0 per sample^70^.

Single nuclei RNA samples were processed using Cellranger (version 6.0.1) with reference genome hg38^71^. Individual samples were processed for quality initially by removing nuclei with less than 500 expressed genes. Doublets were detected for each sample using DoubletFinder (version 2.0.3) using an expected doublet rate of 4% for all samples^72^. In effort to reduce ambient RNA contamination largely driven by acinar cells, we utilized SoupX (version 1.6.2) and selected acinar marker genes, REG1A and PRSS1, to estimate contamination rates^73^. Gene expression count matrices were then corrected for this predicted contamination, these correct counts were used for both clustering and downstream analysis.

Paired multiome data was processed, aligned, and multiplet reads were removed using cellranger arc (version 2.0.0) with the reference genome hg38. Individual sample quality control was done using both modalities to remove low quality nuclei without a minimum of 500 expressed genes and 1000 ATAC-seq fragments. Ambient RNA contamination was removed using SoupX (version 1.6.2) using the same parameters as previously described. Doublets were detected and removed for both modalities using DoubletFinder (version 2.0.3) and Amulet (version 1.0), with the same parameters as above for single modality data^70,72^. Paired multiome datasets were processed, aligned, and multiplet reads were removed using cellranger arc (version 2.0.0) with the reference genome hg38. Individual sample quality control was done using both modalities to remove low quality nuclei without a minimum of 500 expressed genes and 1000 ATAC-seq fragments. Doublets were detected and removed for both modalities using DoubletFinder and Amulet, with the same parameters as above for single modality data^70,72^.

### Clustering

#### Gene expression

After individual sample quality control, high quality barcodes from single modality snRNA-seq and the RNA modality of our multiome data were clustered for 40 samples (32 snRNA and 8 snRNA multiome) using Seurat (version 4.3)^74^. Quality control metrics such as high mitochondrial percentage (>1%), high number of genes detected (>4,000 genes), and high number of RNA counts (>7,500) were used to remove low quality barcodes. A combined clustering was created using principal components (PCs) from PCA of gene expression. We used Harmony^8^ (version 1.0.3) to correct the PCs for batch effects across samples, sex, and sequencing technology. Clusters were removed with low number of cells (<10 cells) and with quality metrics such as number of detected genes and RNA counts lower than other clusters. Additional doublet cells were removed based on the expression of 2+ canonical markers from unrelated cell types.

We leveraged gene expression profiles specific to the wide array of pancreatic cells from previous work to broadly label each snRNA-seq cluster as one of the following types: alpha (*GCG*), beta (*INS*), endothelial (*PLVAP*), lymphatic endothelial (*FLT4*), ductal (*CFTR*), acinar (*REG1A*), stellate (*PDGFRB*), and variety of immune cells including T-cells (*CD3D*), macrophages (*C1QC*), and mast cells (*KIT*) (**Supplementary Table 2**). Using cell type markers previously used to annotate cell type and sub-type populations such as activated stellate (*COL6A3*) and quiescent stellate (*SPARCL1*) we were able to annotate these clusters. We identified previously characterized ductal subtype MUC5b ductal cells from the presence of known marker genes such as *MUC5B*, *TIFF3*, and *CRISP3*^54^.

Marker genes of acinar sub-clusters were identified using DESeq2^76^ (version 1.34), followed by gene set enrichment of sub-cluster marker genes in KEGG^77–79^ and REACTOME^80^ pathways using fGSEA^81^ (version 1.20). In brief this was done by first creating two sets of sample pseudo- bulk count matrices of SoupX corrected gene expression for each cell type, one set which has the summation of count per sample per gene for that cell type and another with the summation of counts per sample per gene for all other cell types. We then performed DESeq for each cell type by concatenating these two matrices as our input and using cell type as outcome variable with sample ID as a covariate.

#### Accessible chromatin

We first merged 40 samples (32 snATAC samples, 8 multiome snATAC samples) from 29 donors using read counts in 5kb windows using Signac^82^(version 1.9.0). We then performed latent semantic indexing (LSI) of the combined snATAC data using Signac^82^. Harmony (version 1.0.3) was used to correct for batch effects using the covariates sample, sex, and sequencing technology^75^. Clustering was performed on the batch-corrected PCs using graph-based Leiden clustering. We removed nuclei with a TSS enrichment (TSSe) score <2, and removed clusters with less than 10 cells or with overall lower quality metrics, such as fraction of read in peaks, number of ATAC fragments per barcode, and fraction of reads in promoters compared to other clusters. After an initial window-based clustering, we called peaks using MACS2^83^ (version 2.2.7.1) (parameters: -q 0.05 --nomodel --keep-dup all) on each cluster and then repeated the entire clustering process using a consensus set of peaks merged across clusters. Additional doublets were manually removed based off the presence of promoter accessibility of other cell type marker genes. This was done using 9 known marker genes (*INS, GCG, REG1A,REG2B, CTRB2, PRSS1,PRSS2, CFTR, C1QC*); promoter region was considered 2kb upstream of the TSS. Data was clustered again after the removal of doublets. To identify cell types, we first assigned gene names to peaks that overlapped 2kb upstream of TSS and gene body using the gene activity function in Signac and then determined gene activity in established marker genes for each cell type and sub-type.

We next performed label transfer on the snATAC object using our gene expression map as reference and the peak-based chromatin data as query in Signac. Due to the size of the chromatin data, prior to label transfer we randomly split the barcodes in the object into smaller subsets. We used the 2k most highly variable features from the gene expression map to derive transfer anchors using canonical correlation analysis (CCA). These anchors were then used to transfer to our chromatin map using the TransferData function in Seurat (version 4.3). After each subset object was done with label transfer, we merged the objects and re-clustered all the chromatin data together using the same methods described above. Finally, we removed cells with low prediction scores (max.predicted.score<0.5), and all cells passing this threshold were labelled with the predicted cell type annotation. For acinar cells, we summed the prediction scores of all acinar subtypes then filtered by a combined acinar max.predicted.score <0.5.

To determine the accuracy of label transfer, we utilized single cell multiome data where the identity of the accessible chromatin profile is already known from the paired gene expression profile. Since the gene expression and chromatin profiles for these nuclei were analyzed separately, we could use them as an independent check to determine how many barcodes were correctly labeled. We identified multiome barcodes present both chromatin and gene expression maps, and then calculated the percentage of accessible chromatin barcodes with matching cell type assignments in label transfer and from the paired gene expression profile. Due to the limited transferring of subtypes in the chromatin modality, we calculated a percentage at both the sub-type and primary cell type levels.

#### Generation of spatial transcriptomics data

Pancreatic tissue from six nPOD organ donors - three with T1D (6228, 6247, 6456) and three without diabetes (6431, 6339, 6229), matched by age and sex - was selected for spatial transcriptomic profiling on the CosMx platform (NanoString, Seattle, WA). For each donor, five consecutive FFPE tissue sections from the pancreatic body region were cut at a thickness of 4 microns. Sections #1, #2, #4, and #5 were mounted on the back of VWR Superfrost Plus Micro Slides, centered within the scanning area. After sectioning, the slides were air-dried overnight at room temperature, sealed, and immediately shipped with desiccant and ice packs to the NanoString facility (Seattle, Washington), where they were processed within two weeks of receipt. Section #3 was triple-stained for CD3, insulin, and glucagon using chromogen-based immunohistochemical staining using the Mach2 Double Stain 1/Mach2 Double Stain 2 HRP-AP Polymer Detection Kit according to the manufacturer’s instructions (Biocare Medical, Pacheco, CA) and chromogens used included Betazoid DAB (CD3), Warp Red (insulin), and Ferangi Blue (glucagon; all from Biocare Medical). Slides were then counterstained with hematoxylin. After staining, the slide was digitized at 20X magnification using an Aperio CS2 slide scanner (Leica Biosystems, Inc., Wetzlar, Germany), and this image served as a reference for field-of-view (FOV) selection during CosMx data processing. The FOVs were selected by prioritizing specific features such as insulitic islets, islets with few insulin-positive cells, insulin-negative islets, and areas of inflammation in acinar tissue. The gene panel used for spatial imaging included 1010 total genes, including a fixed panel of 1000 genes on the Human Universal Cell Characterization RNA Panel and 10 additional custom genes selected for this project. The imaging experiments using the CosMX platform were performed at NanoString (Seattle, WA). Cell segmentation was performed by NanoString using Giotto^84^, which included using immunofluorescence for glucagon to mark islets, CD3 or CD45 to mark immune cells, and PanCK for ductal cells + DAPI.

#### Quality control and transcriptomic clustering of segmented cells

For downstream analysis of spatial transcriptomes, we used the python toolkits Scanpy^85^ and Squidpy^24^. For each slide, we imported matrices containing the gene expression, metadata and positions of segmented cells. We defined a unique cell name and created a merged anndata object with data from all the slides. We adopted a standard filtering strategy, removing cell with less than 10 detected genes and removing genes detected in less than 300 cells. We then normalized the counts per cell, such that every cell has the same total count after normalization (1e6), and we log-transformed the counts.

#### Clustering of segmented cells and cell type annotation in spatial data

To cluster the segmented cells we first integrated the samples using scVI v1.1.2^86^. We performed integration by condition using the slide as a categorical covariate. We then used the latent representation to create a shared nearest neighbor graph and compute UMAP for two-dimensional visualization. We performed hierarchical clustering on the scVI latent space at resolutions of 0.5 and 0.7 and we identified 15 and 16 transcriptomic clusters for ND and T1D respectively. To annotate cell types, we identified marker genes enriched in each cluster for knowledge-based cell type annotation. We detected endocrine cells by hormone expression, beta (*INS*, *IAPP*) and alpha (*GCG*, *TTR*); we also identified exocrine cells positively expressing epithelial marker *EPCAM*, ductal (*SOX9*, *KRT19*) and acinar (*EGF*, *DLL1*, *JAG1*); we further annotated endothelial cells (*PECAM1*, *VWF*), fibroblasts (*VIM*, *COL1A1*), immune cells (*CD4*, *CD8A*), and mast cells (*CPA3*, *TPSAB1*).

#### Cell type label transfer from reference snRNA-seq data

To achieve a finer annotation on the spatial context, we transferred the cell type labels from the dissociated reference to the spatial data using spatial mapping function from moscot v0.3.5^87^. First, we performed pseudo-bulking of dissociated data using decoupler v1.6.0^30^. We found the optimal combination of parameters for the spatial mapping task by hyperparameter tunning per FOV and we used cosine distance between the modalities. For the annotation mapping, we selected the label of the annotated cell with the highest matching probability.

#### Identification of spatial cellular neighborhoods

Cellular neighborhoods in the spatial context were computed per FOV utilizing the squidpy^24^ function spatial_neighbors, where we utilized generic coordinates and considered 30 nearest neighbors.

#### Identification and annotation of multicellular spatial niches

To identify multicellular niches, we computed the covet representation implemented in envi v0.3.0^25^ per FOV. We used the default parameters, which included 64 genes to represent the covariance matrix. We then created a shared nearest neighbor graph using the covet representation and performed unsupervised Leiden clustering with a resolution of 0.2. To annotate the clusters, we evaluated the relative cell type abundance in each group per fov and performed hierarchical clustering. We aggregated ‘Acinar basal’, ‘Acinar High Enz’, ‘Acinar signal’ and ‘Acinar sig/diff’ subtypes in the acinar niche, ‘Ductal’ and ‘MUC5b ductal’subtypes in the ductal niche, ‘Alpha’, ‘Beta’ and ‘Delta’ subtypes in the endocrine niche and ‘Act stellate’, ‘Q. stellate’, ‘Endothelial’, ‘Macrophage’ and ‘T cells’ in the connective tissue niche.

*Downstream analysis:*

#### Final peak calling and signal tracks

Cell type specific set of chromatin peaks were derived using MACS2^83^ v2.2.7.1 on the final cell type annotations of our chromatin map using the following parameters -q 0.05 --nomodel --keep- dup all. These peak calls were used to accessible chromatin signal tracks in UCSC genome browser^88^.

#### Marker CREs

Cell type-specific cREs were derived for each cell type and subtype. We first created a set of union peaks across the whole dataset. This was achieved by limiting peak size for all called peaks to 300bp by centering any peaks larger than 300bp at their summit and extending coordinates 150bp in either direction. We then grouped peaks based on overlap to create clusters of peaks. Within each cluster, the peak with the highest read count at its summit was identified as the reference peak for the region. We then generated a list of peaks that did not overlap any of the reference peaks and iteratively identified additional reference peaks again until no peaks remained.

We used this set of union peaks to calculate two sets of sample level pseudo-bulk matrices per cell type as follows: first, we aggregated the number of ATAC fragments within peaks per donor per cell type, then for each cell type created a second matrix with the summation of fragments from all other cell types. Normalized counts matrices were generated by dividing number of fragments within a peak by total number of fragments for that sample in that cell type then multiplying by scaling factor (1e6). Cell type specific regulatory elements were then determined for each cell type by comparing the normalized counts matrix for a given cell type with the normalized counts matrix of all other cell types summed together. To test enrichment of a given peak for each cell type, we performed a logistic regression model using sample id as a covariate and corrected for multiple tests using the Benjamini-Hochberg correction method (FDR<0.1). We limited the marker cREs per cell type to the top 5,000 cREs ranked by fold-change. We performed sequence motif enrichment of marker cREs for each cell type compared to a background of all cREs in the cell type using HOMER^89^ v5.0.1 and retained enriched motifs at FDR<.01. We also tested for gene set enrichment in marker cREs using GREAT^90^(version 4.0.4).

#### Calculation of TPM

We derived gene expression profiles for each cell type by creating aggregate count matrices by donor per cell type. Using GENCODE v38 ^91^ GRCh38.p13 gene size annotations we calculated transcript per million (TPM) to normalize for gene size.

#### Cell type proportion changes

We first scaled the counts for each cell type in a sample to 10,000 total cells per sample. For several cell types we excluded samples with abnormally high counts (sample 6278 for beta and delta; sample 6393 for T cells and B cells; sample 6375 for MUC5b+ ductal cells). We then created a linear model of the log transformed counts as a function of disease status (ND, ND AAB+, recent-onset T1D, long-duration T1D), age, sex, and BMI, as well as a linear model without the disease status variable. We performed comparison of the nested models using a likelihood ratio test in package lmtest^92^ in R and considered p-values from the test significant at .05.

#### Differential gene expression

To determine disease-related changes in gene expression, we performed differential analysis using DESeq2^76^ v1.34. Using the snRNA- seq data, we derived pseudo-bulk count matrices for each cell type by aggregating all barcodes of a donor for each gene on a per cell type basis. We created the count matrices from the SoupX^73^ corrected expression counts, and then rounded counts in the matrix to the nearest integer. We included sex, age, and BMI, as well as proportion of beta cells, as covariates in the model. For endocrine cell types, we included expression counts from scRNA-seq of 48 donors from the HPAP consortium^93^ derived from a previously created single cell map^34^, and included an additional covariate in the model for cohort. For a given cell type, we only used samples with at least 20 cells, except for long-duration T1D beta cells where we included all samples. In addition, genes were only tested for a cell type if detected in at least 2 samples per tested condition and if there was total of at least 10 counts across all tested conditions. We further excluded genes for each cell type that are established marker genes for a different cell type. Multiple test correction was performed using Benjamini-Hochberg correction and we considered genes significant at FDR<.10.

#### Differential cRE accessibility

Using cell type specific peak calls from MACS2^83^ v2.2.7.1 per cell type we created peak by barcode fragment count matrices all snATAC-seq donors for each disease condition. Lowly accessible peaks were removed from analysis, as determined by the average accessibility of peak across all samples less than median accessibility of all peaks across all samples. In addition, for each cell type samples were removed with less than 10 barcodes in that cell type. Lastly, cell types with less than 10 cells were not used in this analysis. We tested each disease condition against non-diabetic using glmer^94^ in R using the logistic regression model [Peak accessibility ∼ Disease + scale(FRiP) + scale(count) + (1|Sample)] using a binary peak count matrix. We used the fixed covariates of fraction reads in peak (FRiP) and ATAC fragment count (count) to account for sequencing depth variation and used sample ID as a random effect to adjust for sample variation. We used sample as random effect to mediate the pseudo-replication side effect of barcode level analysis. Cell types with more than 30k cells were subsampled down to 10k for this analysis. Disease related fold change was calculated by the following formula: (mean(disease peak accessibility) /mean(non-diabetic peak accessibility)). Multiple test correction was performed using the Benjamini-Hochberg method and we considered cREs significant at FDR<.10.

#### Pathway enrichment during T1D using gene expression input

To test for pathways enriched by disease, we performed gene set enrichment analysis (GSEA)^95,96^. Using the results from our differential expression analysis input genes were ranked using the following formula (-log10(pvalue)*log2FoldChanges), and fGSEA^81^ v1.20 was run using both KEGG^77–79^ and REACTOME^97–103^ databases [parameters: eps=0.0, minSize = 0, maxSize = 1000]. Enriched pathways were filtered down using an FDR cutoff of 10%.

#### Motif Enrichment

We used chromVAR^20^ to measure z-scored motif accessibility in snATAC-seq data. To do so, we prepared peak count data for input to chromVAR by converting the fixed peak sparse count matrix into a SummarizedExperiment and estimated GC content bias using chromVAR’s built in method^20,21^. Human TF motifs from JASPAR 2022^22^ were accessed using the JASPAR2022 Bioconductor package^23^ and motifs were annotated to peaks using motifmatchr^24^. The SummarizedExperiment and motif annotations were used as inputs into chromVAR’s computeDeviations function to derive GC bias corrected motif accessibility z-scores.

#### Motifs Enriched in Cell Types

TF motifs were filtered for those with an accessibility >1.2 based on chromVAR’s built in computeVariability function. Cell types with fewer than 50 cells were excluded. Cell type motif accessibility z-scores were averaged and plotted with pheatmap^25^ and RColorBrewer^26^.

#### Motifs Enriched in Acinar Subtypes

After sub-setting the motif matrix to barcodes from acinar cells, we averaged motif accessibility of each acinar subtype per sample then tested each motif using a two-way ANOVA across acinar subtypes also including a donor variable. We then calculated FDR on the p-values using qvalue^104^. To identify which specific subtype a significant motif was most enriched in, motifs were further tested using a two-way ANOVA comparing motif accessibility within the subtype to the average motif accessibility for the other acinar subtypes together also including a donor variable. P-values for each motif were corrected by Holm’s method. Motifs were annotated to sub-clusters based on being significant in the pan-subtype ANOVA, significant in the post-hoc ANOVA with Holm’s correction, and having the highest average deviation score in the given cluster.

#### Motif Differential Accessibility

To identify motifs with differential accessibility across disease states we used a linear mixed model using the lmerTest package^105^. We identified motifs in a cell type enriched in cREs with altered activity in ND AAB+ or T1D. For these motifs, accessibility was modeled by barcode using encoded variables to contrast autoantibody, recent-onset and long-duration T1D independently against non-diabetic controls. Scaled fractions of reads in peaks and scaled number of counts were used as fixed effect covariates and a random effect for sample was used to control for pseudo-replication. Samples with less than 10 cells in the given cell type were excluded and cell types with fewer than 50 cells, or disease states with fewer than 20 cells and 3 samples were not tested. We obtained p-values from the resulting models. Motif accessibility was averaged by sample and disease state to make boxplots. Average motif accessibility per condition was generated by averaging sample average motif accessibility and volcano plots were generated by comparing difference in motif accessibility vs negative log10 q values, with a difference threshold of 0.25 and q-value cutoff of 0.05 (5% FDR) for dashed lines and coloring and labeling of samples.

#### Motif enrichment in differential accessible CREs

To identify TF motifs enriched in cREs differential accessibility in each cell type, we used HOMER^106^ (version 5.0.1). For each cell type, we identified cREs with nominal association (uncorrected p<.05) and split cREs by fold change as input, and user HOMER function findMotifsGenome with a background of all cREs for the cell type with a size parameter of 200 and a masked version of the human genome hg38. Multiple test correction was done using the Benjamini-Hochberg method, and significant motifs were considered at FDR<.10.

#### ABC analysis

To link cREs to target genes we used Activity-by-contact (ABC)^107^ v0.2. This was done by first creating .bam files for each cell containing only barcodes from the accessible chromatin map. Since the HiC reference panel used was in hg19 genome build, cell type bams and peaks were coverted to hg19 using CrossMap^108^ v0.6.3, and we called peaks for each cell type with MACS2 v2.2.7.1 using this genome build. To further improve enhancer activity prediction, we used publicly available H3K27ac ChIP-seq data for acinar, ductal, alpha, beta, and delta cells^109^. We predicted candidate regions and enhancer activity for each cell type using the following flags: -- peakExtendFromSummit 250, --nStrongestPeaks 150000, and all genes with a TPM greater than 1 as ubiquitously expressed genes. After ABC analysis, links were converted back to hg38 using CrossMap. We identified genes with cell type-specific cRE link profiles by calculating the proportion of the total number of ABC links for that gene by cell type and calculating Shannon entropy based on the proportion.

#### Constructing TF gene regulatory networks

To determine gene regulatory networks (GRNs), we constructed units of transcription factors linked to cCREs linked to genes. We first used a position frequency matrix (PFMatrixList object) of TF DNA-binding preferences from the JASPAR 2022 database^110^ and width-fixed peaks^111^ as input to perform TF binding motif analysis. We used the ‘matchMotifs’ function in the R package motifmatchr^112^ v.1.21.0 to infer cell type specific cREs bound by each TF. We linked cREs bound by each TF to target genes based on proximity to the gene promoter (±5 kb of a TSS in GENCODE V19 or through Activity-by-contact (ABC) links ^107,113^ at a score cutoff of .015. TF GRNs were retained for analysis if the network includes fewer genes then the 90^th^ percentile of number of genes linked to a given TF. In addition to ensure TF GRNs were active in the associated cell type, we removed any TF GRNs with an average TF expression (TPM) less than 5.

#### Identification of cell type specific TF-modules and pathways enrichment

For each pancreatic cell population, we identified pathways and TF modules enriched using our identified marker CREs. In brief this was done for each cell type by deriving CREs associated with KEGG and REACTOME paths using the bedtools intersection TF-module gene linked CREs with union peaks accessible in that cell type. These union peak based pathways were tested for enrichment using fGSEA. We used the logistic regression marker CRE results to rank peaks using the following formula (-log10(pvalue)*log2FoldChanges). Similarly, we tested for TF modules enriched in each cell type by defining union peaks associated with a TF either proximally or through ABC; then using the logistic regression marker CRE results to rank peaks and test for enrichment using fGSEA. For both analyses, we used the Benjamini-Hochberg method for multiple test correction and retained results with an FDR < .10.

#### Identification of TF GRNs linked to biological pathways altered in T1D

To identify regulators of enriched pathways for each cell type, we next tested enrichment of each TF-module in pathways identified in our fGSEA analysis. We performed Fisher’s exact test to test for overlap in genes in each TF GRN and genes in each biological pathway in KEGG and REACTOME for each cell type. We performed multiple test correction using FDR and considered TF GRNs linked to a pathway at FDR<.10. Next, we filtered results to biological pathways with significantly altered expression in T1D and TF motifs belonging to TF sub- families with differentially accessibility in T1D from chromVAR^114^ results.

#### Genetic association enrichment

We tested for enrichment of T1D associated variants using summary statistics from a published genome-wide association study^5^. We defined groups of cREs in multiple ways; first, by identifying all cREs in each cell type linked to genes in each biological pathway in KEGG and REACTOME using ABC and promoter proximity links and, second, by identifying cREs in each cell type in GRNs for each TF. We calculated Bayes Factors (BFs) for each variant with minor allele frequency (MAF)>.05 genome-wide, excluding all variants at the MHC locus, using the method of Wakefield^115^. We then tested for enrichment of T1D associated variants in groups of cREs genome-wide using fgwas v0.3.6^39^ with a block size (-k) of 2,500.

We also tested for enrichment of fine-mapped T1D risk variants using finrich^116^, which compares the cumulative posterior probability of a set of variants in an annotation to a null distribution drawn from permutations of a background set of annotations. For each enrichment analyses using subsets of cREs in a cell type, we used the posterior probabilities in credible sets from a previously published GWAS^5^, the full set of cREs for the cell type as background, and 10,000 permutations.

We overlapped cREs in each cell type with credible sets of variants at known T1D signals from a published fine-mapping study. We further determined which cREs had at least nominal evidence (uncorrected p<.05) for differential accessibility in ND AAB+ or T1D.

#### Cell-cell interactions

The gene expression data was pre-filtered prior to running CellChat^40^ v1.1.3. First, any cell type represented by fewer than 20 cells for a sample was excluded. Next, cell types that appeared in fewer than two samples within a control or disease group were excluded from that group.

We considered a ligand expressed in a specific cell type if the average expression of the ligand in the cell type was greater than half the standard deviation (SD) of its average expression across all cells in at least two samples. After applying these filters, we ran CellChat using the RNA data slot of the Seurat object across the entire CCdb with default parameters except for ‘trim = 0’ in the “computeCommunProb” command and ‘thresh=1’ in the “subsetCommunication” command^40^. Each control and disease group were processed independently. Ligands from the CellChat database (CCdb) were grouped into high level categories by manual curation using UniProt^117^ and GeneCard^118,119^ (listed in **Supplementary Table 14**). Gene families were downloaded using biomart^120^ in R.

Results from different conditions were consolidated and subjected to FDR correction using the Benjamini-Hochberg method with the q-value^104^ package. Predicted interactions were considered with an FDR<0.1 and an IS above the second quartile were considered for downstream interpretation. To remove residual background contamination due to highly expressed genes, the following interactions were blacklisted in all cell sources except the ones listed: *INS* in beta cells, *GCG* in alpha cells, *SST* in delta cells, *PRSS1/2/3* in acinar cells, *CD8A, CD8B, CD8B2* in T cells, and *CD4* in T cells and macrophages.

To assess the significance of differences between conditions, we randomly permuted sample IDs among conditions and re-performed the CellChat analysis 100 times and comparing these outcomes with the observed CellChat results. The permutations were produced and filtered using the identical parameters as those for the observed data. Next, we aggregated the Interaction Strength (IS) across different “units” by summing all Ligand-Receptor (LR) pairs within a unit and normalizing this sum by the number of significant interactions for each condition (for example, the total of all incoming ligands to Beta cells in non-diabetic samples divided by the number of significant interactions identified in that condition). We then quantified the difference in effect size (IS-effect size) across contrasts: ND AAB+ vs. ND, recent-onset T1D vs. ND, and long-duration T1D vs. ND. A p-value was calculated by comparing the observational results against the simulations using the formula: the number of instances where simulation IS-effect size exceeded observational IS-effect size, divided by the number of permutations. Subsequently, p-values were corrected for multiple tests using the Benjamini-Hochberg method. We considered only interactions with an FDR less than 0.10 as significant.

#### Functional Analysis of spatial genomics profiles

We inferred TF and pathway activities utilizing the package Liana v1.1.0^35^. For TF activity inference, we use the cell type-specific GRNs derived from single cell multiome. We then fit a univariate linear model to infer the interaction wights. To identify cell type-specific TFs we performed a t-test overestimating the variance of each group and filtered TFs according to an adjusted p-value <0.05. We inferred pathway activities using the PROGENy model^35^. We used weights of the top 500 responsive genes ranked by p-value. We then fit a multivariate linear model to obtain the weights corresponding to pathway interactions. As with the TF activity analyses, we identified cell type specific pathways by performing a t-test overestimating the variance of each group.

#### Cell-Cell communication

We analyzed cell-cell communication in spatial transcriptomic data using SpatialDM v0.2.0^41^. We performed the analysis per condition, and per donor, having each FOV as technical replicate. For this study, the parameters l and cutoff were set to 100, and 0.2 to represent the spatial context. Additionally, we computed the weight matrix using the single-cell mode and we extracted the ligand-receptor interactions from the CellChat database^40^. To compute the global Morans’ I score and the local spot detection, we used the z-score method.

#### EndoC-Bh1 stimulation experiments and RNA-sequencing

A total of 25,000 EndoC-BH1 cells were seeded in media composed of DMEM (Corning, 10014CV), 2% BSA (Sigma, A1470), 3.5 × 10^−4% 2-mercaptoethanol (Gibco, 21985023), 0.12% Nicotinamide (MilliporeSigma, 481907), 5.5 ng/mL transferrin (MilliporeSigma, T8158), 6.7 pg/mL Sodium Selenite (Sigma, 214485), and 1% Penicillin-Streptomycin (Gibco, 15140122) on a 96-well (CellTreat Scientific Products, 229105) plate coated with ECM (Sigma, E1270) and Fibronectin (Sigma, F1141). The recombinant protein concentrations used were: 1ug/ml PGRN and 50ng/ml BMP5. EndoC-βH1 cells were obtained from Human Cell Design. RNA was isolated using the RNeasy Mini Kit (Qiagen) from EndoC-Bh1 cells either stimulated or unstimulated with each ligand. Samples included three replicates each for PGRN and its untreated controls, and six replicates each for BMP5 and its untreated controls. RNA integrity was assessed using a 2200 TapeStation (Agilent Technologies), and all samples achieved an RNA Integrity Number (RIN) greater than 7. Ribodepleted total RNA libraries were prepared using the TruSeq Stranded Total RNA Library Prep Gold kit (Illumina, Catalog #20020599) and sequenced at the UCSD Institute for Genomic Medicine on an Illumina NovaSeq S4 platform.

#### Bulk RNA-seq analysis

Quality control of the sequencing data was assessed using FastQC^121^. Transcript quantification was performed using Salmon^122^ with default parameters and the hg38 reference indexes. Counts were imported into R using the tximport^123^ package, and genes with fewer than 10 reads were excluded. Differential gene expression analysis was conducted using DESeq2^76^, applying a false discovery rate (FDR) threshold of less than 0.1. For pathway enrichment analysis, the fGSEA package was employed using the “stat” column from DESeq2 results. fGSEA analysis was restricted to gene sets containing more than 10 and fewer than 500 terms. Pathways were corrected for multiple testing using FDR with a threshold of 0.1, and only pathways belonging to the KEGG^77,79^ or REACTOME^80^ databases were considered.

## Code availability

All code used for this article has been made available at https://github.com/Gaulton-Lab/nPOD and https://github.com/theislab/spatial_pancreas

## Data availability

Raw sequence data will be made available in GEO upon publication and spatial imaging data will be available in Zenodo. Single cell maps can be visualized at http://t1d-pancreas.isletgenomics.org.

## Author contributions

R.L.M. performed single cell analyses and wrote the manuscript. S.J. performed spatial analyses and wrote the manuscript. W.E., L.T., E.B., C.M., K.K, R.E., H.M., J.C., E.G., and A.H. performed single cell analyses. C.Z. and D.B. performed sample collection and processing.

G.W. performed single cell and spatial analyses. M.M. generated single cell genomics data.

K.V. performed in vitro treatment experiments. I.K. and M.A. performed sample collection and processing. S.P. supervised single cell data analysis and contributed to the design of the study.

F.J.T. supervised the spatial data analyses. M.S. designed and supervised the study. K.J.G. designed and supervised the study, performed analyses, and wrote the manuscript.

## Acknowledgements

The work in this study was funded by DK120429 and DK122607 to K.J.G. and M.S. and T32 GM00866 to R.L.M. This research was performed with the support of the Network for Pancreatic Organ donors with Diabetes (nPOD; RRID:SCR_014641), a collaborative type 1 diabetes research project supported by JDRF (nPOD: 5-SRA-2018-557-Q-R) and The Leona M. & Harry B. Helmsley Charitable Trust (Grant#2018PG-T1D053). The content and views expressed are the responsibility of the authors and do not necessarily reflect the official view of nPOD. Organ Procurement Organizations (OPO) partnering with nPOD to provide research resources are listed at https://npod.org/for-partners/npod-partners/. This manuscript used data acquired from the Human Pancreas Analysis Program (HPAP-RRID:SCR_016202) Database (https://hpap.pmacs.upenn.edu), a Human Islet Research Network (RRID:SCR_014393) consortium (UC4-DK-112217, U01-DK-123594, UC4-DK-112232, and U01-DK-123716).

---

**Supplementary Figure 1:**
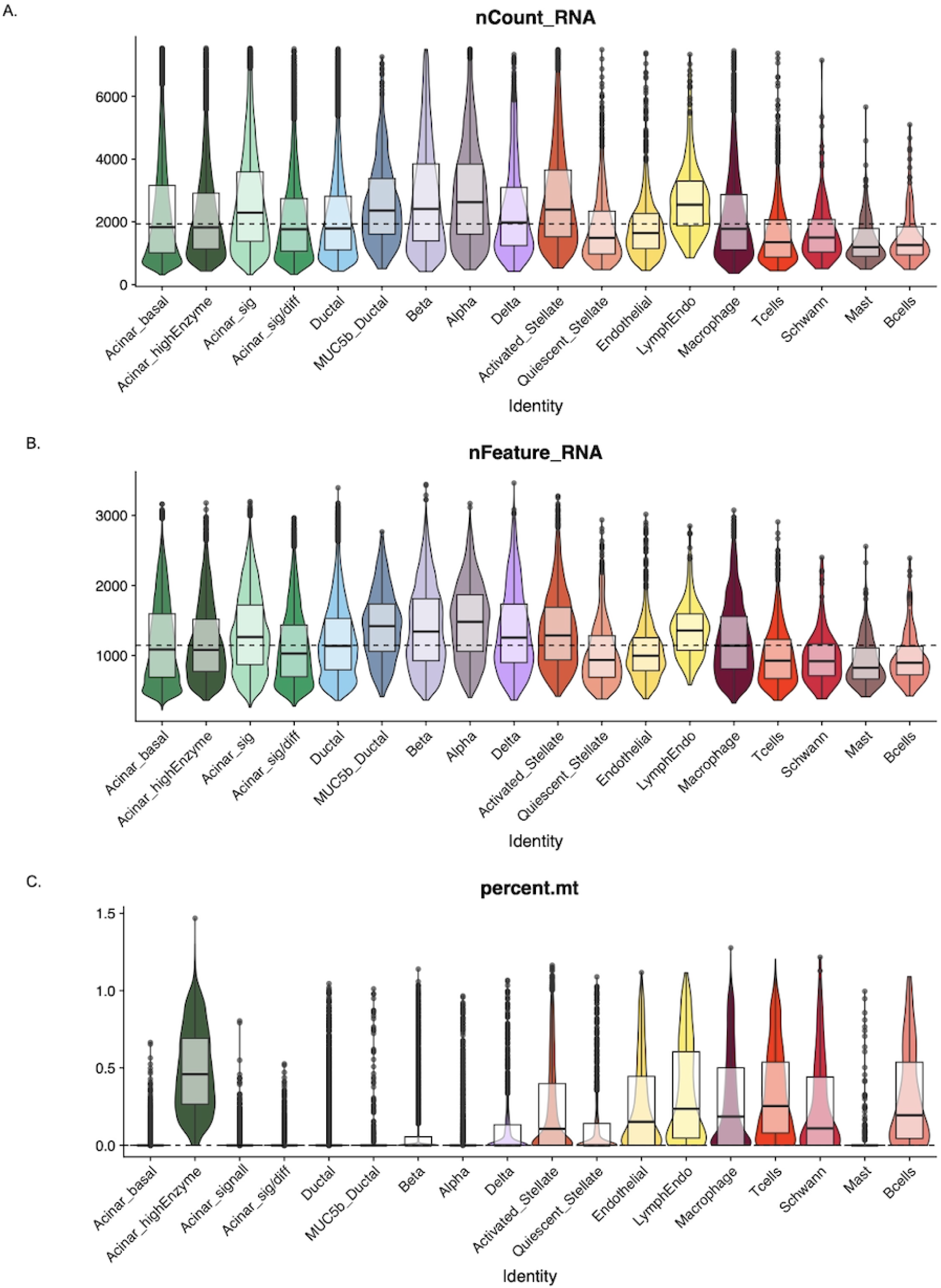
Violin plots of quality control metrics for RNA expression data including (A) counts, (B) genes per nuclei, and (C) percent mitochondria.

**Supplementary Figure 2:**
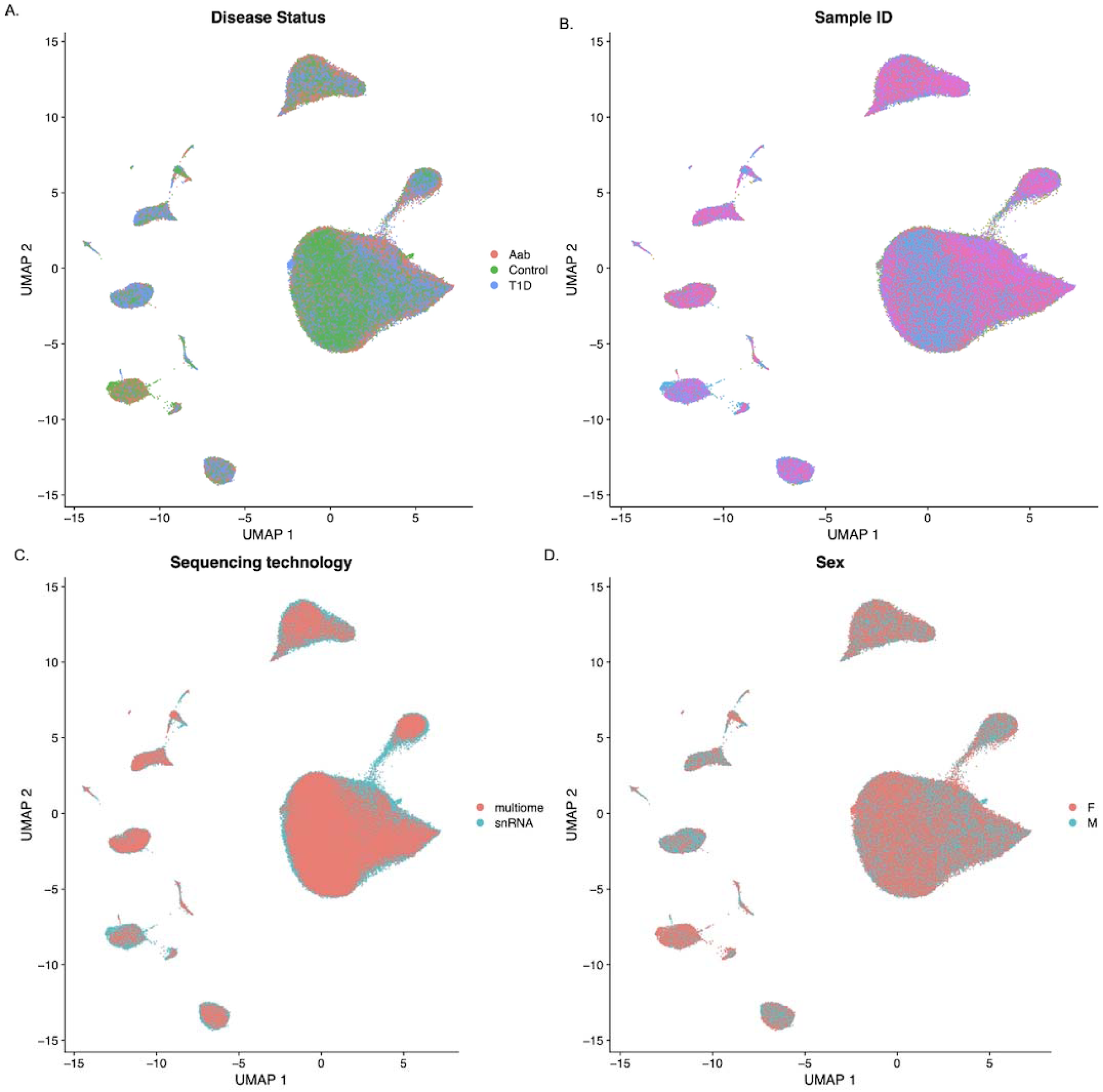
Uniform manifold approximation and projection (UMAP) of gene expression data annotated by (A) diabetes status, (B) donor ID, (C) sequencing technology, and (D) sex.

**Supplementary Figure 3:**
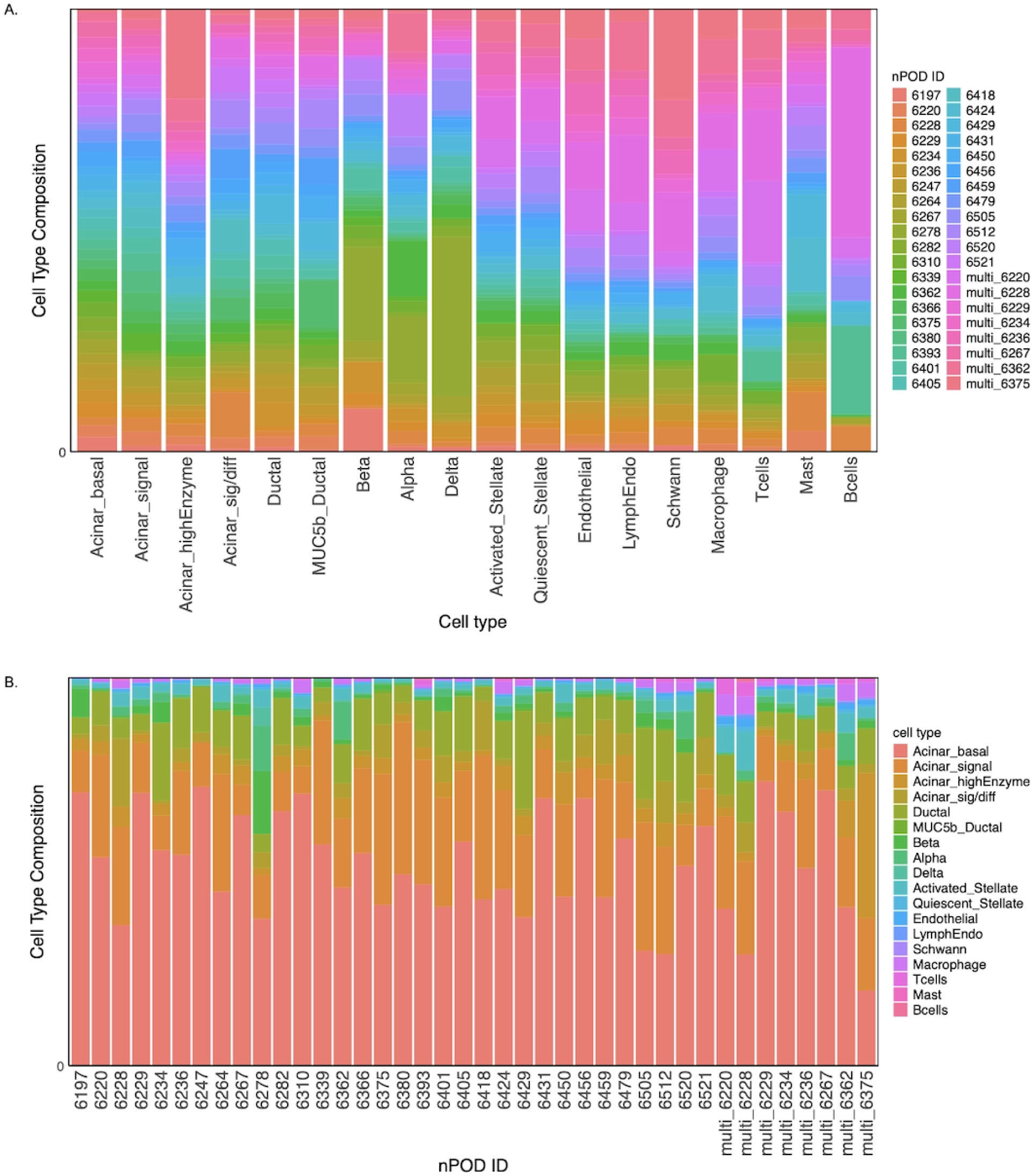
Proportion of cells: (A) per cell type per donor, (B) per donor per cell type

**Supplementary Figure 4:**
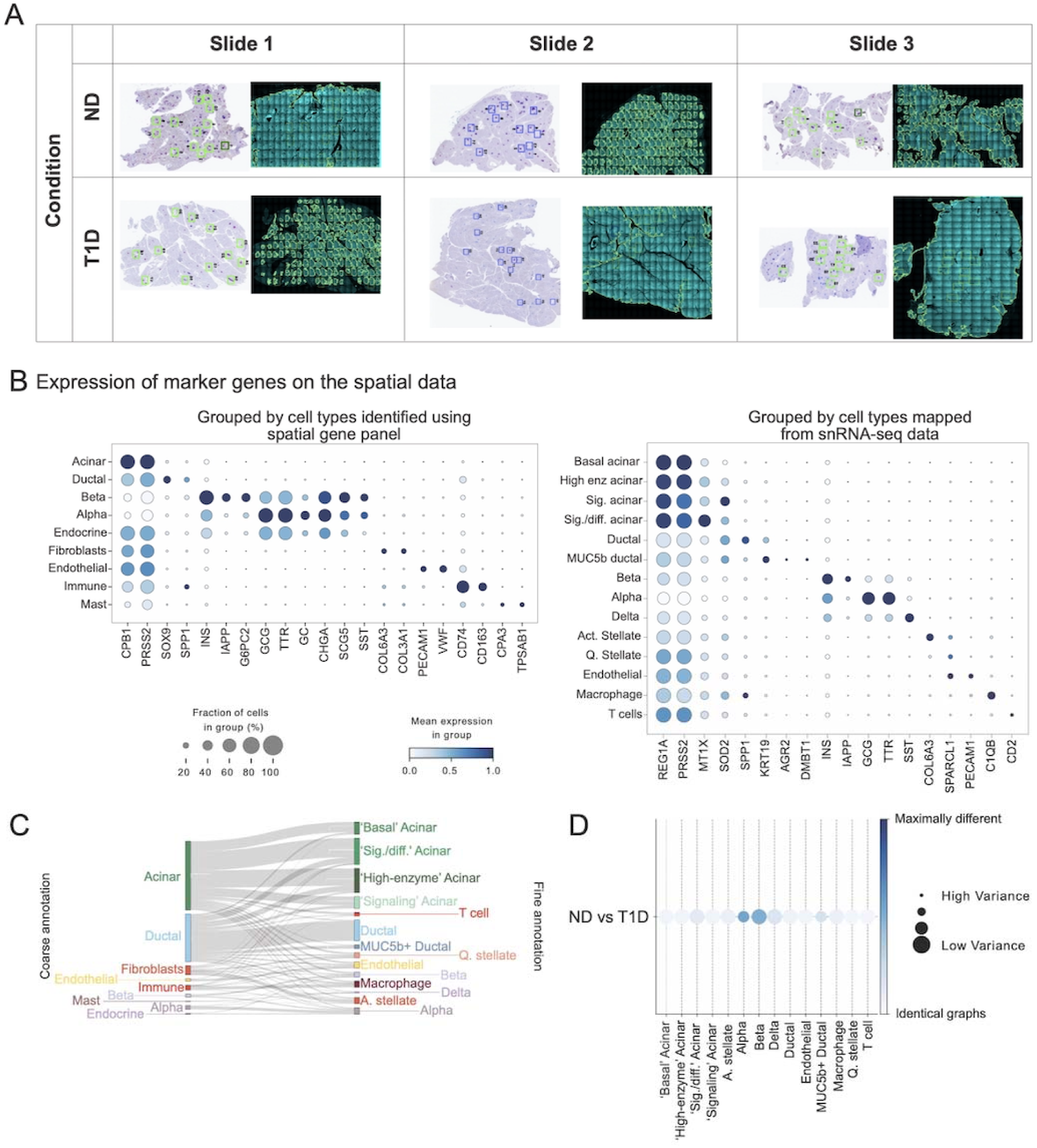
A. Wole slide images of the sections profiled with CosMx. B. Dotplots of the cluster expression of canonical markers used for cell type annotations at different levels of granularity in the spatial slides. Coarse annotation (left), finer annotation (right). C. Sankey plot showing the relative mapping from coarse annotation to fine annotation. The mapping was done using optimal-transport­based method moscot. D. Cell-type-specific subgraph comparison across conditions (ND vs. T1D), The size of the dot is indicative of the dissimilarity score variance over samples. The larger the dot size, the lower the score variance and the higher the score confidence is.

**Supplementary Figure 5:**
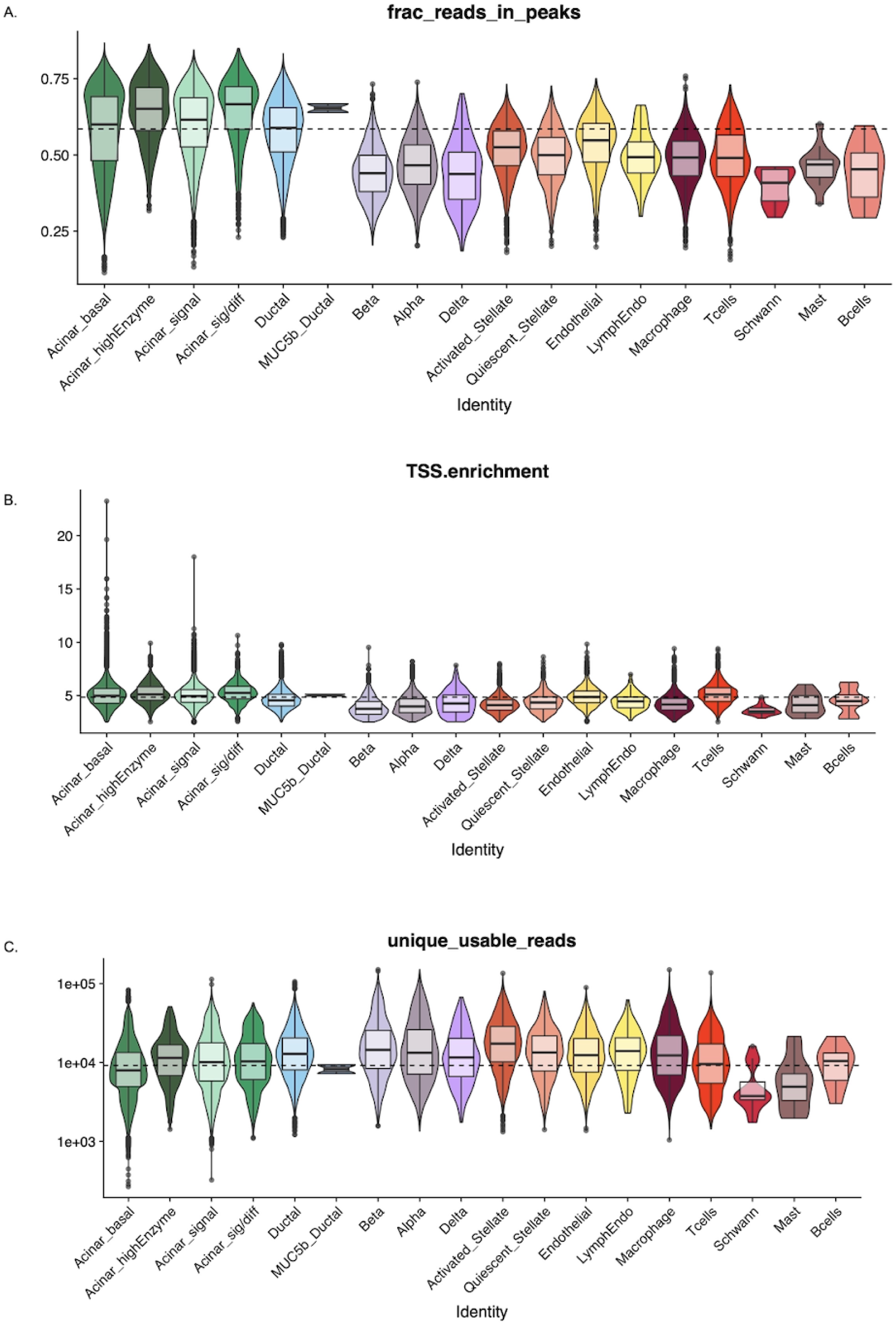
Violin plots of quality control metrics for chromatin accessibility data including (A) fraction of reads in peaks, (B) transcription start site enrichment (TSSe), and (C) percent mitochondria

**Supplemental Figure 6:**
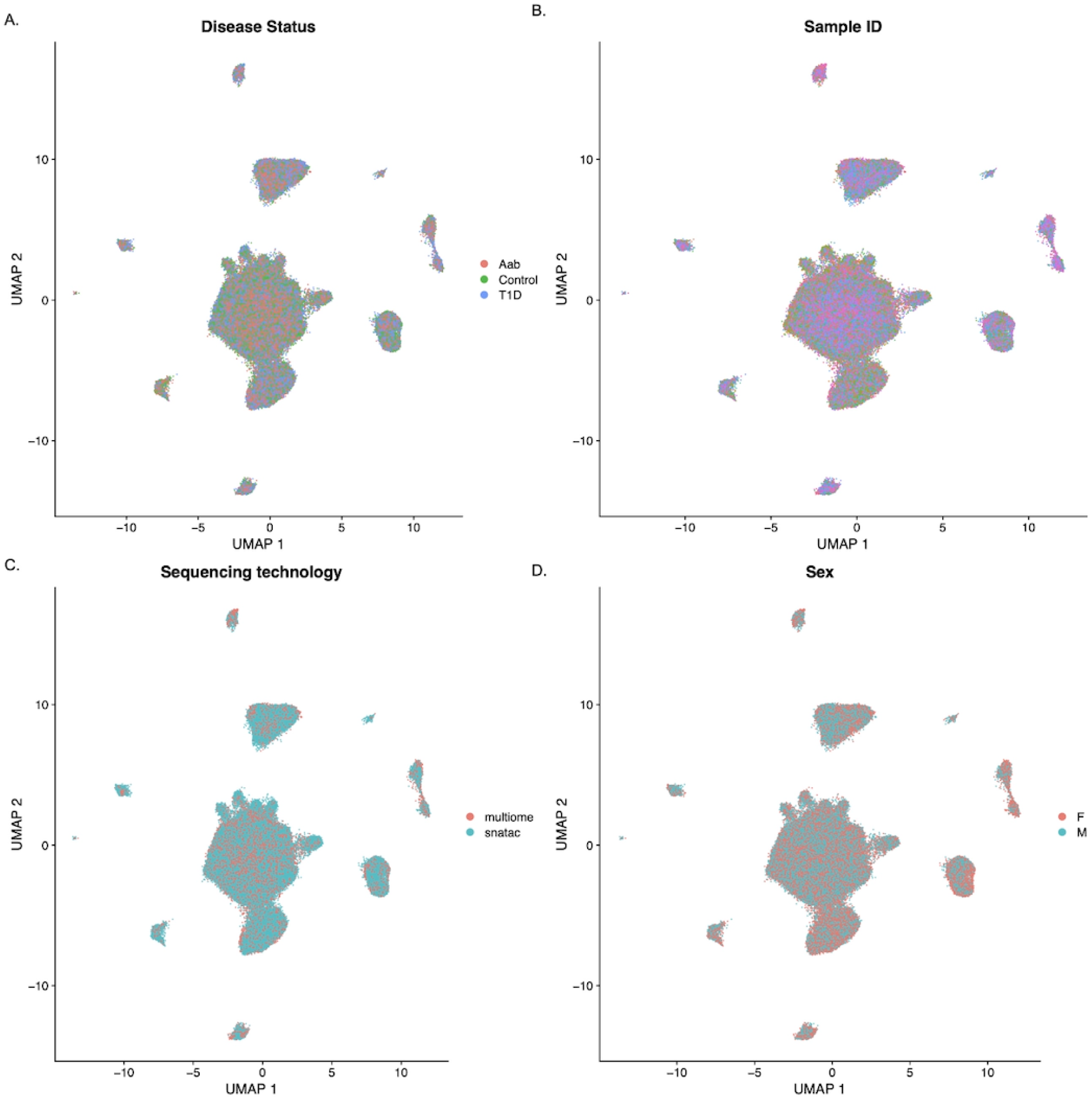
Uniform manifold approximation and projection (UMAP) of chromatin accessibility data annotated by (A) diabetes status, (B) donor ID, (C) sequencing technology, and (D) sex

**Supplemental Figure 7:**
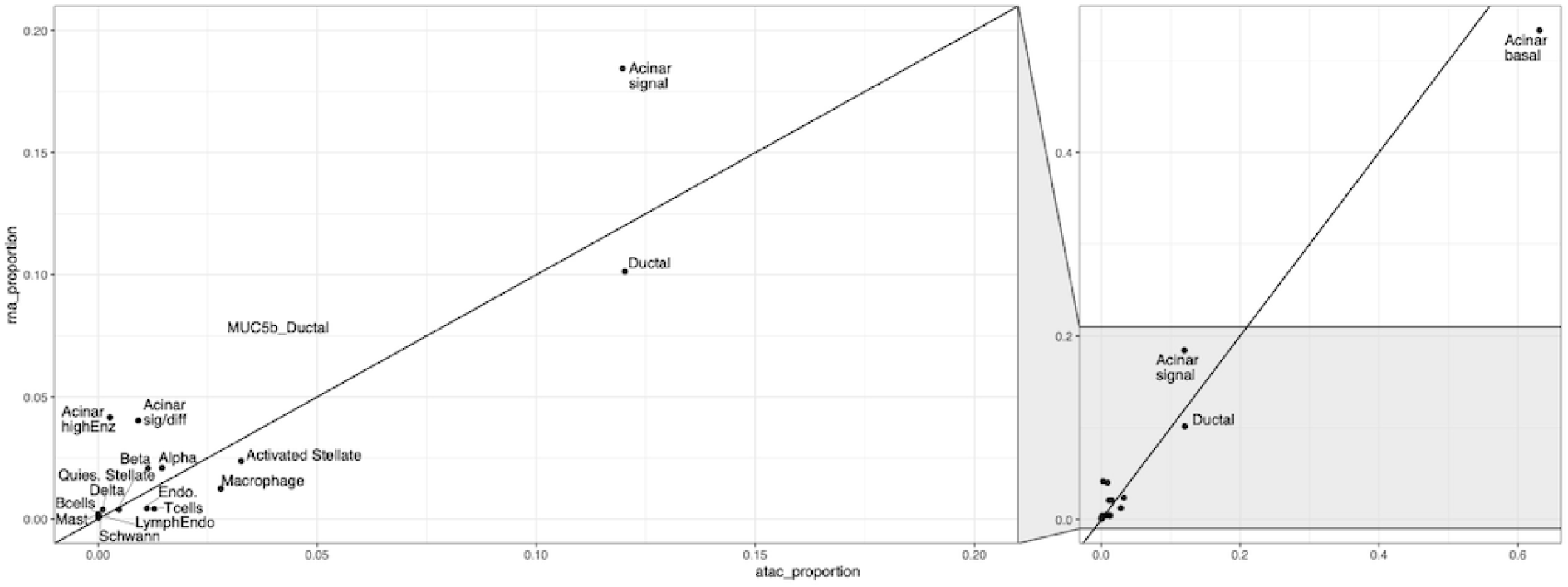
Scatterplot of cell type proportion between gene expression and chromatin accessibility data. Y=X line plotted

**Supplemental Figure 8:**
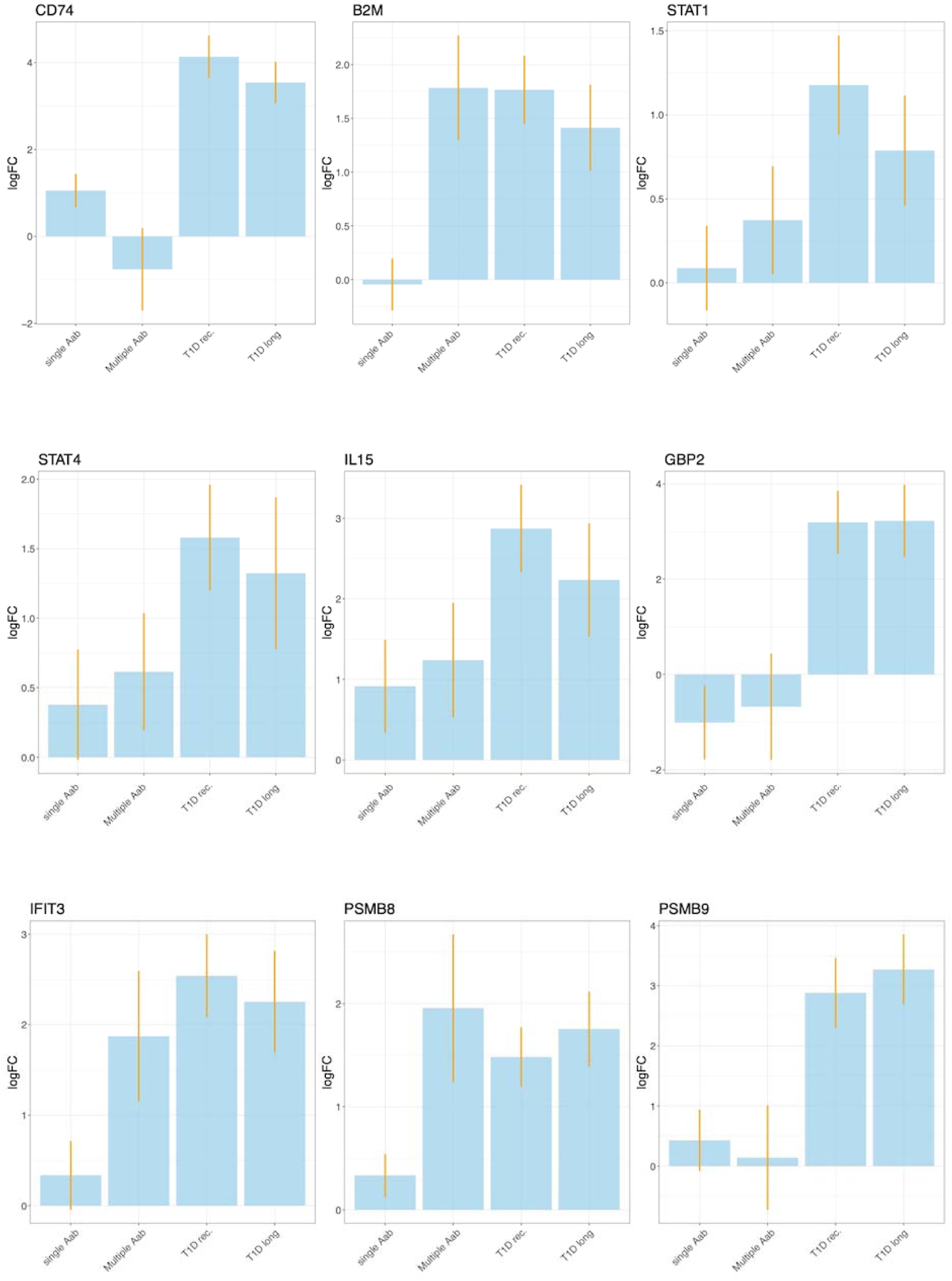
Barplot of Iog2fc for each diabetes status relative to non-diabetic for a subset of differential expressed genes in beta cells

**Supplementary Figure 9:**
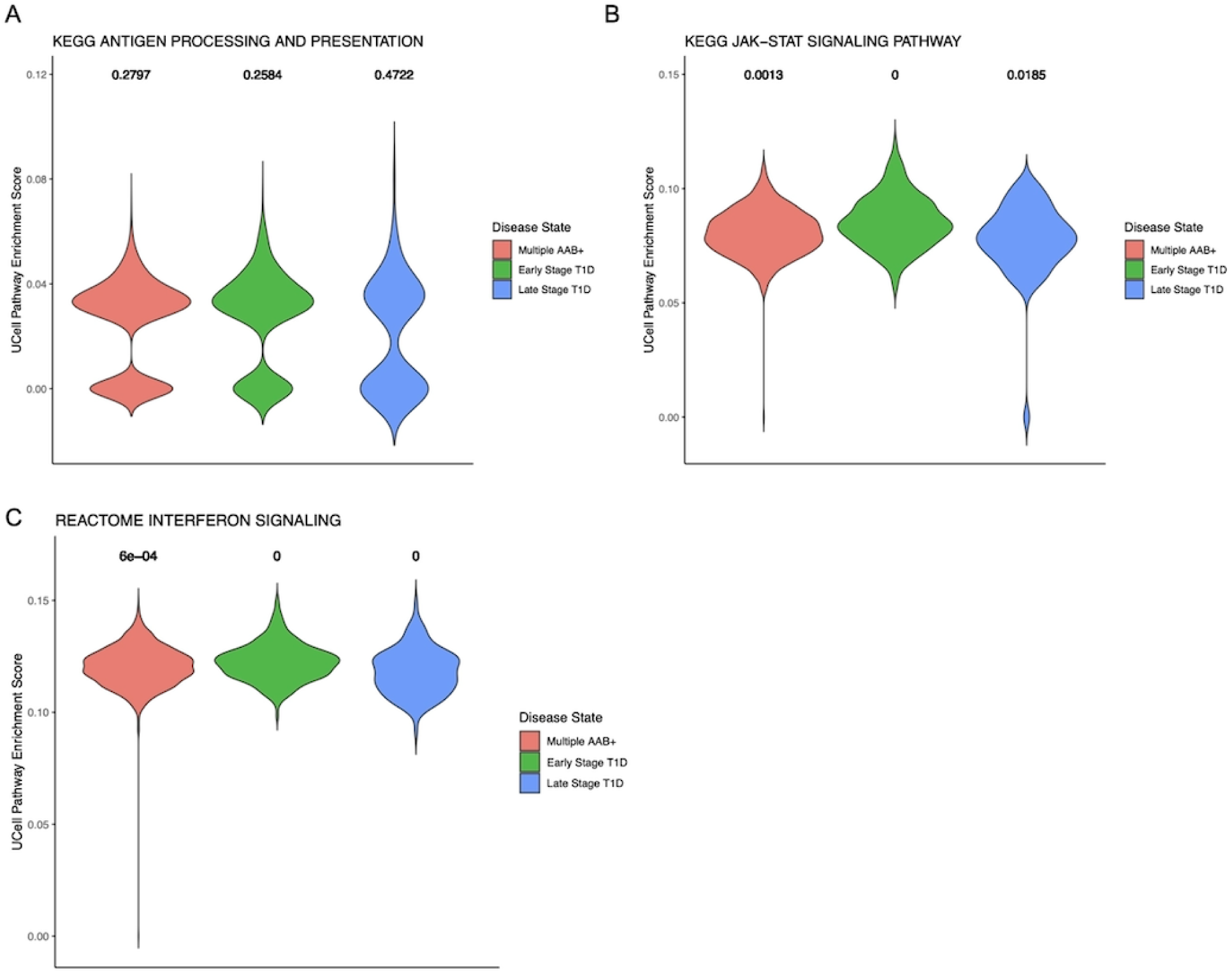
Pathways up-regulated in recent onset T1D relative to non-diabetic controls exhibiting **A)** heterogenous pathway enrichment and **B-C)** a continuous distribution of UCell scores

**Supplementary Figure 10:**
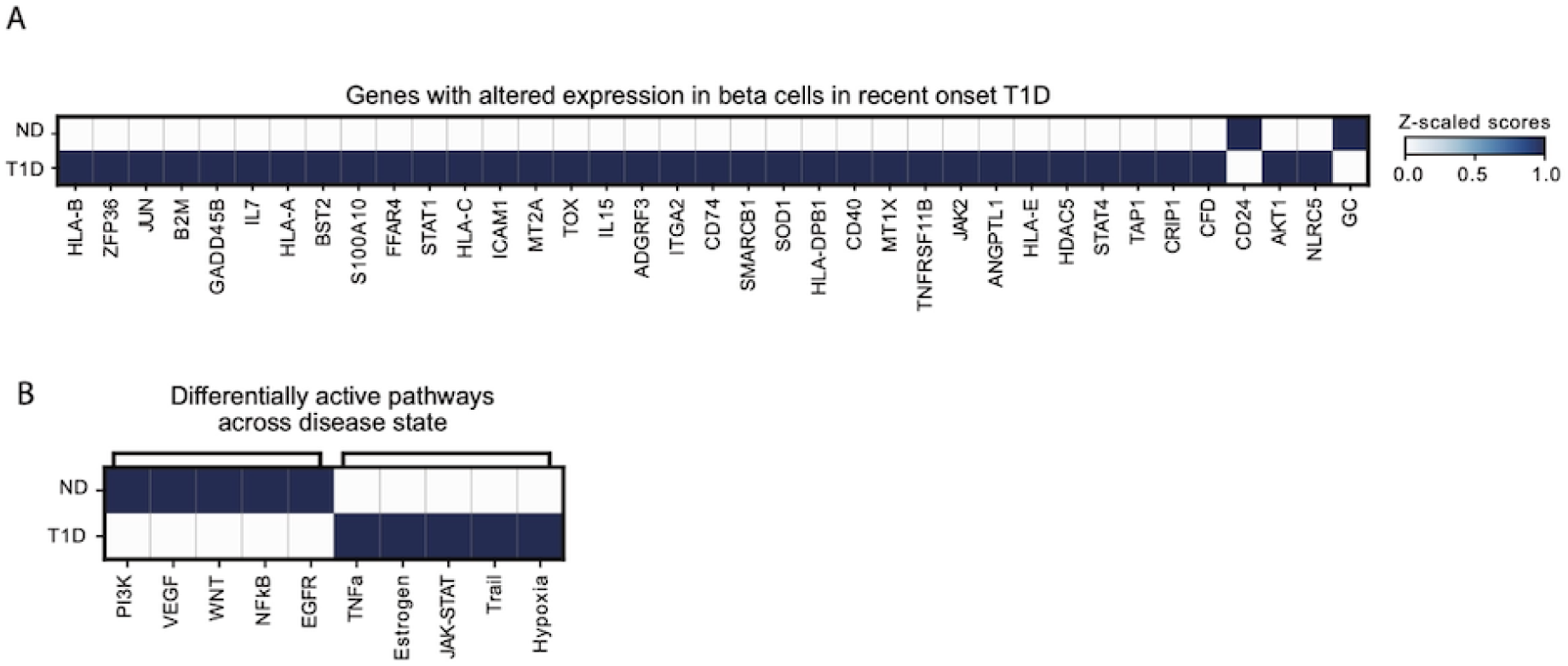
A. Matrix plot showing scaled genes with altered expression in beta cells in recent onset T1D and their pattern on the spatial gene expression across conditions. B. Matrix plot showing the scaled pathway activity of the differentially active pathways across disease state computed using progeny.

**Supplementary Figure 11:**
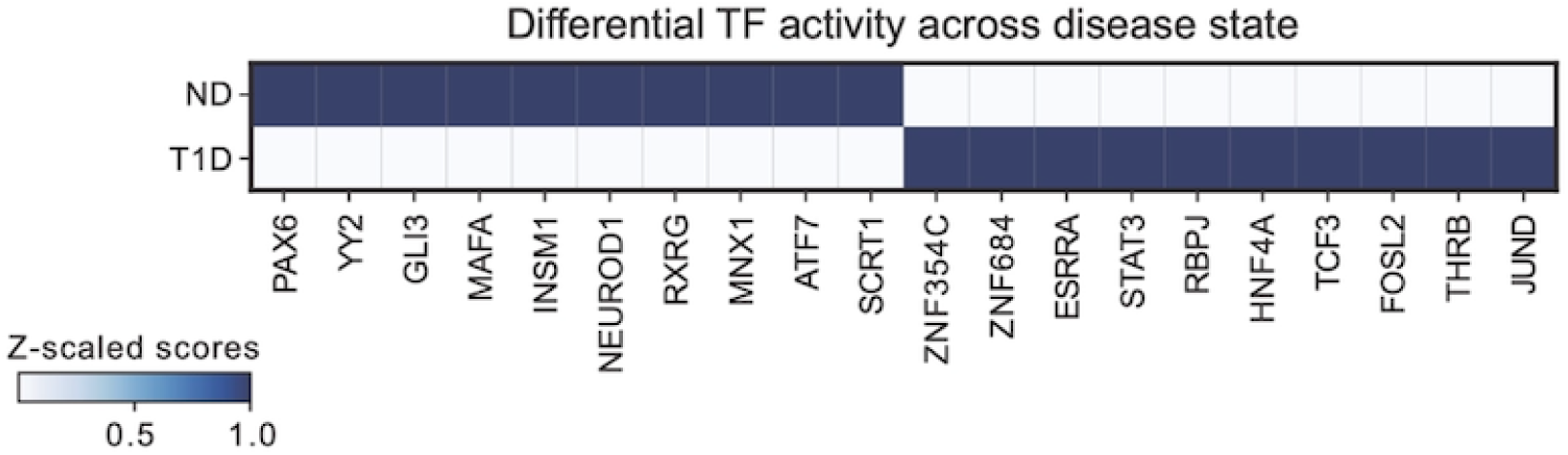
Matrix plot showing the scaled TF activity of the differentially active TFs across disease state.

**Supplemental Figure 12:**
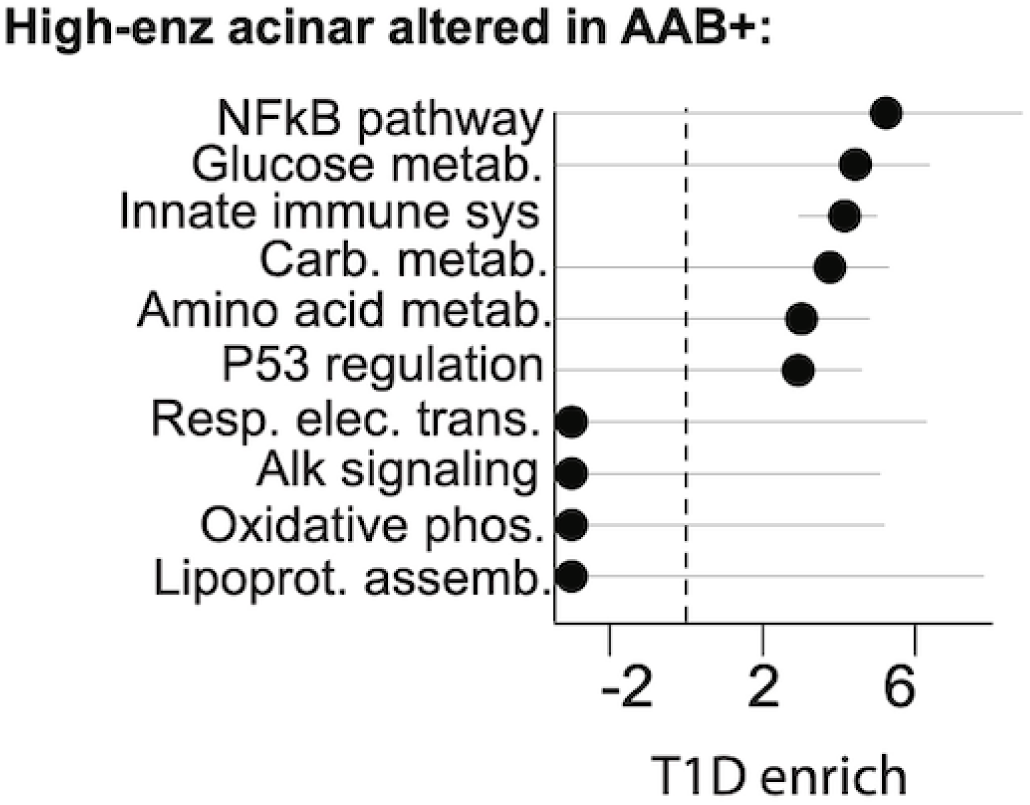
Enrichment of T1D risk loci in pathways altered in T1D Aab+ donor in high enz. acinar population.

**Supplemental Figure 13:**
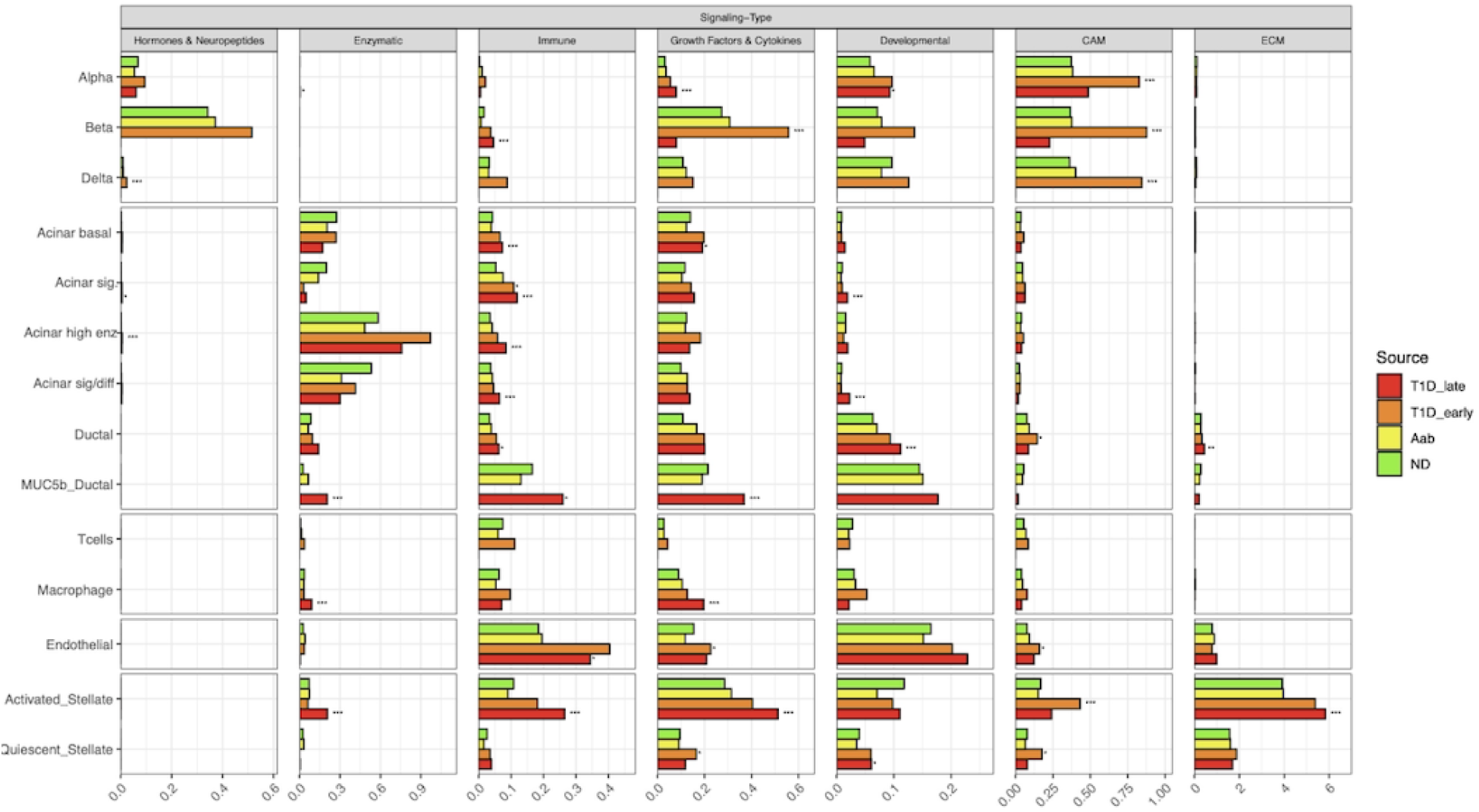
Cell to cell signaling during diabetes broken down by signaling category per cell type and diabetes status.

**Supplementary Figure 14:**
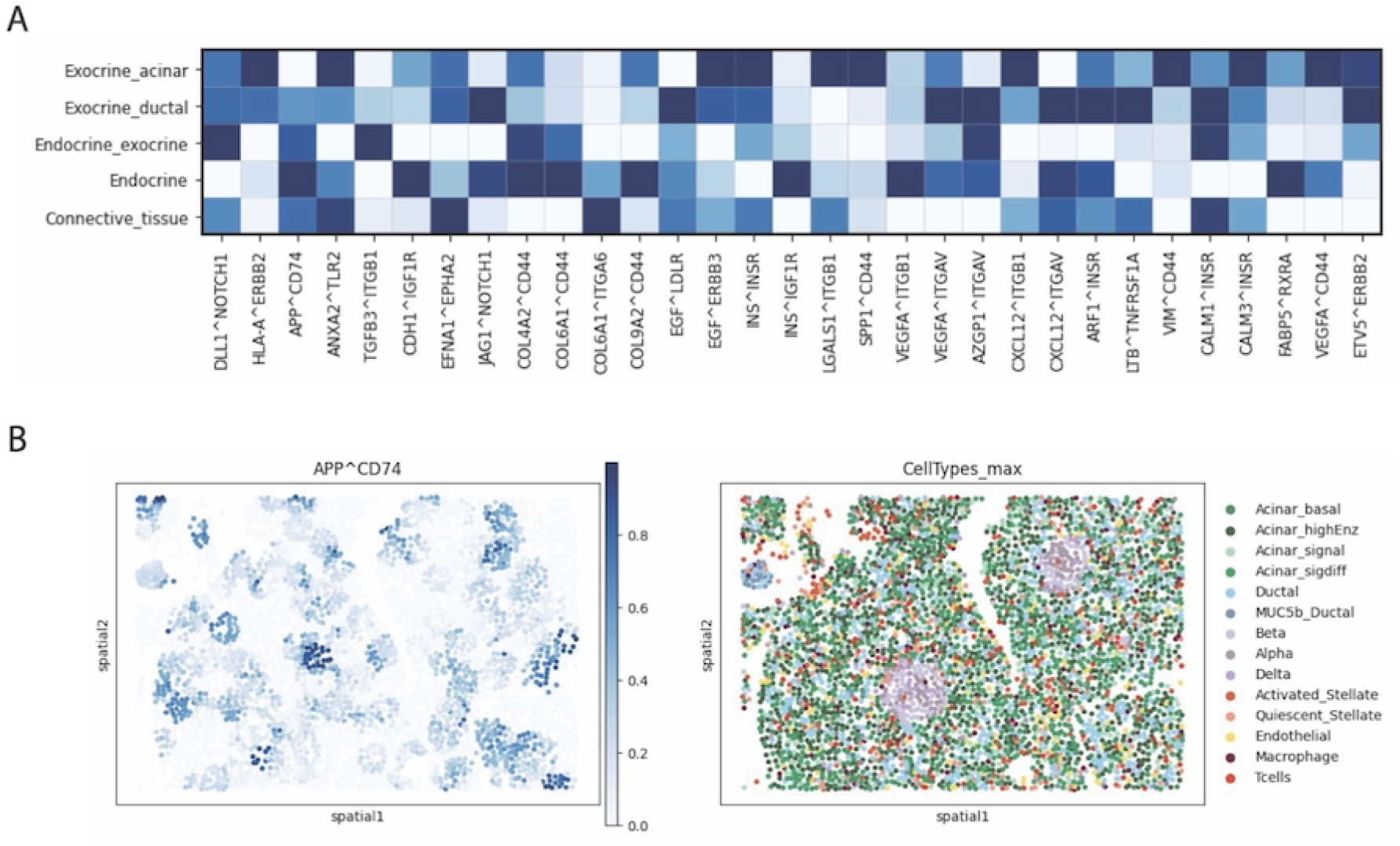
A. Matrix plot showing ligand-receptor scaled score computed using Liana+ across spatial niches B. Spatial plot showing the APP^A^CD74 score in fov 10 of sile 1 (T1D representative fov).

## References

1. Lehuen, A., Diana, J., Zaccone, P. & Cooke, A. Immune cell crosstalk in type 1 diabetes. Nat. Rev. Immunol. 10, 501–513 (2010).

2. Eguchi, K. & Nagai, R. Islet inflammation in type 2 diabetes and physiology Find the latest versionL: Islet inflammation in type 2 diabetes and physiology. J. Clin. Invest. 127, 14–23 (2017).

3. Boldison, J. & Wong, F. S. Immune and Pancreatic β Cell Interactions in Type 1 Diabetes. Trends Endocrinol. Metab. 27, 856–867 (2016).

4. Fasolino, M. et al. Single-cell multi-omics analysis of human pancreatic islets reveals novel cellular states in type 1 diabetes. Nat. Metab. 4, 284–299 (2022).

5. Chiou, J. et al. Interpreting type 1 diabetes risk with genetics and single-cell epigenomics. Nature 594, 398–402 (2021).

6. Mallone, R., Halliez, C., Rui, J. & Herold, K. C. The β-Cell in Type 1 Diabetes Pathogenesis: A Victim of Circumstances or an Instigator of Tragic Events? Diabetes 71, 1603–1610 (2022).

7. Atkinson, M. A. The pathogenesis and natural history of type 1 diabetes. Cold Spring Harb. Perspect. Med. 2, 1–18 (2012).

8. Pihoker, C., Gilliam, L. K., Hampe, C. S. & Lernmark, Å. Autoantibodies in Diabetes. 54, (2005).

9. Bingley, P. J. et al. Combined analysis of autoantibodies improves prediction of IDDM in islet cell antibody-positive relatives. Diabetes 43, 1304–1310 (1994).

10. Bingley, P. J. et al. Prediction of IDDM in the general population: strategies based on combinations of autoantibody markers. Diabetes 46, 1701–1710 (1997).

11. Fousteri, G., Ippolito, E., Ahmed, R. & Hamad, A. R. A. Beta-cell specific autoantibodies: Are they just an indicator of type 1 diabetes? Curr. Diabetes Rev. 13, 322–329 (2017).

12. Oram, R. A., Sims, E. K. & Evans-Molina, C. Beta cells in type 1 diabetes: mass and function; sleeping or dead? Diabetologia 62, 567–577 (2019).

13. Mănescu, M., Mănescu, I. B. & Grama, A. A Review of Stage 0 Biomarkers in Type 1 Diabetes: The Holy Grail of Early Detection and Prevention? J. Pers. Med. 14, 878 (2024).

14. Preissl, S., Gaulton, K. J. & Ren, B. Characterizing cis-regulatory elements using single-cell epigenomics. Nat. Rev. Genet. 24, 21–43 (2023).

15. Elgamal, R. M. et al. An Integrated Map of Cell Type–Specific Gene Expression in Pancreatic Islets. Diabetes 72, 1719–1728 (2023).

16. Bressan, D., Battistoni, G. & Hannon, G. J. The dawn of spatial omics. Science 381, eabq4964 (2023).

17. Pugliese, A. Insulitis in the pathogenesis of type 1 diabetes. Pediatr. Diabetes 17 **Suppl 22**, 31–36 (2016).

18. Chiou, J. et al. Interpreting type 1 diabetes risk with genetics and single-cell epigenomics. Nature 594, 398–402 (2021).

19. Benaglio, P. et al. Type 1 diabetes risk genes mediate pancreatic beta cell survival in response to proinflammatory cytokines. Cell Genomics 2, 100214 (2022).

20. Bingley, P. J., Boulware, D. C. & Krischer, J. P. The implications of autoantibodies to a single islet antigen in relatives with normal glucose tolerance: development of other autoantibodies and progression to type 1 diabetes. Diabetologia 59, 542–549 (2016).

21. Foster, T. P. et al. Exocrine Pancreas Dysfunction in Type 1 Diabetes. Endocr. Pract. Off. J. Am. Coll. Endocrinol. Am. Assoc. Clin. Endocrinol. 26, 1505–1513 (2020).

22. Korsunsky, I. et al. Fast, sensitive and accurate integration of single-cell data with Harmony. Nat. Methods 16, 1289–1296 (2019).

23. Klein, D. et al. Mapping cells through time and space with moscot. Nature (2025) doi:10.1038/s41586-024-08453-2.

24. Palla, G. et al. Squidpy: a scalable framework for spatial omics analysis. Nat. Methods 19, 171–178 (2022).

25. Haviv, D., Gatie, M., Hadjantonakis, A.-K., Nawy, T. & Pe’er, D. The covariance environment defines cellular niches for spatial inference. BioRxiv Prepr. Serv. Biol. 2023.04.18.537375 (2023) doi:10.1101/2023.04.18.537375.

26. Ali, M. et al. GraphCompass: Spatial metrics for differential analyses of cell organization across conditions. Preprint at 10.1101/2024.02.02.578605 (2024).

27. Stuart, T. et al. Comprehensive Integration of Single-Cell Data. Cell 177, 1888–1902.e21 (2019).

28. Schep, A. N., Wu, B., Buenrostro, J. D. & Greenleaf, W. J. chromVAR: inferring transcription-factor-associated accessibility from single-cell epigenomic data. Nat. Methods 14, 975–978 (2017).

29. Zhang, K. et al. A single-cell atlas of chromatin accessibility in the human genome. Cell 184, 5985–6001.e19 (2021).

30. Badia-i-Mompel, P., et al. decoupleR: ensemble of computational methods to infer biological activities from omics data. Bioinforma. Adv. 2, vbac016 (2022).

31. Mitchell, R. K. et al. The transcription factor Pax6 is required for pancreatic β cell identity, glucose-regulated ATP synthesis, and Ca2+ dynamics in adult mice. J. Biol. Chem. 292, 8892–8906 (2017).

32. Jiang, M. et al. MIST1 and PTF1 Collaborate in Feed-Forward Regulatory Loops That Maintain the Pancreatic Acinar Phenotype in Adult Mice. Mol. Cell. Biol. 36, 2945–2955 (2016).

33. Elgamal, R. M. et al. An Integrated Map of Cell Type–Specific Gene Expression in Pancreatic Islets. Diabetes 72, 1719–1728 (2023).

34. Kaestner, K. H., Powers, A. C., Naji, A., HPAP Consortium & Atkinson, M. A. NIH Initiative to Improve Understanding of the Pancreas, Islet, and Autoimmunity in Type 1 Diabetes: The Human Pancreas Analysis Program (HPAP). Diabetes 68, 1394–1402 (2019).

35. Dimitrov, D. et al. LIANA+ provides an all-in-one framework for cell–cell communication inference. Nat. Cell Biol. 26, 1613–1622 (2024).

36. Matsui, J., Wakabayashi, T., Asada, M., Yoshimatsu, K. & Okada, M. Stem cell factor/c-kit signaling promotes the survival, migration, and capillary tube formation of human umbilical vein endothelial cells. J. Biol. Chem. 279, 18600–18607 (2004).

37. Wang, C.-H. et al. Stem cell factor attenuates vascular smooth muscle apoptosis and increases intimal hyperplasia after vascular injury. Arterioscler. Thromb. Vasc. Biol. 27, 540–547 (2007).

38. Wang, G. et al. Integrating genetics with single-cell multiomic measurements across disease states identifies mechanisms of beta cell dysfunction in type 2 diabetes. Nat. Genet. 55, 984–994 (2023).

39. Pickrell, J. K. Joint analysis of functional genomic data and genome-wide association studies of 18 human traits. Am. J. Hum. Genet. 94, 559–573 (2014).

40. Jin, S. et al. Inference and analysis of cell-cell communication using CellChat. Nat. Commun. 12, 1088 (2021).

41. Li, Z., Wang, T., Liu, P. & Huang, Y. SpatialDM for rapid identification of spatially co-expressed ligand–receptor and revealing cell–cell communication patterns. Nat. Commun. 14, 3995 (2023).

42. Korf, H. et al. MIF inhibition interferes with the inflammatory and T cell-stimulatory capacity of NOD macrophages and delays autoimmune diabetes onset. PloS One 12, e0187455 (2017).

43. Dooley, J. et al. Genetic predisposition for beta cell fragility underlies type 1 and type 2 diabetes. Nat. Genet. 48, 519–527 (2016).

44. Liston, A., Todd, J. A. & Lagou, V. Beta-Cell Fragility As a Common Underlying Risk Factor in Type 1 and Type 2 Diabetes. Trends Mol. Med. 23, 181–194 (2017).

45. Marroquí, L. et al. BACH2, a candidate risk gene for type 1 diabetes, regulates apoptosis in pancreatic β-cells via JNK1 modulation and crosstalk with the candidate gene PTPN2. Diabetes 63, 2516–2527 (2014).

46. Santin, I. & Eizirik, D. L. Candidate genes for type 1 diabetes modulate pancreatic islet inflammation and β-cell apoptosis. Diabetes Obes. Metab. 15 **Suppl 3**, 71–81 (2013).

47. Santin, I. et al. PTPN2, a candidate gene for type 1 diabetes, modulates pancreatic β-cell apoptosis via regulation of the BH3-only protein Bim. Diabetes 60, 3279–3288 (2011).

48. Størling, J. & Pociot, F. Type 1 Diabetes Candidate Genes Linked to Pancreatic Islet Cell Inflammation and Beta-Cell Apoptosis. Genes 8, 72 (2017).

49. Colli, M. L., Moore, F., Gurzov, E. N., Ortis, F. & Eizirik, D. L. MDA5 and PTPN2, two candidate genes for type 1 diabetes, modify pancreatic beta-cell responses to the viral by- product double-stranded RNA. Hum. Mol. Genet. 19, 135–146 (2010).

50. Tsan, M.-F. & Gao, B. Heat shock proteins and immune system. J. Leukoc. Biol. 85, 905– 910 (2009).

51. Kregel, K. C. Invited Review: Heat shock proteins: modifying factors in physiological stress responses and acquired thermotolerance. J. Appl. Physiol. 92, 2177–2186 (2002).

52. Bogdani, M. et al. Extracellular matrix components in the pathogenesis of type 1 diabetes. Curr. Diab. Rep. 14, 552 (2014).

53. Doliba, N. M. et al. α Cell dysfunction in islets from nondiabetic, glutamic acid decarboxylase autoantibody-positive individuals. J. Clin. Invest. 132, e156243 (2022).

54. Tosti, L. et al. Single-Nucleus and In Situ RNA–Sequencing Reveal Cell Topographies in the Human Pancreas. Gastroenterology 160, 1330–1344.e11 (2021).

55. Chiou, J. et al. Single-cell chromatin accessibility identifies pancreatic islet cell type- and state-specific regulatory programs of diabetes risk. Nat. Genet. 53, 455–466 (2021).

56. Thomas, H. E., Trapani, J. A. & Kay, T. W. H. The role of perforin and granzymes in diabetes. Cell Death Differ. 17, 577–585 (2010).

57. Notkins, A. L. Immunologic and Genetic Factors in Type 1 Diabetes. J. Biol. Chem. 277, 43545–43548 (2002).

58. Goulley, J., Dahl, U., Baeza, N., Mishina, Y. & Edlund, H. BMP4-BMPR1A signaling in beta cells is required for and augments glucose-stimulated insulin secretion. Cell Metab. 5, 207– 219 (2007).

59. Chmielowiec, J. et al. Human pancreatic microenvironment promotes β-cell differentiation via non-canonical WNT5A/JNK and BMP signaling. Nat. Commun. 13, 1952 (2022).

60. Urizar, A. I. et al. Beta cell dysfunction induced by bone morphogenetic protein (BMP)-2 is associated with histone modifications and decreased NeuroD1 chromatin binding. Cell Death Dis. 14, 399 (2023).

61. Barbu, A., Lejonklou, M. H. & Skogseid, B. Progranulin Stimulates Proliferation of Mouse Pancreatic Islet Cells and Is Overexpressed in the Endocrine Pancreatic Tissue of an MEN1 Mouse Model. Pancreas 45, 533–540 (2016).

62. Cheung, P. F. et al. Progranulin mediates immune evasion of pancreatic ductal adenocarcinoma through regulation of MHCI expression. Nat. Commun. 13, 156 (2022).

63. Insel, R. A. et al. Staging Presymptomatic Type 1 Diabetes: A Scientific Statement of JDRF, the Endocrine Society, and the American Diabetes Association. Diabetes Care 38, 1964– 1974 (2015).

64. Battaglia, M. et al. Introducing the Endotype Concept to Address the Challenge of Disease Heterogeneity in Type 1 Diabetes. Diabetes Care 43, 5–12 (2020).

65. Redondo, M. J. & Morgan, N. G. Heterogeneity and endotypes in type 1 diabetes mellitus. Nat. Rev. Endocrinol. 19, 542–554 (2023).

66. McGrail, C. et al. Genetic discovery and risk prediction for type 1 diabetes in individuals without high-risk HLA-DR3/DR4 haplotypes. Preprint at 10.1101/2023.11.11.23298405 (2023).

67. McGrail, C., Sears, T. J., Kudtarkar, P., Carter, H. & Gaulton, K. Genetic association and machine learning improves discovery and prediction of type 1 diabetes. MedRxiv Prepr. Serv. Health Sci. 2024.07.31.24311310 (2024) doi:10.1101/2024.07.31.24311310.

68. Abedi, M. et al. Aberrant TNF signaling in pancreatic lymph nodes of patients with Type 1 Diabetes. Preprint at 10.1101/2024.05.31.596885 (2024).

69. Chiou, J. et al. Single-cell chromatin accessibility identifies pancreatic islet cell type– and state-specific regulatory programs of diabetes risk. Nat. Genet. 53, 455–466 (2021).

70. Thibodeau, A. et al. AMULET: a novel read count-based method for effective multiplet detection from single nucleus ATAC-seq data. Genome Biol. 22, 252 (2021).

71. Zheng, G. X. Y. et al. Massively parallel digital transcriptional profiling of single cells. Nat. Commun. 8, 14049 (2017).

72. McGinnis, C. S., Murrow, L. M. & Gartner, Z. J. DoubletFinder: Doublet Detection in Single-Cell RNA Sequencing Data Using Artificial Nearest Neighbors. Cell Syst. 8, 329–337.e4 (2019).

73. Young, M. D. & Behjati, S. SoupX removes ambient RNA contamination from droplet-based single-cell RNA sequencing data. GigaScience 9, giaa151 (2020).

74. Hao, Y. et al. Integrated analysis of multimodal single-cell data. Cell 184, 3573–3587.e29 (2021).

75. Korsunsky, I. et al. Fast, sensitive and accurate integration of single-cell data with Harmony. Nat. Methods 16, 1289–1296 (2019).

76. Love, M. I., Huber, W. & Anders, S. Moderated estimation of fold change and dispersion for RNA-seq data with DESeq2. Genome Biol. 15, 1–21 (2014).

77. Kanehisa, M. & Goto, S. KEGG: Kyoto Encyclopedia of Genes and Genomes. Nucleic Acids Res. 28, 27–30 (2000).

78. Kanehisa, M. Toward understanding the origin and evolution of cellular organisms. Protein Sci. Publ. Protein Soc. 28, 1947–1951 (2019).

79. Kanehisa, M., Furumichi, M., Sato, Y., Kawashima, M. & Ishiguro-Watanabe, M. KEGG for taxonomy-based analysis of pathways and genomes. Nucleic Acids Res. 51, D587–D592 (2022).

80. Milacic, M. et al. The Reactome Pathway Knowledgebase 2024. Nucleic Acids Res. 52, D672–D678 (2024).

81. Korotkevich, G. et al. Fast gene set enrichment analysis. 060012 Preprint at 10.1101/060012 (2021).

82. Stuart, T., Srivastava, A., Lareau, C. & Satija, R. Multimodal single-cell chromatin analysis with Signac. bioRxiv 1–17 (2020) doi:10.1101/2020.11.09.373613.

83. Zhang, Y. et al. Model-based Analysis of ChIP-Seq (MACS). Genome Biol. 9, R137 (2008).

84. Dries, R. et al. Giotto: a toolbox for integrative analysis and visualization of spatial expression data. Genome Biol. 22, 78 (2021).

85. Wolf, F. A., Angerer, P. & Theis, F. J. SCANPY: large-scale single-cell gene expression data analysis. Genome Biol. 19, 15 (2018).

86. Lopez, R., Regier, J., Cole, M. B., Jordan, M. I. & Yosef, N. Deep generative modeling for single-cell transcriptomics. Nat. Methods 15, 1053–1058 (2018).

87. Klein, D. et al. Mapping cells through time and space with moscot. Preprint at 10.1101/2023.05.11.540374 (2023).

88. Nassar, L. R. et al. The UCSC Genome Browser database: 2023 update. Nucleic Acids Res. 51, D1188–D1195 (2023).

89. Heinz, S. et al. Simple combinations of lineage-determining transcription factors prime cis-regulatory elements required for macrophage and B cell identities. Mol. Cell 38, 576–589 (2010).

90. McLean, C. Y. et al. GREAT improves functional interpretation of cis-regulatory regions. Nat. Biotechnol. 28, 495–501 (2010).

91. Frankish, A. et al. GENCODE: reference annotation for the human and mouse genomes in 2023. Nucleic Acids Res. 51, D942–D949 (2023).

92. Hothorn, T., Zeileis, A., Farebrother, R. W. & Cummins, C. lmtest: Testing Linear Regression Models. 0.9-40 10.32614/CRAN.package.lmtest (1999).

93. Kaestner, K. H., Powers, A. C., Naji, A., HPAP Consortium & Atkinson, M. A. NIH Initiative to Improve Understanding of the Pancreas, Islet, and Autoimmunity in Type 1 Diabetes: The Human Pancreas Analysis Program (HPAP). Diabetes 68, 1394–1402 (2019).

94. Bates, D., Mächler, M., Bolker, B. & Walker, S. Fitting Linear Mixed-Effects Models Using **lme4**. J. Stat. Softw. 67, (2015).

95. Mootha, V. K. et al. PGC-1α-responsive genes involved in oxidative phosphorylation are coordinately downregulated in human diabetes. Nat. Genet. 34, 267–273 (2003).

96. Subramanian, A. et al. Gene set enrichment analysis: A knowledge-based approach for interpreting genome-wide expression profiles. Proc. Natl. Acad. Sci. 102, 15545–15550 (2005).

97. Jassal, B. et al. The reactome pathway knowledgebase. Nucleic Acids Res. 48, D498–D503 (2020).

98. Griss, J. et al. ReactomeGSA - Efficient Multi-Omics Comparative Pathway Analysis. Mol. Cell. Proteomics MCP 19, 2115–2125 (2020).

99. Fabregat, A. et al. Reactome graph database: Efficient access to complex pathway data. PLoS Comput. Biol. 14, e1005968 (2018).

100. Fabregat, A. et al. Reactome diagram viewer: data structures and strategies to boost performance. Bioinforma. Oxf. Engl. 34, 1208–1214 (2018).

101. Sidiropoulos, K. et al. Reactome enhanced pathway visualization. Bioinforma. Oxf. Engl. 33, 3461–3467 (2017).

102. Wu, G. & Haw, R. Functional Interaction Network Construction and Analysis for Disease Discovery. Methods Mol. Biol. Clifton NJ 1558, 235–253 (2017).

103. Gillespie, M. et al. The reactome pathway knowledgebase 2022. Nucleic Acids Res. 50, D687–D692 (2022).

104. John D. Storey With Contributions From Andrew J. Bass, A. qvalue. Bioconductor 10.18129/B9.BIOC.QVALUE (2017).

105. Kuznetsova, A., Brockhoff, P. B. & Christensen, R. H. B. **lmerTest** Package: Tests in Linear Mixed Effects Models. J. Stat. Softw. 82, (2017).

106. Duttke, S. H., Chang, M. W., Heinz, S. & Benner, C. Identification and dynamic quantification of regulatory elements using total RNA. Genome Res. 29, 1836–1846 (2019).

107. Fulco, C. P. et al. Activity-by-contact model of enhancer–promoter regulation from thousands of CRISPR perturbations. Nat. Genet. 51, 1664–1669 (2019).

108. Zhao, H. et al. CrossMap: a versatile tool for coordinate conversion between genome assemblies. Bioinformatics 30, 1006–1007 (2014).

109. Arda, H. E. et al. A Chromatin Basis for Cell Lineage and Disease Risk in the Human Pancreas. Cell Syst. 7, 310–322.e4 (2018).

110. Castro-Mondragon, J. A. et al. JASPAR 2022: the 9th release of the open-access database of transcription factor binding profiles. Nucleic Acids Res. 50, D165–D173 (2022).

111. Satpathy, A. T. et al. Massively parallel single-cell chromatin landscapes of human immune cell development and intratumoral T cell exhaustion. Nat. Biotechnol. 37, 925–936 (2019).

112. Schep, A. motifmatchr: Fast Motif Matching in R.

113. Nasser, J. et al. Genome-wide enhancer maps link risk variants to disease genes. Nature 593, 238–243 (2021).

114. Schep, A. N., Wu, B., Buenrostro, J. D. & Greenleaf, W. J. chromVAR: inferring transcription-factor-associated accessibility from single-cell epigenomic data. Nat. Methods 14, 975–978 (2017).

115. Wakefield, J. Bayes factors for genome-wide association studies: comparison with P-values. Genet. Epidemiol. 33, 79–86 (2009).

116. Aylward, A. et al. Glucocorticoid signaling in pancreatic islets modulates gene regulatory programs and genetic risk of type 2 diabetes. PLoS Genet. 17, e1009531 (2021).

117. UniProt: the Universal Protein Knowledgebase in 2023 | Nucleic Acids Research | Oxford Academic. https://academic.oup.com/nar/article/51/D1/D523/6835362?login=true.

118. Stelzer, G. et al. The GeneCards Suite: From Gene Data Mining to Disease Genome Sequence Analyses. Curr. Protoc. Bioinforma. 54, 1.30.1-1.30.33 (2016).

119. Safran, M. et al. The GeneCards Suite. in Practical Guide to Life Science Databases (eds. Abugessaisa, I. & Kasukawa, T.) 27–56 (Springer Nature, Singapore, 2021). doi:10.1007/978-981-16-5812-9_2.

120. Steffen Durinck <Biomartdev@Gmail. Com>, W. H. biomaRt. Bioconductor 10.18129/B9.BIOC.BIOMART (2017).

121. Andrews, S. No Title. FastQC: a quality control tool for high throughput sequence data. http://www.bioinformatics.babraham.ac.uk/projects/fastqc (2010).

122. Patro, R., Duggal, G., Love, M. I., Irizarry, R. A. & Kingsford, C. Salmon provides fast and bias-aware quantification of transcript expression. Nat. Methods 14, 417–419 (2017).

123. Soneson, C., Love, M. I. & Robinson, M. D. Differential analyses for RNA-seq: transcript-level estimates improve gene-level inferences. F1000Research 4, 1521 (2015).

