## Supplemental figures for "Single-cell multiome and spatial profiling reveals pancreas cell type-specific gene regulatory programs driving type 1 diabetes progression"

Supplementary figure 1

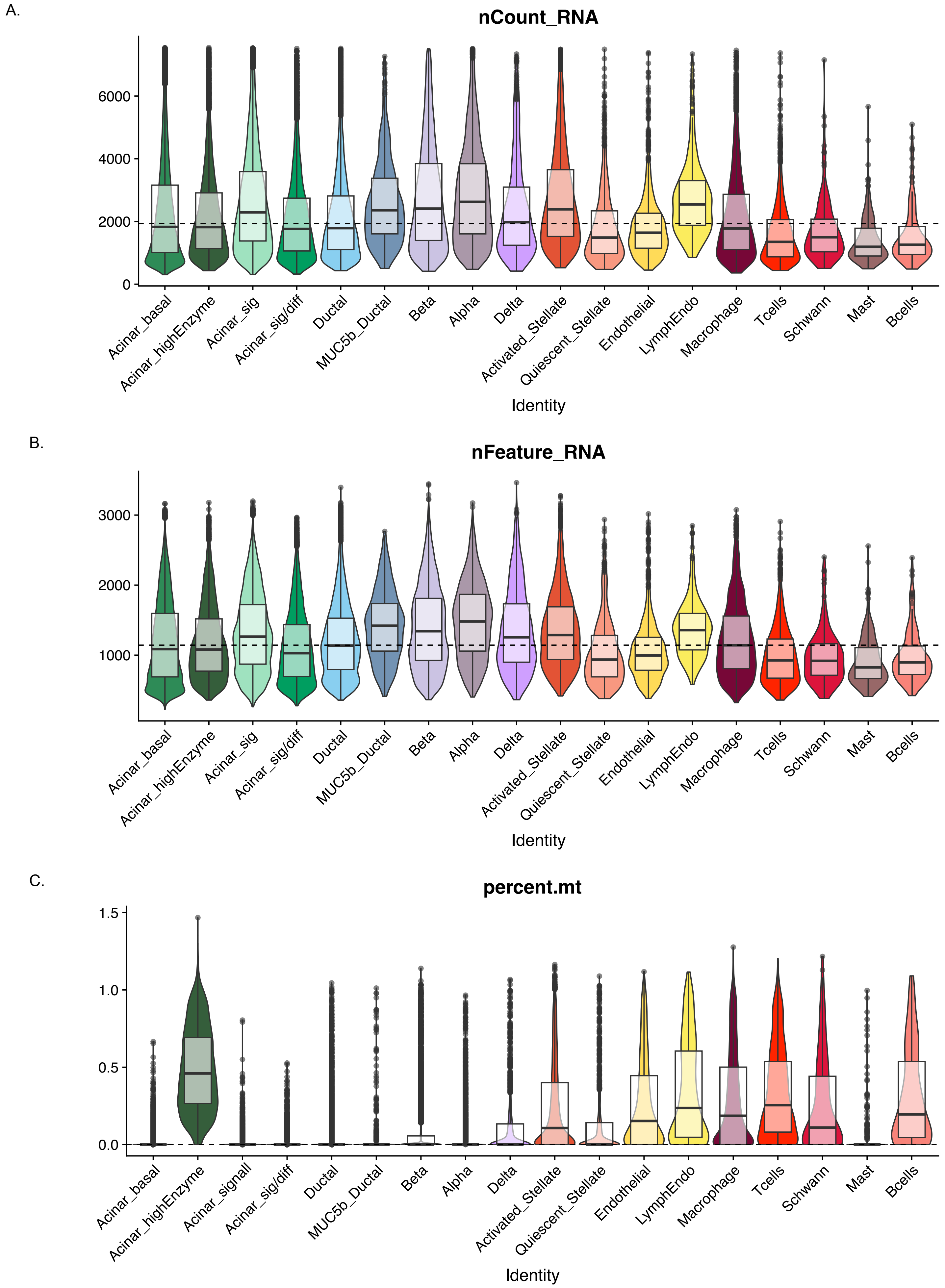


**Supplementary Figure 1**: Violin plots of quality control metrics for RNA expression data including (A) counts, (B) genes per nuclei, and (C) percent mitochondria

Supplementary figure 2

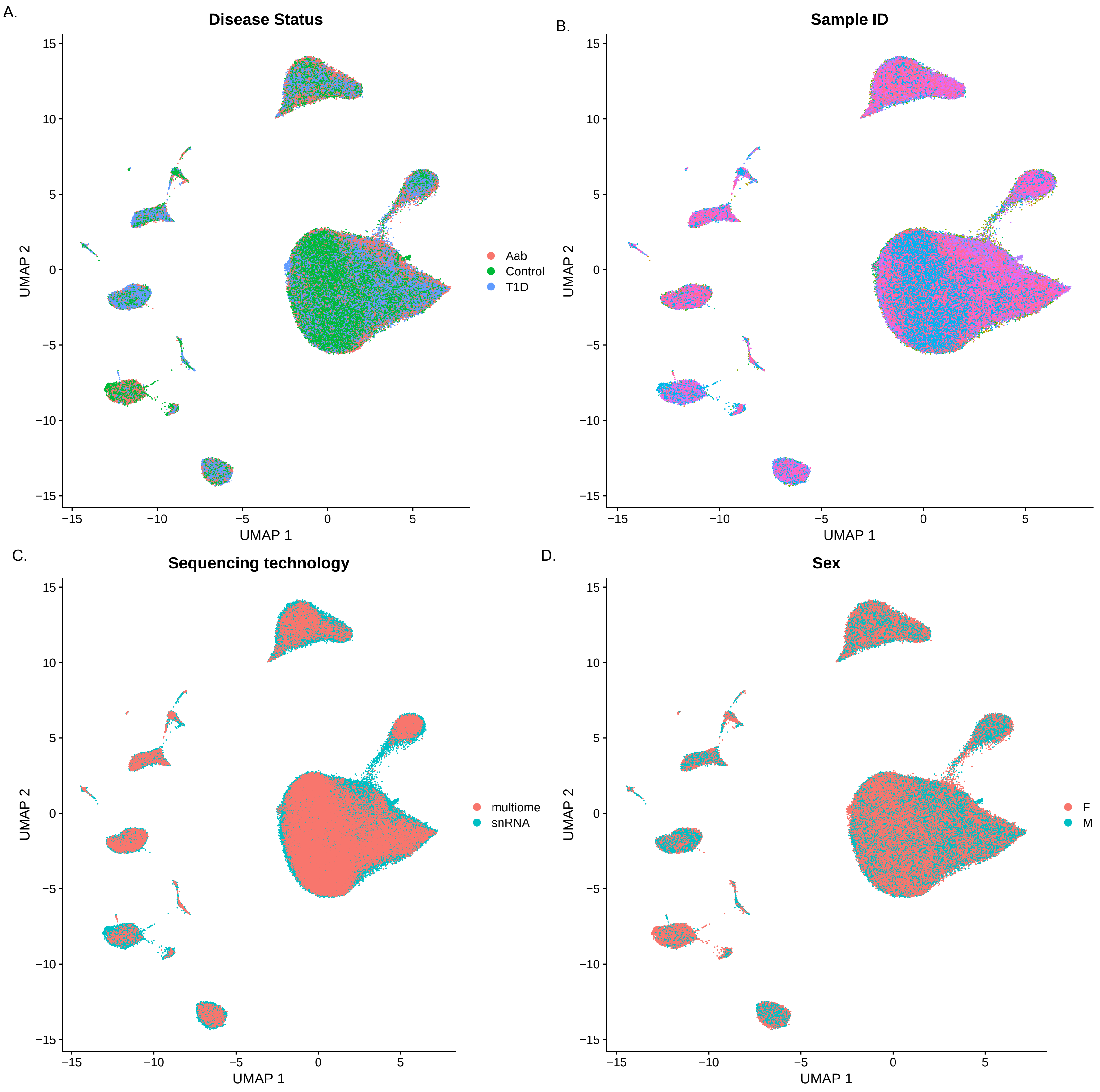


**Supplementary Figure 2:** Uniform manifold approximation and projection (UMAP) of gene expression data annotated by (A) diabetes status, (B) donor ID, (C) sequencing technology, and (D) sex.

Supplementary figure 3

A.

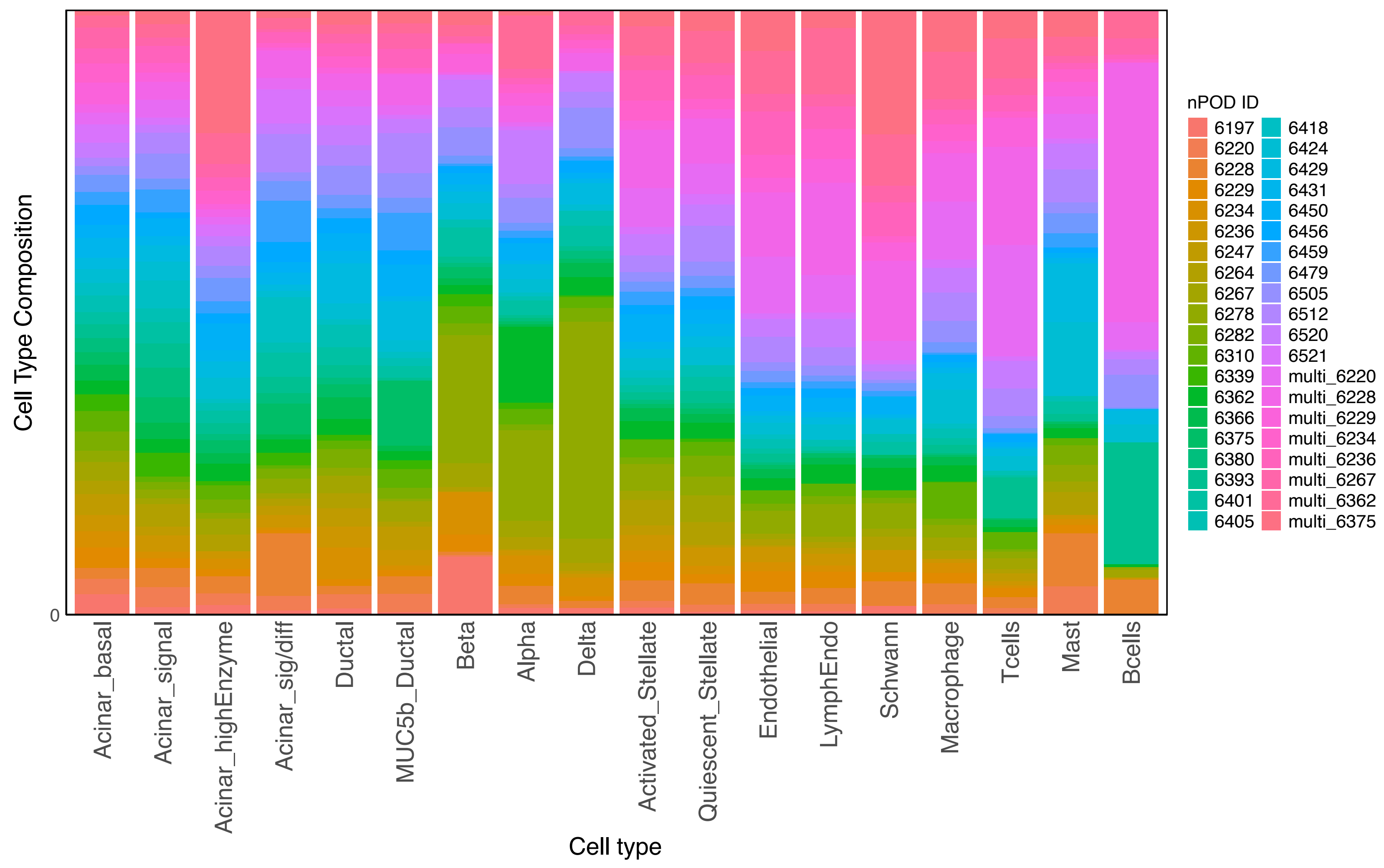


B.

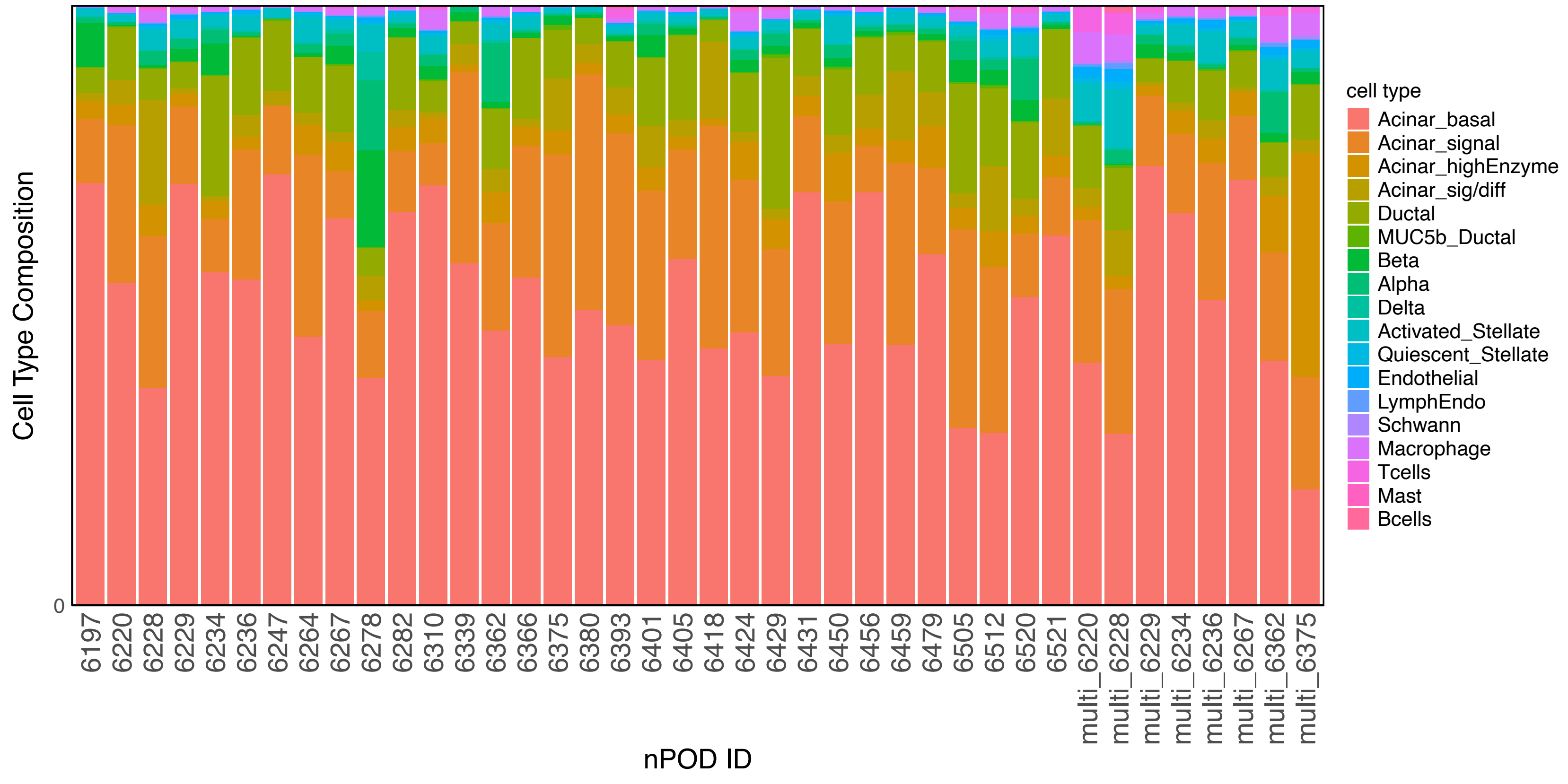


**Supplementary Figure 3:** Proportion of cells: (A) per cell type per donor, (B) per donor per cell type

Supplementary figure 4

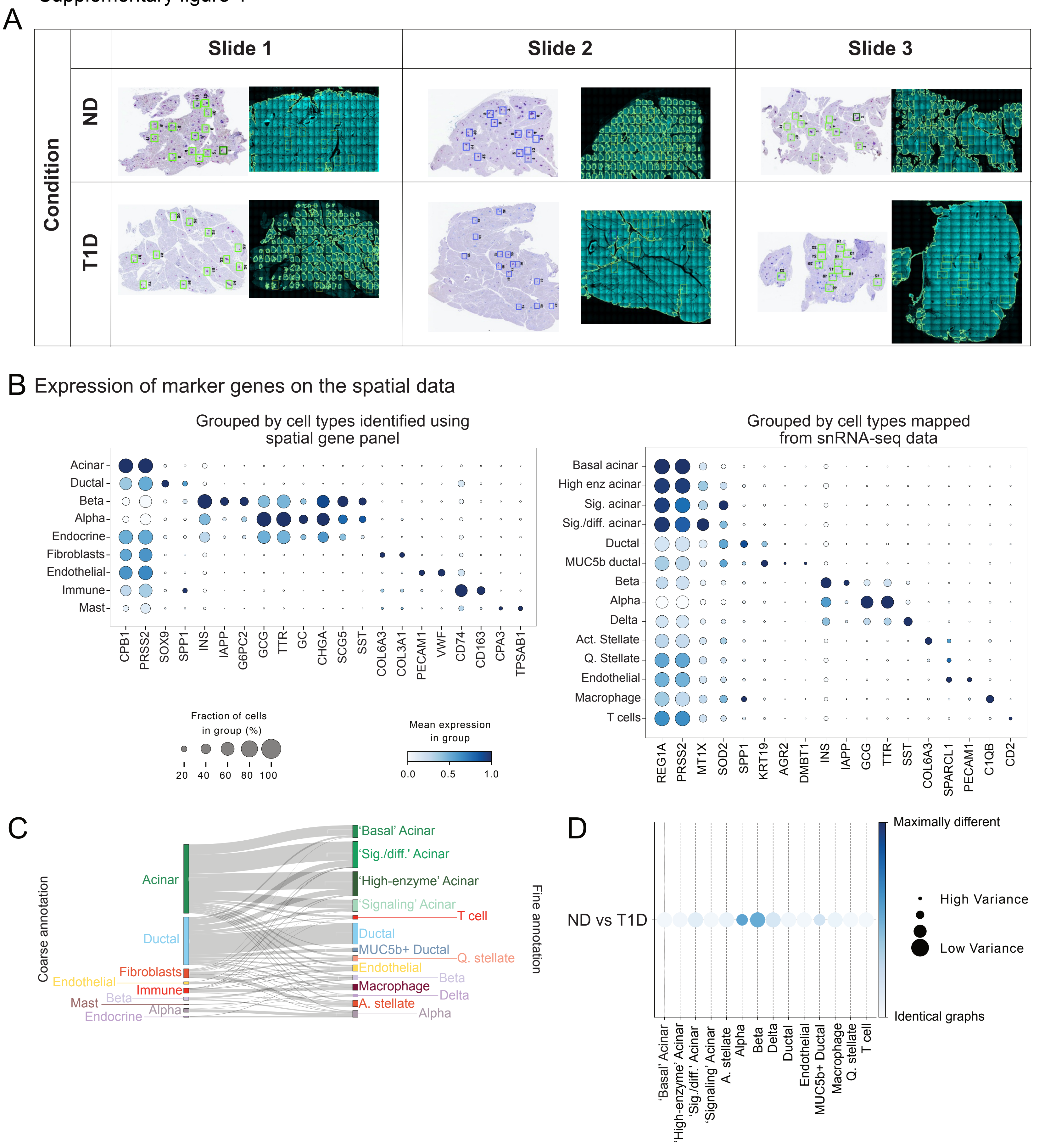


**Supplementary Figure 4**: A. Wole slide images of the sections profiled with CosMx. B. Dotplots of the cluster expression of canonical markers used for cell type annotations at different levels of granularity in the spatial slides. Coarse annotation (left), finer annotation (right). C. Sankey plot showing the relative mapping from coarse annotation to fine annotation. The mapping was done using optimal-transport-based method moscot. D. Cell-type-specific subgraph comparison across conditions (ND vs. T1D). The size of the dot is indicative of the dissimilarity score variance over samples. The larger the dot size, the lower the score variance and the higher the score confidence is.

Supplementary figure 5

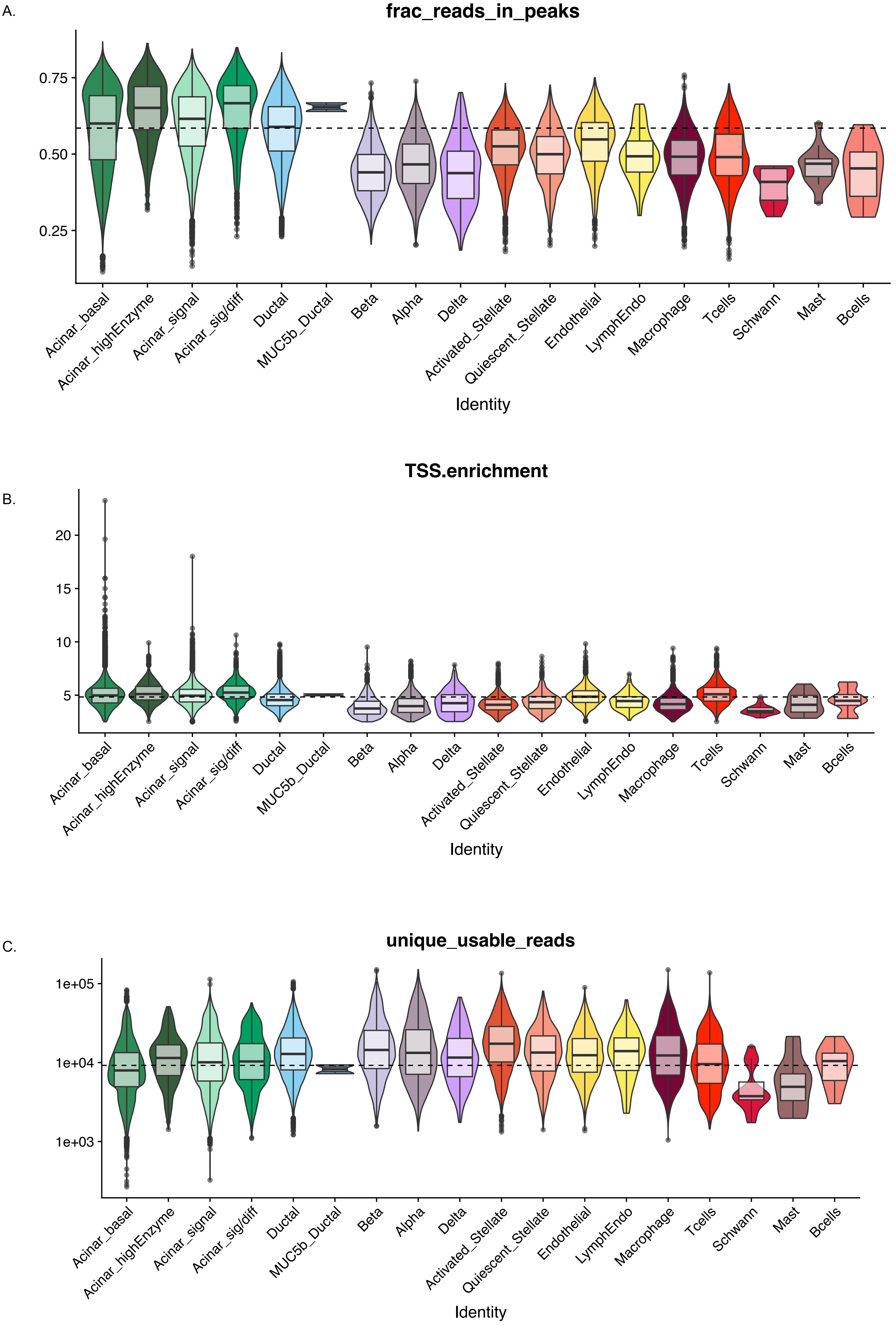


**Supplementary Figure 5:** Violin plots of quality control metrics for chromatin accessibility data including (A) fraction of reads in peaks, (B) transcription start site enrichment (TSSe), and (C) percent mitochondria

Supplementary figure 6

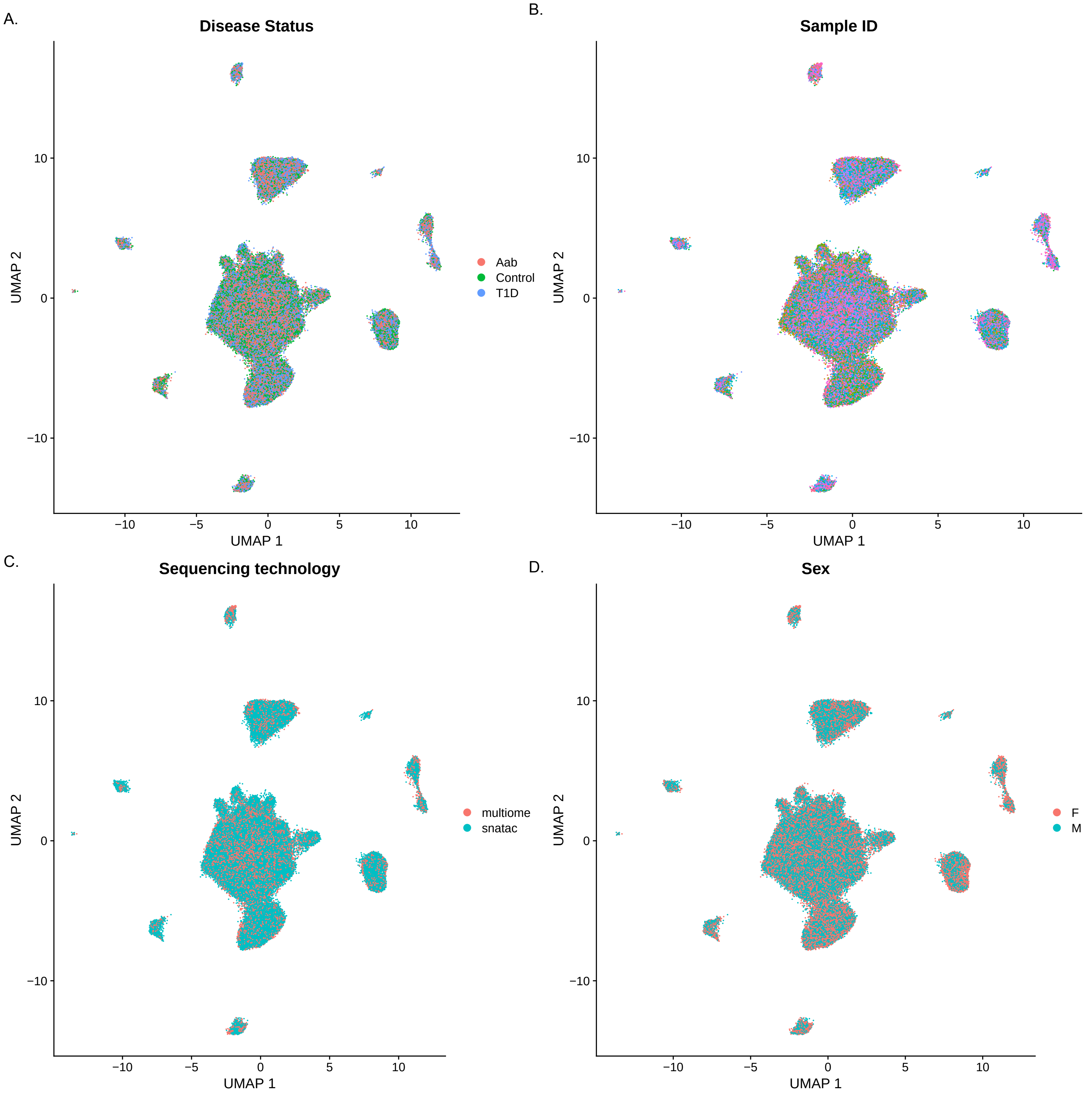


**Supplemental Figure 6:** Uniform manifold approximation and projection (UMAP) of chromatin accessibility data annotated by (A) diabetes status, (B) donor ID, (C) sequencing technology, and (D) sex

Supplementary figure 7

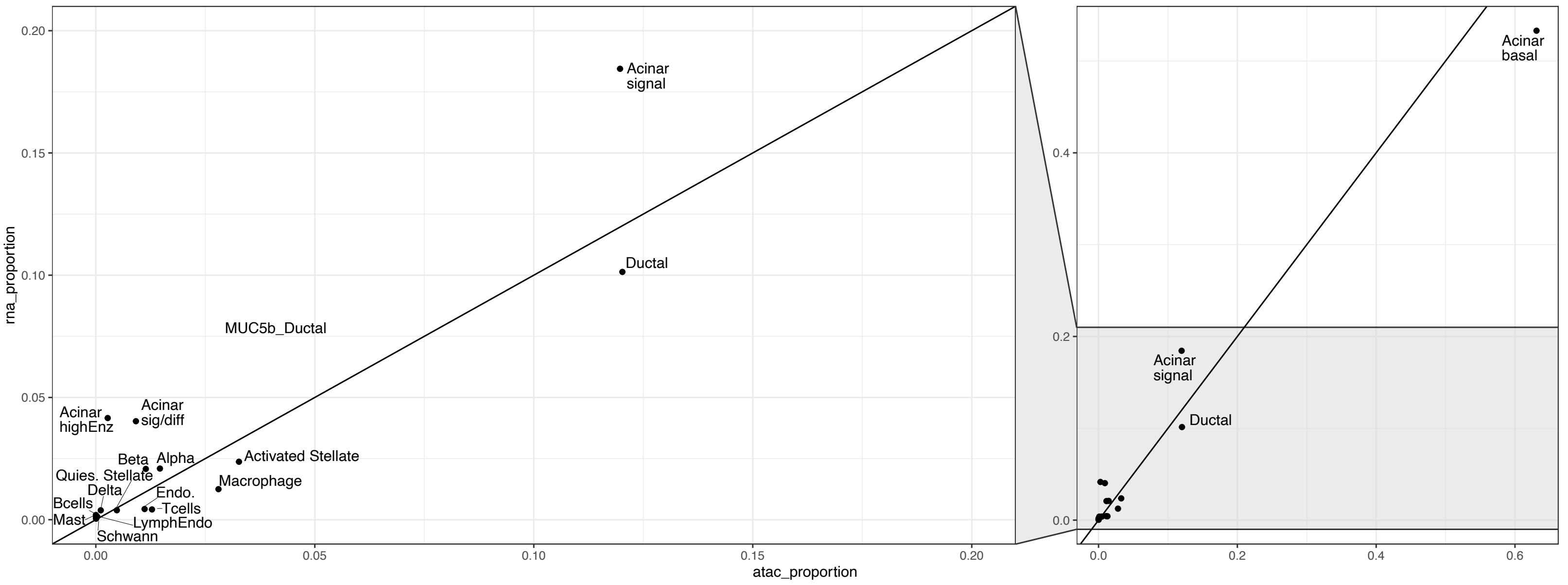


**Supplemental Figure 7:** Scatterplot of cell type proportion between gene expression and chromatin accessibility data. Y=X line plotted

Supplementary figure 8

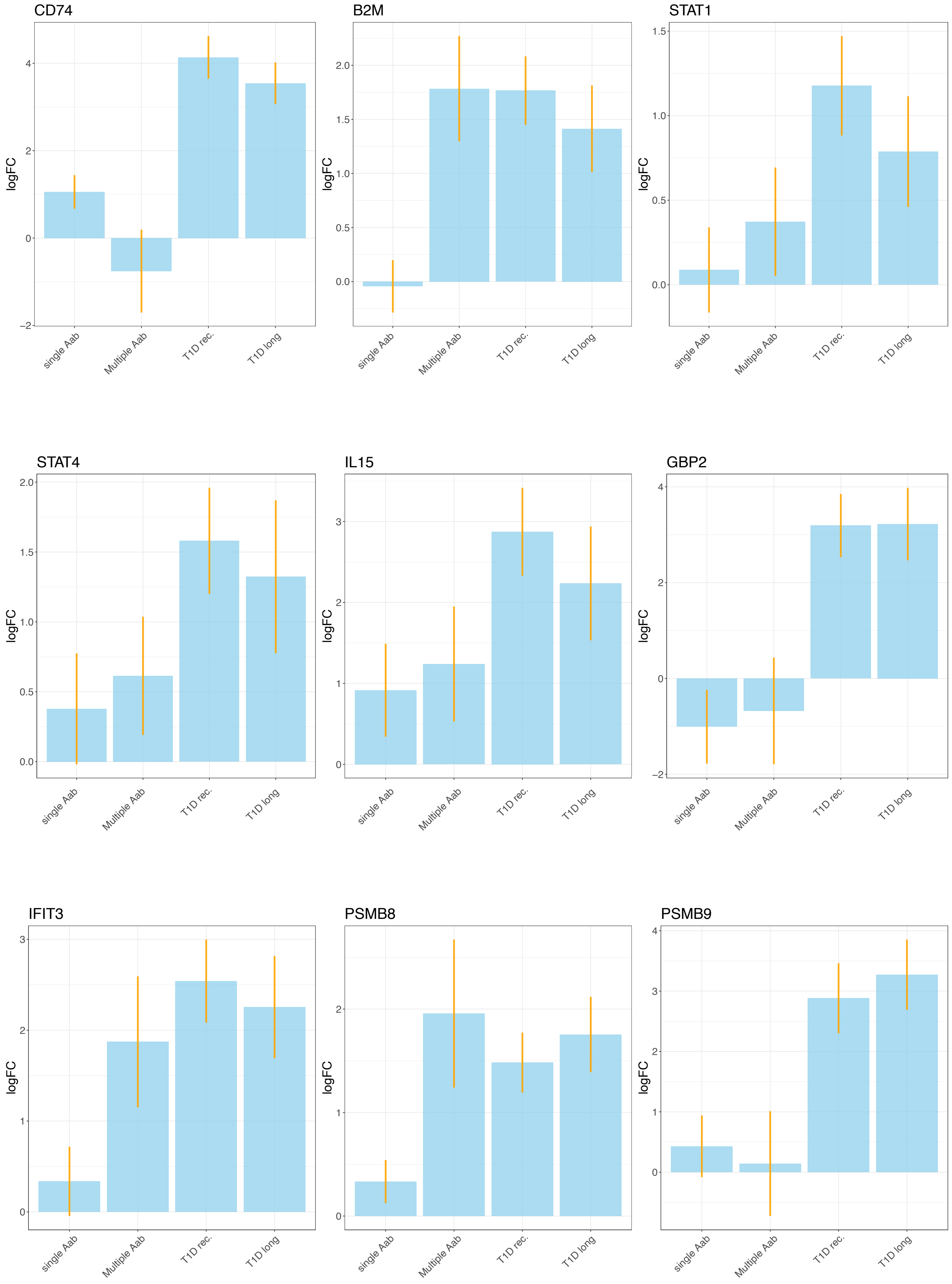


**Supplemental Figure 8:** Barplot of log2fc for each diabetes status relative to non-diabetic for a subset of differential expressed genes in beta cells

Supplementary Figure 9

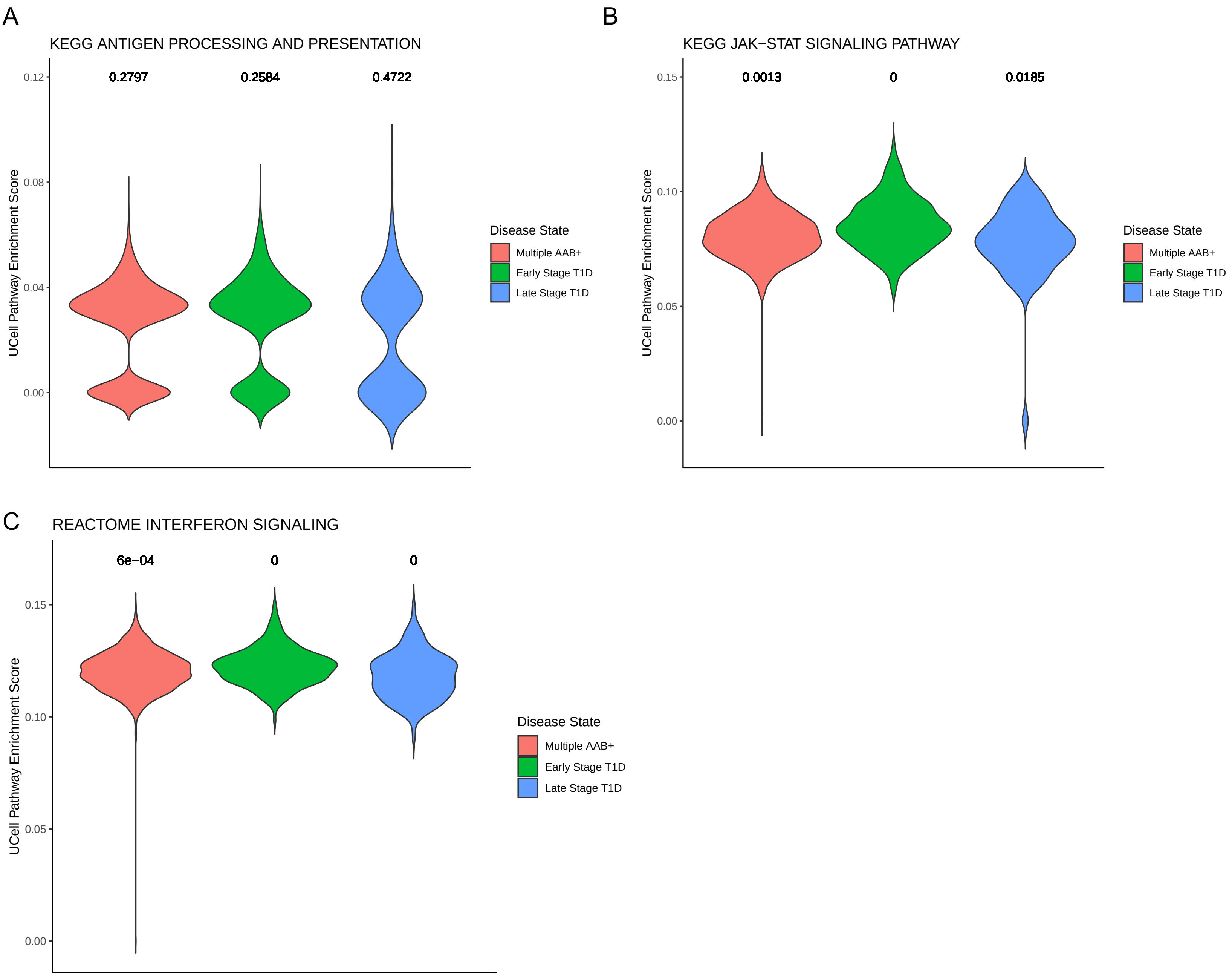


**Supplementary Figure 9**: Pathways up-regulated in recent onset T1D relative to non-diabetic controls exhibiting **A)** heterogenous pathway enrichment and **B-C)** a continuous distribution of UCell scores

Supplementary figure 10

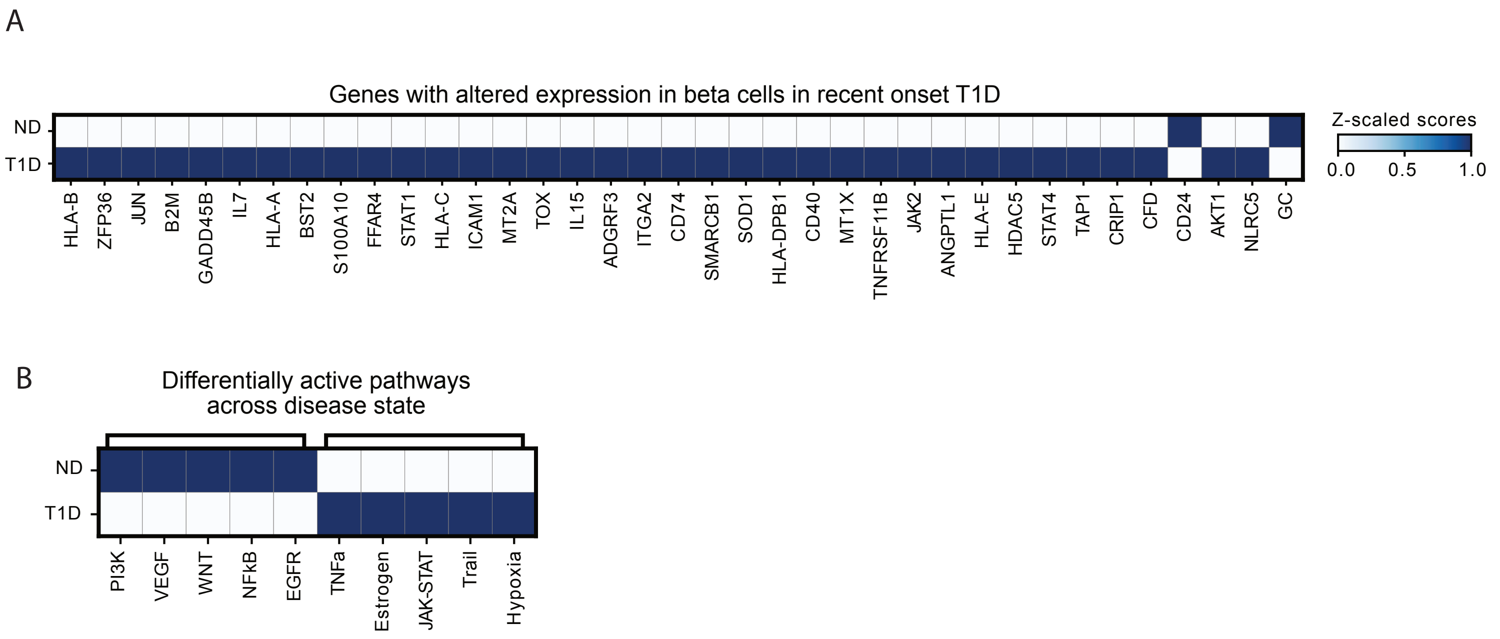


**Supplementary Figure 10**: A. Matrix plot showing scaled genes with altered expression in beta cells in recent onset T1D and their pattern on the spatial gene expression across conditions.  B. Matrix plot showing the scaled pathway activity of the differentially active pathways across disease state computed using progeny.

Supplementary figure 11

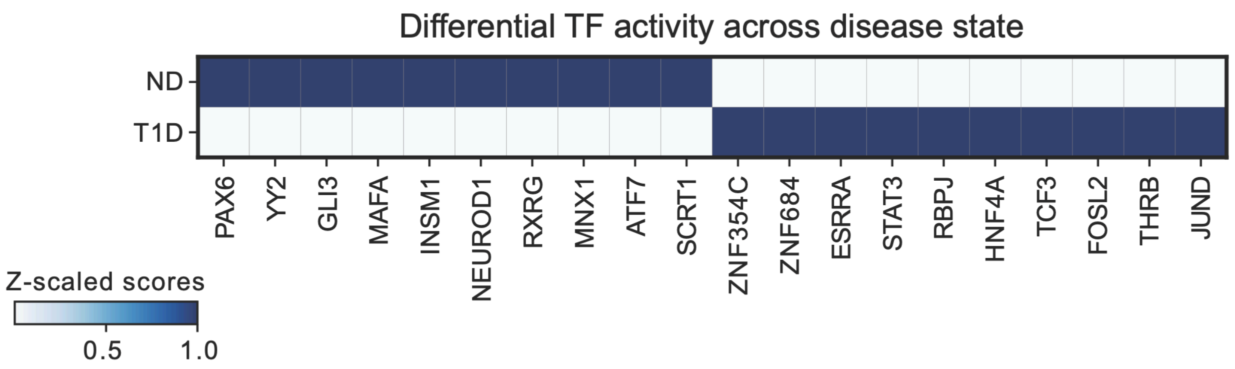


**Supplementary Figure 11**: Matrix plot showing the scaled TF activity of the differentially active TFs across disease state.

Supplementary figure 12

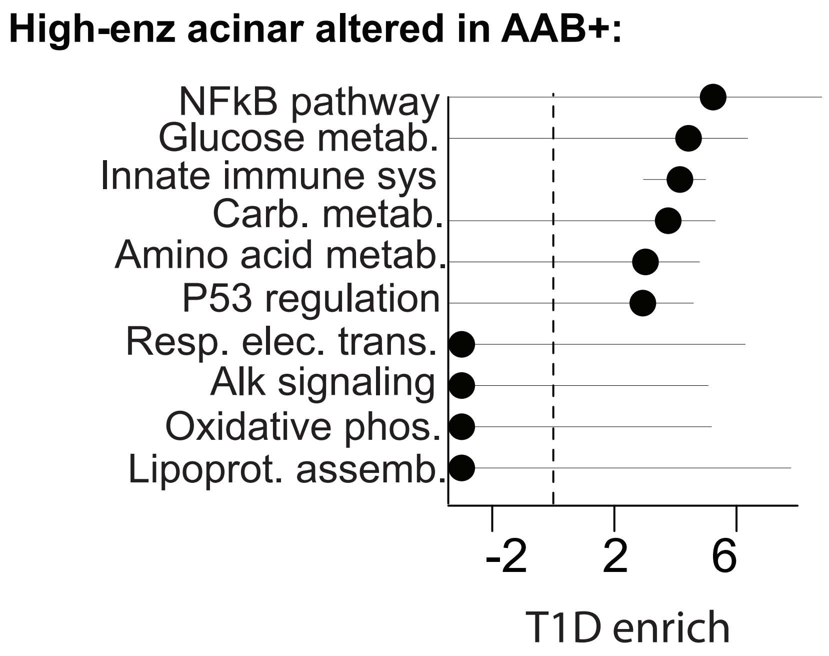


**Supplemental Figure 12:** Enrichment of T1D risk loci in pathways altered in T1D Aab+ donor in high enz. acinar population.

Supplementary figure 13

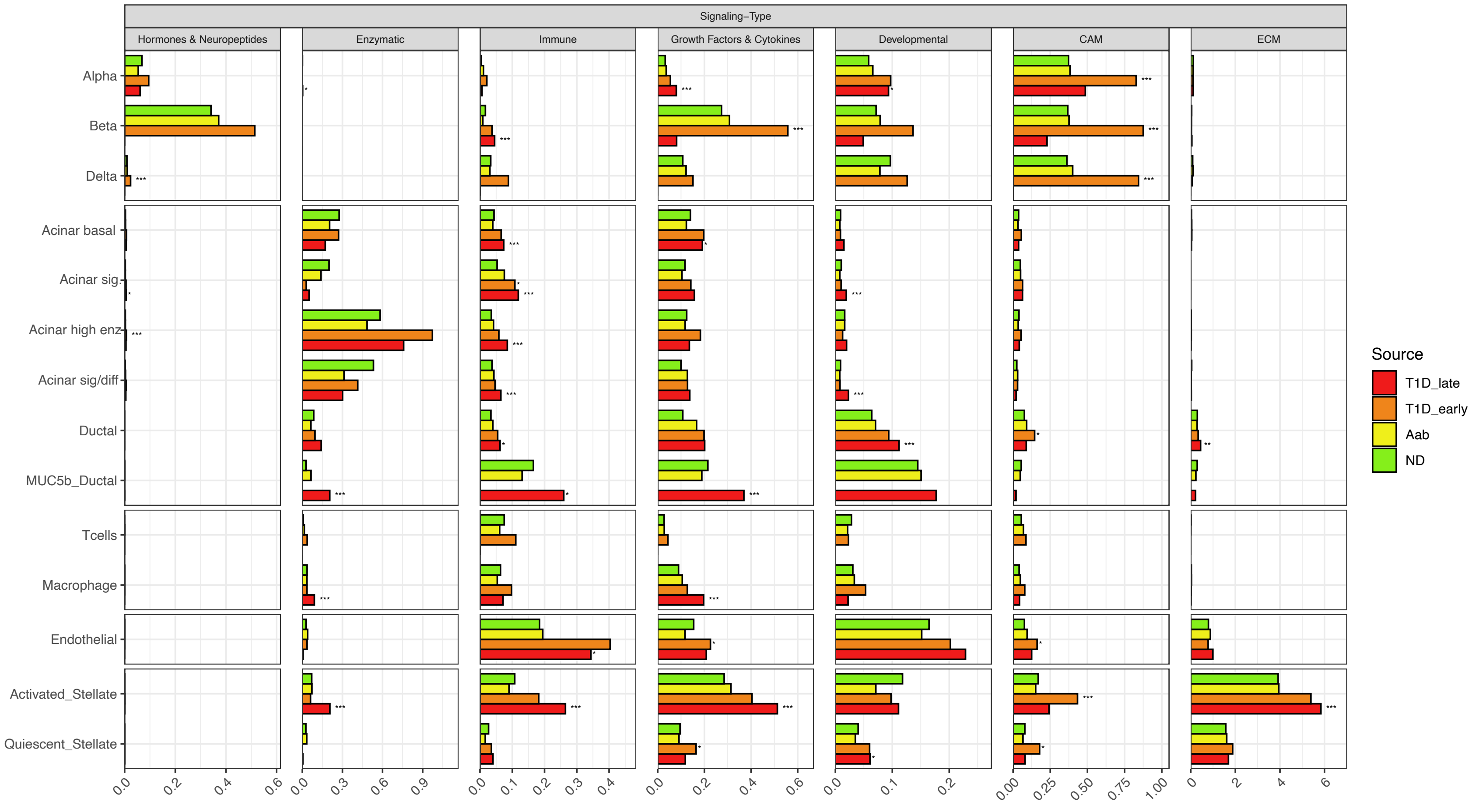


**Supplemental Figure 13:** Cell to cell signaling during diabetes broken down by signaling category per cell type and diabetes status.

Supplementary figure 14

A

B

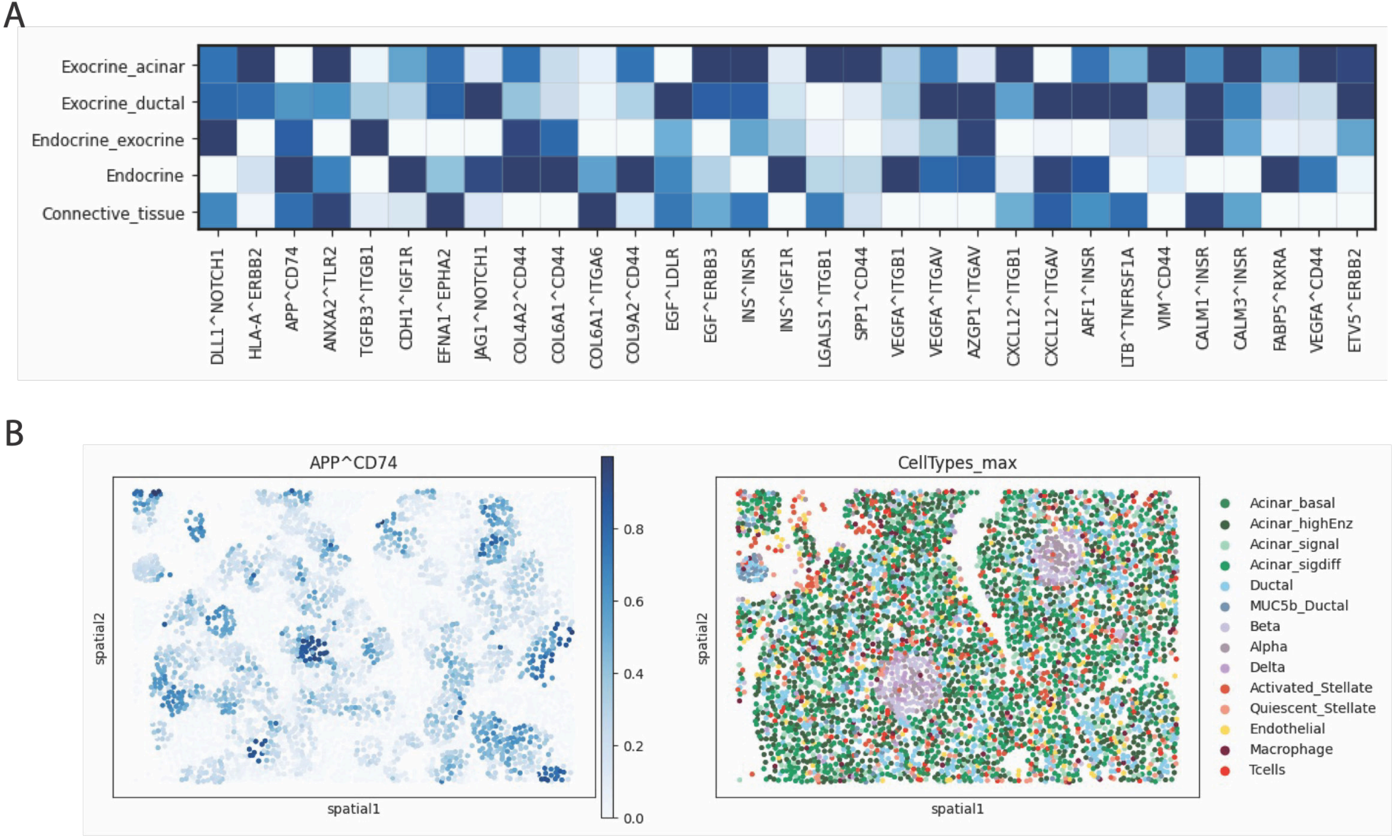


**Supplementary Figure 14**: A. Matrix plot showing ligand-receptor scaled score computed using Liana+ across spatial niches B. Spatial plot showing the APP^CD74 score in fov 10 of sile 1 (T1D representative fov).
